# Multiplexed measurements of protein-protein interactions and protein abundance across cellular conditions using Prod&PQ-seq

**DOI:** 10.64898/2026.01.01.697286

**Authors:** Tianyao Xu, Jingyao Wang, Yoonju Shin, Yuwei Cao, Lingzhi Zhang, Eduardo Modolo, Tamar Dishon, Jeremy Fisher, Matthew Norton, Christopher J Fry, Yossi Farjoun, Eric Mendenhall, Sven Heinz, Christopher Benner, Alon Goren

## Abstract

Methods to profile protein-protein interactions (PPIs) have limited scalability and can only study a handful of conditions and/or targets. Here, we introduce Prod&PQ-seq, a framework for multiplexed detection and quantification of PPIs and proteins. Our framework uses cross-linked cells, antibody-oligonucleotide conjugates (ab-oligos), and captures PPIs by the DNA-caliper, a specialized oligonucleotide for bidirectional priming of proximal ab-oligos. We benchmarked Prod&PQ-seq using recombinant complexes, titrations and cell mixture experiments and show that our framework is quantitative, reproducible, sensitive and specific. Applying Prod&PQ-seq to study Polycomb Repressive Complex 2 (PRC2) shows that EZH2 inhibition and expression of the oncohistone H3.3K27M weakens both PRC2-H3K27me3 interactions and PPIs within PRC2. Further, H3.1K27M and H3.3K27M variants lead to distinct PPI profiles such as the intensity of H3K27ac-K27M or H3K27ac-EED. Together, Prod&PQ-seq enables detection of changes in PPI composition and intensity and protein quantification across biological conditions, small molecule inhibition and genetic perturbations.

**GRAPHICAL ABSTRACT:** 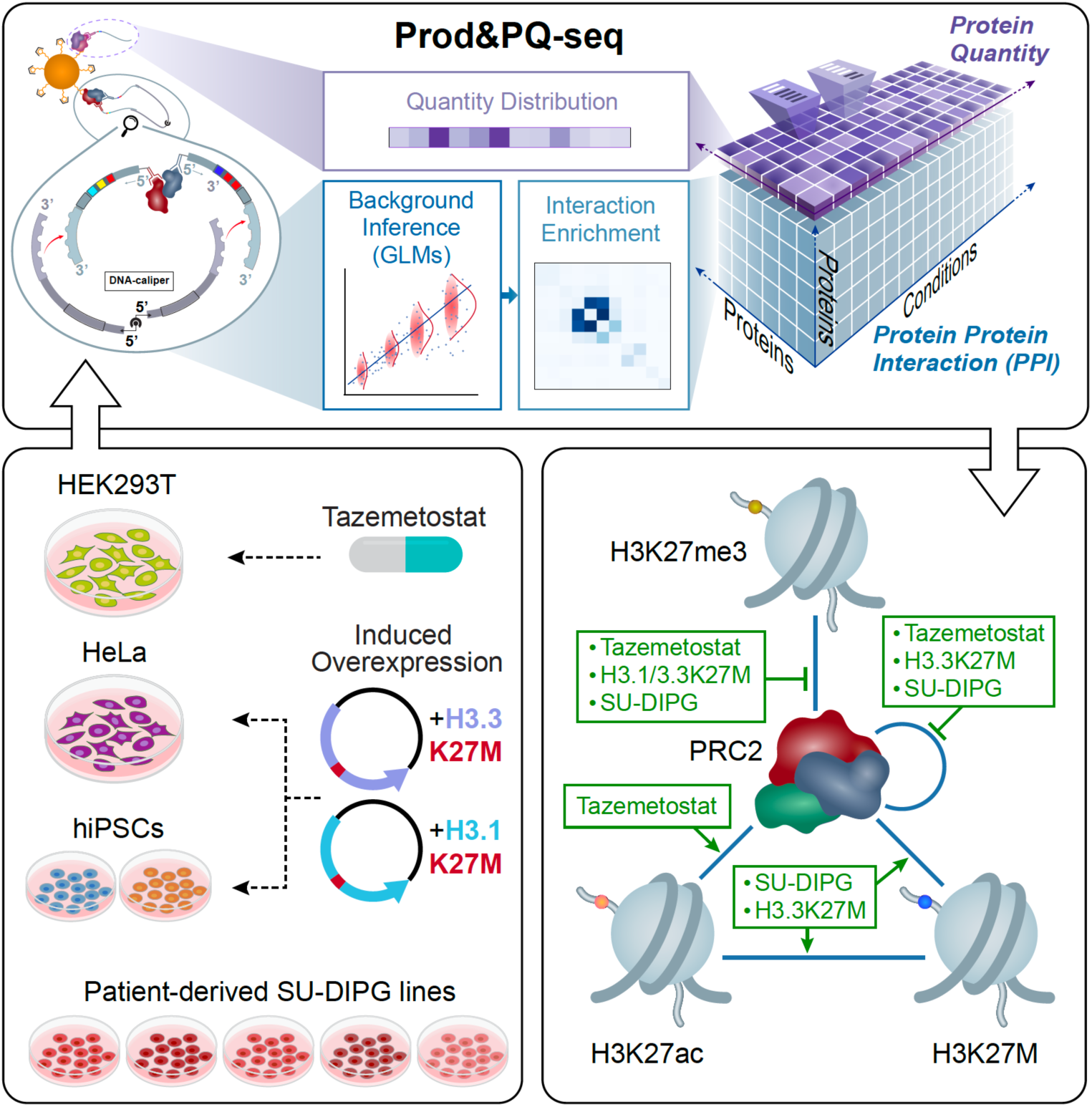

## INTRODUCTION

The assembly and interaction of cellular protein complexes as well as protein-abundance is key for biological processes such as metabolism, signal transduction networks, and regulation of chromatin structure^1–4^. The activity of such complexes could greatly depend on their modular composition and interactions with external proteins, which can highly vary among different biological conditions^5^. These observations highlight the need for reliable, cost-effective, multiplexed profiling of PPIs in cell contexts.

To date, multiple methods and tools have been developed to profile PPIs. Yeast-two-hybrid based methods target a set of proteins and examine their interactions by introducing transgenic fusion proteins in yeast cells^6^. Methods that build on mass spectrometry (e.g., AP-MS^7^, XL-MS^8^) focus on a handful of protein targets and study the proteome-wide interaction partners with the target proteins. Proximity-ligation assay (PLA) and other microscopy-based methods detect PPIs in cells and tissues through imaging technologies^9^. Other methods focus on computationally modeling and predicting PPIs using large scale PPI datasets to train the models^10^. These assays are complemented by large scale PPI datasets and databases (e.g., hu.MAP3.0^11^, BioGRID^12^).

However, the currently available tools and methods are mostly limited to the study of a small number of conditions, mainly due to the need to engineer a system for each target separately, or the requirement for specialized instrumentation (e.g., MS) incurring high costs. The limited ability to simultaneously detect the changes in PPIs across a variety of conditions is a major roadblock on many fronts in biological and medical research^13,14^.

A number of methods for high-throughput PPI detection methods were recently introduced, but these have key limitations. For instance, Prox-seq^15^ uses a proximity ligation assay to co-detect cell surface PPIs, protein, and mRNA levels at a single-cell level, but can only capture a subset of the PPIs as it requires matching oligo-barcoded antibodies and is limited to surface proteins. PROPER-seq^16^ converts cell transcriptomes into RNA-barcoded protein libraries before performing library-wise proximity ligation detection, but as an *in vitro* assay, it has limited power to detect changes in PPIs across diverse cellular conditions. TETRIS^17^ profiles higher-order protein interactions in parallel via bidirectionally ligating antibody-oligonucleotide conjugates bound to the epitopes, but as the sequencing libraries tend to be mostly composed of non-interacting products, conducting TETRIS on many samples could be challenging due to the high costs of the deep sequencing required for each library.

To address these limitations, we developed a novel framework that enables multiplexed detection of PPIs (proximity detection - Prod-seq) and measurements of protein quantification (PQ-seq) from a scalable number of biological conditions with an estimated cost per sample of $25 (or $41 when including in-house next-generation sequencing; **Supplementary Table 1**). The detection is achieved by converting such cellular measurements into next-generation sequencing libraries. Our framework uses cross-linked cell lysates and antibody-oligonucleotide conjugates (ab-oligos) for simultaneous detection and characterization of PPIs and protein-abundance (**Figure 1**). We built our framework to be accessible for a standard molecular biology laboratory and to allow generic, non-engineered cells and tissues as the starting material, avoiding the need for specialized instrumentation (e.g., mass spectrometry) or the creation of transgenic cell lines. Of note, given the relative ease of scaling up sample numbers, for each of the experiments we report here we included at least two biological replicates that were each profiled in two different Prod-seq&PQ-seq reactions (technical replicates) so that contributions of biological and assay-specific noise could be directly observed (**Supplementary Figure S1**). We accompany this manuscript with the optimized protocol vetted for accuracy and reproducibility, complete with reagent generation and purification guidelines (**Appendix**), as well as the computational analysis pipeline (**Code Availability**) to foster easy adoption of the method across a range of experimental systems.

**Figure 1.**
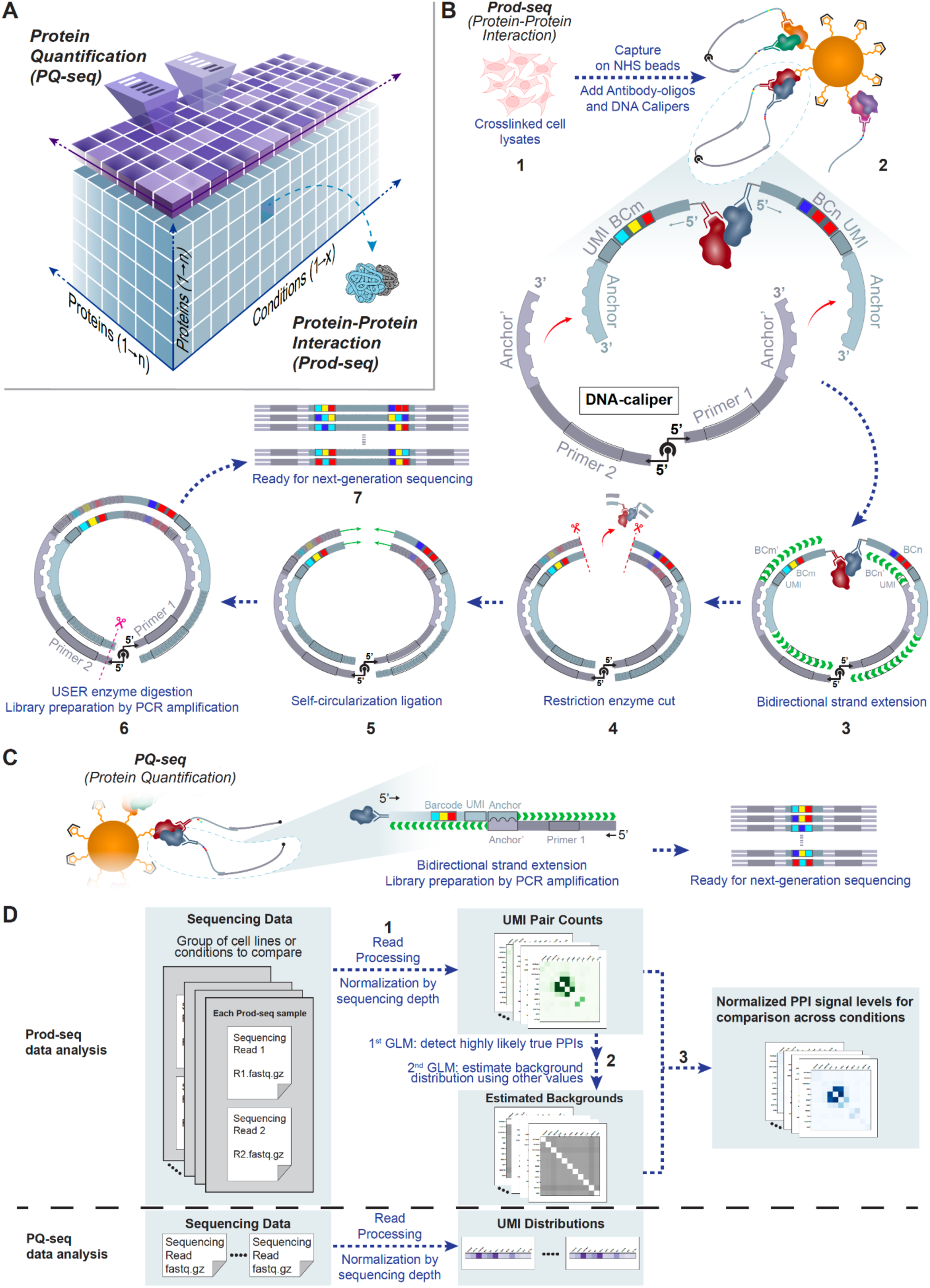
Prod&PQ-seq enables high-throughput detection of protein-protein interactions (PPIs) and protein quantification across different cells and conditions. **(A)** Schematics of the Prod&PQ-seq readout: PPIs detection (blue) and protein-abundance (purple). Each Prod-seq sample simultaneously detects interactions for all possible pairwise protein combinations within a scalable protein target pool for one cellular condition; using a subset of the reagents, the same sample can also go through optional protein quantification (PQ-seq). Conducted in parallel on a variety of cells and conditions, Prod&PQ-seq profiles the changes in PPIs and protein-abundance from different cellular contexts. **(B)** Schematics of the Prod-seq experimental workflow for PPIs detection. **(1)** Cells are crosslinked and sheared to generate lysates. Proteins and protein complexes in the cell lysates are captured on NHS (N-hydroxy succinimide) magnetic beads. Antibody-oligonucleotides (ab-oligos) are added to the cell lysates. The ab-oligos are antibodies linked to an oligo that encompasses a barcode (BC), a unique molecular identifier (UMI) and a universal anchor site (“*Anchor”*) which is identical for all ab-oligos. **(2)** The DNA-caliper (a single-stranded DNA molecule with two free 3’ ends for bidirectional priming), is generated by conjugating a pair of single-stranded oligonucleotides at their 5’ ends by click chemistry (depicted as ball and socket). Each 3’ end of the DNA-caliper (grey) harbors an *Anchor’* which anneals to the *Anchor* of any ab-oligo (red arrows). **(3-6)** The subsequent steps generate double-stranded DNA libraries via bidirectional strand extension, restriction enzyme digestion, circularization via ligation, and thermolabile USER enzyme cut (see **Methods** for details). **(7)** The product is PCR amplified to generate a library for next-generation sequencing. **(C)** Schematics of the optional PQ-seq experimental workflow. Using a subset of the reagents, PQ-seq can generate protein quantification readouts from the same cell lysates used in Prod-seq. More specifically, in a parallel reaction, one arm of the DNA-caliper is used to anneal with the ab-oligos and generate next-generation sequencing libraries via bidirectional strand extension and PCR amplification. **(D)** Schematics of the Prod-seq sequencing analysis framework. For PPI detection readout, the analysis pipeline uses a two-round generalized linear model (GLM) to estimate general detection background for PPI enrichment calculations (see **Methods** for details): **(1)** Next-generation sequencing reads are processed to extract pairwise protein barcodes and UMIs associations. **(2)** A two-round GLM uses the counts of UMI pairs from the group of samples to infer the distribution of PPI detection background. **(3)** The inferred background levels of each PPI in each sample are used to normalize the result (termed as PPI enrichment values hereafter), followed by additional normalization using the enrichment values of the interacting control PPIs. For PQ-seq readout, the analysis pipeline processes the sequencing reads to extract barcode and UMI associations; then the UMI counts of each protein barcode is normalized by the counts of the positive control proteins to calculate the normalized protein quantities.

We initially benchmarked our framework using recombinant protein complexes, where the PPI and protein compositions have known ground truths, to optimize signal-to-noise ratios and detection efficiencies. To extend the evaluation of Prod&PQ-seq to cell samples, we mixed cell samples from different conditions in predefined ratios to generate samples with known ground truths. In these mixed cell samples, the composition of PPIs and protein-abundance are expected to follow the mixing ratios, providing key reference points to fine-tune the experimental and computational parameters of our framework. Using these samples, we demonstrate the ability of Prod&PQ-seq to reproducibly detect PPIs and protein-abundances with high sensitivity and specificity.

To showcase our framework, we focused on the well-studied Polycomb Repressive Complex 2 (PRC2) and generated a set of 12 ab-oligos that target this complex as well as key histone modifications including H3K27me3, H3K27ac, and H3K4me3. Using this set, we surveyed the impact of a variety of perturbations in a range of cell conditions in cancer cell lines (HeLa and HEK293T) and human induced pluripotent stem cells (hiPSCs). We studied the impact of EZH2 inhibition on the PPIs and protein-abundance targeted by our ab-oligos. Additionally, we profiled the change in PPIs and protein-abundance following the expression and washing out (2-4 time points) of the oncohistone H3K27M in cancer and hiPSC lines. Lastly, we used our ab-oligo set to study an array of patient-derived Glioma cell lines harboring K27M mutations on histone H3.1 or H3.3.

Our major observations include: (1) When expressed, H3.3K27M has stronger colocalization with H3K27ac and PRC2 compared to H3.1K27M; (2) in patient-derived Glioma cell lines, H3.1K27M and H3.3K27M both result in reduced PPIs within core PRC2 members and with H3K27me3, and there is variability across patient-derived cell lines in the colocalization of H3K27M with H3K27ac and with PRC2; (3) Treatment with EZH2 inhibitor Tazemetostat results in reduced interaction between PRC2 members.

Altogether, we demonstrate the ability of Prod&PQ-seq to efficiently query the modularity of protein complexes as well as abundance changes across a range of biological conditions and time points – from small molecule treatment to genetic perturbation. The computational analysis pipeline uses estimation and normalization of PPI enrichment to allow comparison across samples, conditions, and perturbations, complimenting the reduced requirements of specialized instrumentations and cell engineering to enable the scaling of comparative studies measuring PPIs and protein-abundance.

## RESULTS

### Multiplexed quantification of PPIs and protein-abundance in cellular context

The Prod&PQ-seq framework uses cross-linked cell lysates and antibody-oligonucleotide conjugates (ab-oligos) for multiplexed detection and quantification of PPIs and protein-abundance. The ab-oligos contain barcode sequences corresponding to the epitope, unique molecular identifiers (UMIs^18^), and a universal 3’ end “*Anchor*” sequence for in-parallel experimental processing of all barcode pairwise proximities simultaneously (**Figure 1**, **Supplementary Note S1**).

Efficient profiling of PPIs via detection of proximal nucleic acids requires capturing both on a single molecule, which is challenging as the extension of oligonucleotides must be initiated in a 5’ to 3’ direction. Thus, commonly used reagents are incompatible. To overcome this challenge, we invented the DNA-caliper: a specialized single-stranded oligonucleotide with two free 3’ ends that allows bidirectional priming and thus serves as a detector of molecular proximity by converting biological information into easily read DNA sequences (**Supplementary Note S1**). The DNA-caliper is generated by covalently conjugating the 5’ ends of two single-stranded DNA arms using click chemistry^19^. The two 3’ arms of the DNA-caliper can anneal to the ab-oligos via the universal anchor sequences, thus enabling bidirectional strand extension. The extended DNA-caliper arms are then converted to sequencing libraries through additional steps, including designated linearizations, ligation, and PCR amplification (**Figure 1B**). Using a subset of the reagents, protein-abundance is quantified in a parallel reaction (**Figure 1C**).

Different from Proximity Ligation Assay (PLA)^9^, the Prod-seq component of our framework detects PPI with specificity which is provided by hybridization of reverse-complement DNA molecules. In particular, the 3’ end of each single-stranded oligonucleotide of the ab-oligos includes 16 bases of designated “*Anchor*” region which is a reverse complement to the two 3’ ends of the DNA-caliper arms (**Figure 1B**). Thus, to avoid the DNA-caliper arms from binding both ends to the same ab-oligo, our framework requires ab-oligo reagents that are significantly enriched for a degrees of labeling of 1 (DOL=1), meaning one oligo is conjugated to each individual antibody molecule. Therefore, commonly used approaches for the conjugation and purification of ab-oligos (e.g., as used by CITE-seq^20^, Prox-seq^15^) were incompatible with our framework. To address this challenge, we established a process to obtain ab-oligos with the vast majority having one oligo conjugated to each individual antibody molecule (**Supplementary Note S2, Supplementary Figure S2-S3**). Our approach for the conjugation and purification process is general and can be applied to any monoclonal IgG antibody.

### Prod&PQ-seq enables reproducible, sensitive and specific quantification of PPIs and protein-abundance in a range of cellular conditions

We benchmarked Prod&PQ-seq in a stepwise manner. Initially, all reagents and enzymatic steps in our framework were individually optimized. These optimizations included evaluation of different designs of the oligonucleotide sequences for the ab-oligos and DNA-caliper (e.g., accounting for sequence similarity), enzymatic reaction conditions (for increased efficiency), generation and purification methods for the DNA-calipers, as well as strategy for enrichment of single oligo labeled antibodies (**Supplementary Note S2-3, Supplementary Figure S2-S4**). We additionally evaluated the epitope specificity of multiple antibodies using either CRISPR interference (CRISPRi), or induced expression or degradation conditions followed by immunoblots (**Supplementary Figure S5**). Following these procedures, we focused on 12 epitopes and generated and purified a set of ab-oligos targeting them. This ab-oligo set is focused on PRC2 (EZH2, SUZ12 and EED) and related interactors (AEBP2 which is part of PRC2.2^21,22^), key histone modifications (e.g., H3K27me3 and H3K27ac) as well as interacting positive controls PPIs (MED12 and CycC^23^), and non-interacting negative controls (HA-tag, EGFR) (**Supplementary Table 2**).

Next, using our set of ab-oligos we evaluated and further optimized the performance of the Prod&PQ-seq protocol using recombinant protein complexes which served as a known ground truth for protein-abundance and PPI distribution. We focused on two recombinant complexes that were targeted by our ab-oligo set: (1) mononucleosomes with both H3K4me3 and H3K27ac modifications on each histone tail, and (2) a recombinant PRC2 complex containing EZH2, SUZ12, EED and RbAp46/48 (**Methods**). Using this recombinant system, we evaluated the level of non-specific detection (e.g., signal from targets that are not present), signal-to-noise ratios and reproducibility between replicates. This system also allowed us to establish an optimized protocol. Employing this optimized Prod&PQ-seq protocol, we profiled the recombinant complexes and observed a high signal-to-noise level (about 98-99% of UMI pairs associated with expected PPIs and about 99% of UMIs associated with expected proteins for Prod-seq and PQ-seq respectively). In addition, the distribution of UMI pairs is highly consistent among technical replicates (**Supplementary Figure S6**). As our protocol captures protein complexes on magnetic beads, protein complexes randomly colocalize on bead surfaces and could result in PPI detection noise. To evaluate the level of this noise, we mixed the recombinant PRC2 and mononucleosomes prior to immobilizing them on the protein-capture magnetic beads. We observed that the noise from this source is low and of similar magnitude to unspecific PPI detection (0-1.5% of reads, **Supplementary Figure S7**).

While the recombinant complexes provided a convenient system to optimize our framework, this system does not capture the inherent noise and broad dynamic range of PPIs and protein-abundances in cellular conditions. Thus, we used Prod&PQ-seq data generated from fixed cells to develop a designated computational pipeline for processing and analyzing sequencing readouts (**Figure 1D**). This pipeline first processes the reads to remove UMI duplicates and then extracts barcode pairwise-associations (defined as interaction counts for Prod-seq) and individual barcode-counts (PQ-seq). The pipeline then estimates PPI-detection background via a two-round generalized linear model (GLM) similar to the approach used by HiC-DC^24^ to estimate background. In the two-round GLM, the background level of each PPI in each sample is assumed to follow a beta-binomial distribution. The mean of the beta-binomial distribution of each epitope-pair is a linear combination of the interaction counts of the negative control (HA-Tag; which is absent in the samples used in this study) with each of the epitopes in that pair plus an intercept term. The two-round GLM is applied collectively to all beta-binomial distributions from a set of Prod-seq samples to enable comparison of conditions using maximum likelihood estimation. The first GLM excludes the probable true PPIs to increase power to estimate background, and the second GLM estimates the background distribution. We then calculate a PPI enrichment value for each epitope-pair in each sample as the ratio of the interaction count to its estimated background level. A detailed description of the computational analysis framework is provided in **Supplementary Note S4**.

We then assessed the ability of our joint experimental-computational framework to quantitatively detect changes in PPI and protein compositions in cellular contexts. To generate samples with known variations within the magnitude of changes in PPIs and protein abundances, we mixed the lysates from HEK293T cells and a patient-derived Glioma line (SU-DIPG36 described below) at various ratios (**Figure 2A**). We then used these mixture samples to calibrate our framework, where PPI enrichment and protein abundances are expected to follow the mixture ratios linearly (assessed by linear regression hereafter). Using this approach, we show that our framework can capture PPI enrichment and protein-abundance with high linearity (R^2^ > 0.6). As expected, for PPIs and proteins with minimal difference (absolute value of slope coefficients < 0.1) between the conditions, the random detection-noise dominated, resulting in lower linearity (R^2^ < 0.6; **Figure 2B, C, E, Supplementary Figures S8-S9**).

**Figure 2.**
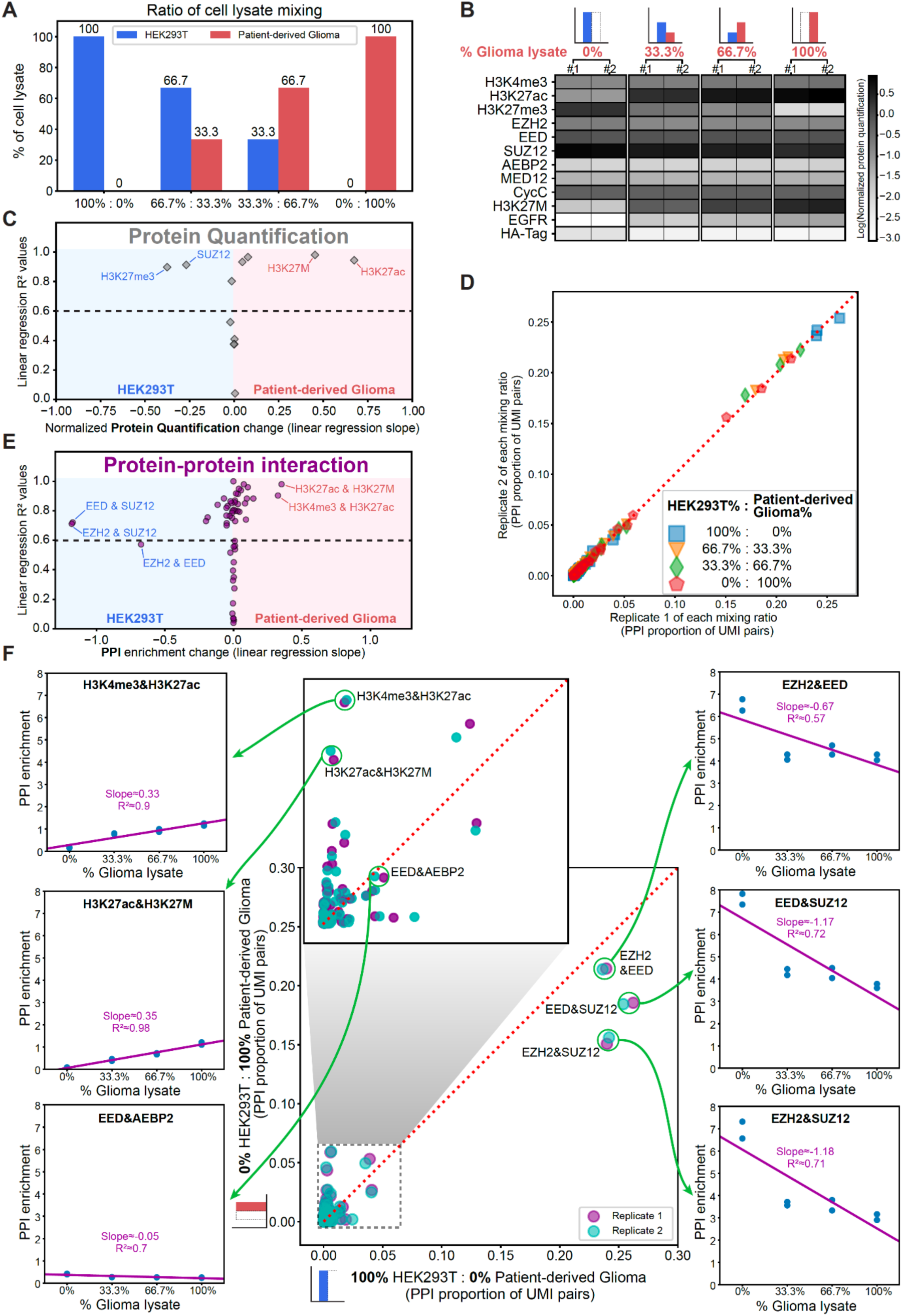
Prod&PQ-seq accurately detect quantitative changes in PPIs and abundances. **(A)** HEK293T (blue) and patient-derived Glioma (red) cell lysates are mixed at different ratios to mimic lysates with a change in PPI compositions. Each lysate mixing condition was performed with two technical replicates. **(B)** PQ-seq detects the change in protein-abundance from mixed cell lysate samples. PQ-seq readouts from each of the cell lysate mixing ratios are displayed on the heatmap as normalized and log transformed protein quantification (two replicates; normalized by the positive control UMI counts, **Methods**). **(C)** Protein quantifications change linearly with respect to the compositions of cell lysate mixtures. For each protein, a linear regression is performed using all replicates from all mixing ratios (2 replicates for each of the 4 mixing ratios). Linearity of quantification is measured by R^2^ value from each linear regression fit (plotted on the vertical axis); the magnitude of quantification change is measured by the slope from each linear regression fit (plotted on the horizontal axis). Dashed line=R^2^ value cutoff of 0.6. **(D)** PPI detection profiles are reproducible across a range of mixing conditions. Each point represents the UMI proportions of one PPI between two replicates of the different mixing ratios; the different colors and shapes represent the range of mixing ratios. **(E)** PPI enrichment values change linearly with respect to the compositions of cell lysate mixtures. A similar analysis to the linear regressions in C was conducted using the PPI enrichment values instead of the normalized protein quantification values as dependent variables. Dashed line=R^2^ value cutoff of 0.6. **(F)** Accurate detection of the differential PPIs between two mixed cell lines. Center: scatter plot of the UMI pair proportions of each PPI compared between HEK293T lysate (horizontal axis) and Glioma lysate (vertical axis). The two technical replicates are plotted separately (purple and cyan). Left and right: scatter plots of PPI enrichments of the five PPIs with maximal change in enrichment values between the two cell lines (EZH2&EED, EED&SUZ12, EZH2&SUZ12, H3K4me3&H3K27ac, H3K27ac&H3K27M) and one PPI with minimal change (EED&AEBP2) in the 4 mixing ratios, and their corresponding linear regression fit lines.

Moreover, the PPI pairs and protein abundances with the largest changes in magnitude (defined according to the slope of the linear regression) are the same ones that differ the most in the raw Prod&PQ-seq readout prior to application of our model (**Figure 2F, Supplementary Figures S8-S9**). For instance, the PPIs between H3K27ac and H3K27M showed the most positive linear regression slope (0.35; **Figure 2F**) with increasing percentage of Glioma lysate in the mixtures. In the raw Prod-seq readout, this proximity also showed a strong enrichment in the Glioma line compared to HEK293T (4.4% vs 0.3% of interaction counts respectively, **Supplementary Figure S9**), further corroborating our computational approach. Altogether, these results provide support to the ability of our framework to quantitatively detect a broad dynamic range in changes in PPI and protein abundances between different cell samples.

### Prod&PQ-seq detects the interaction and abundance changes of PRC2 upon drug treatment and cellular perturbations

Leveraging the calibrated version of our framework to quantify PPIs and protein abundances, we next studied the impact of drug treatment and genetic perturbations centering on PRC2 in different cell contexts. We focused on Tazemetostat, an FDA-approved EZH2 inhibitor^25^ that blocks the enzymatic function of EZH2 in writing the H3K27me3 histone modification^26^. As a competitive inhibitor, it is expected to reduce the interaction between PRC2 and H3K27me3. We treated HEK293T cells for 48 hours with Tazemetostat (or DMSO as a control) and performed Prod&PQ-seq. Following treatment, we observed the expected decrease in the level of H3K27me3 by PQ-seq, and confirmed a similar decrease by immunoblots (**Figure 3A-C**). We also detected a reduction in the interaction between H3K27me3 and core PRC2 members EED and SUZ12 (**Figure 3C-E, Supplementary Figures S10-11:** p-value < 0.05 for each PPI; **Methods**), aligning with the mechanism of action of Tazemetostat. Additionally, our PQ-seq analysis shows that Tazemetostat treatment leads to an increase in the levels of H3K27ac (**Figure 3C**), in line with observations from previous studies conducted in other cell lines^27^.

**Figure 3.**
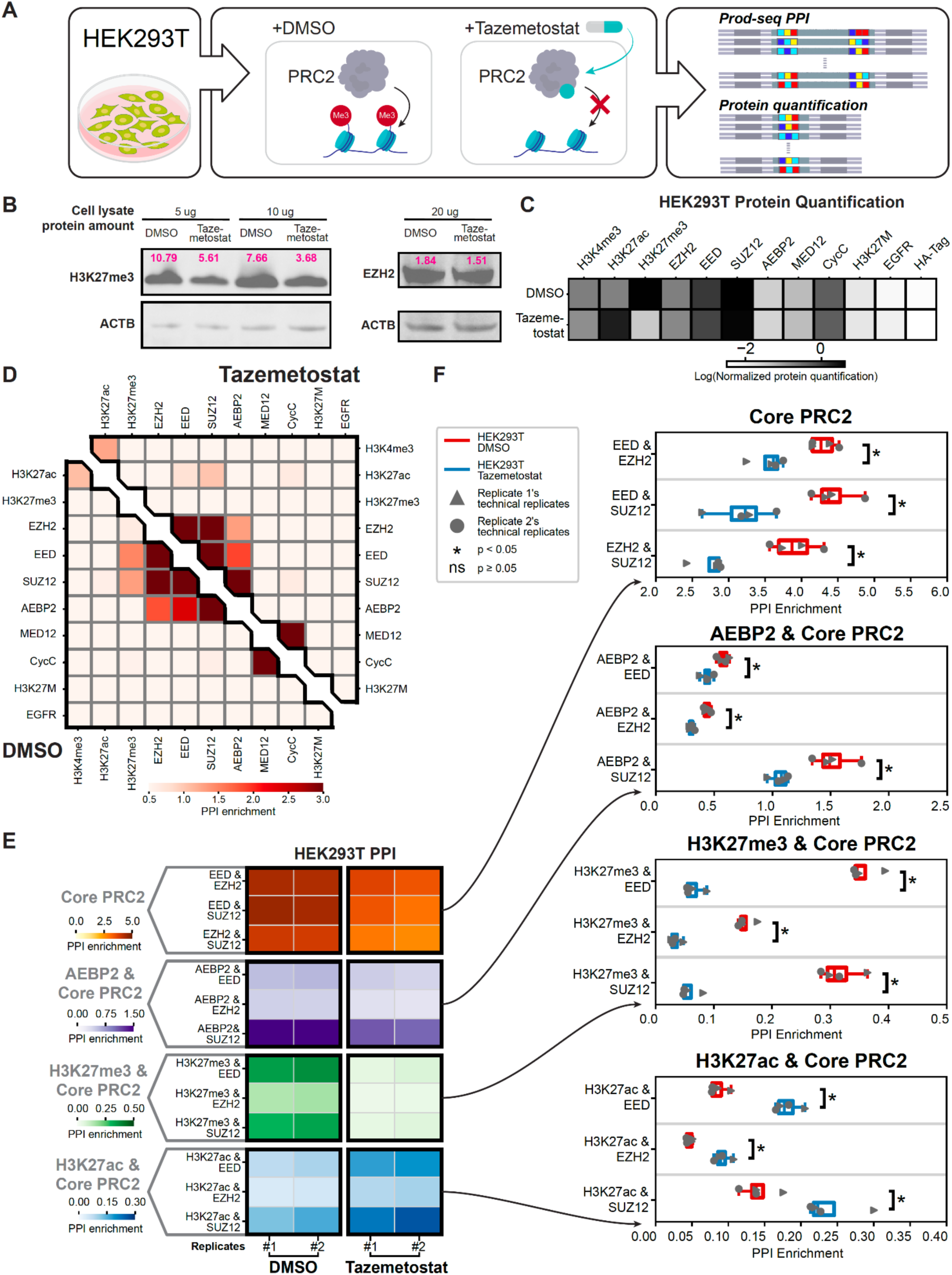
Prod&PQ-seq detects reduction in PRC2-H3K27me3 interaction upon Tazemetostat treatment. **(A)** Tazmetostat/DMSO treatment procedure. HEK293T cells are treated with Tazemetostat (10 μM) or DMSO (control) for 48 hours and then harvested for Prod&PQ-seq experiments (see **Methods** for details). **(B)** Immunoblot confirms the decrease in H3K27me3 upon Tazemetostat treatment. Left: immunoblot images of the H3K27me3 and Beta actin loading control (ACTB) upon DMSO or Tazemetostat treatment with different amounts of cell lysate proteins loaded. Right: immunoblot images of the EZH2 and Beta actin loading control (ACTB) upon DMSO or Tazemetostat treatment. Magenta: intensity of the target band normalized to control analyzed using ImageJ (**Methods**). **(C)** PQ-seq detects the decrease in H3K27me3 upon Tazemetostat treatment. PQ-seq readouts are averaged across the replicates for each condition and displayed on the heatmap. **(D)** Prod-seq detects the assembly of PRC2 and the change in its interaction with H3K27me3 following Tazemetostat treatment. The PPI enrichment values are calculated using the Prod-seq data analysis pipeline (**Methods**), and enrichment values from the PPIs are averaged across replicates to generate the heatmaps. Lower-left: HEK293T cells treated with DMSO; upper-right: HEK293T cells treated with Tazemetostat. **(E)** The change in interactions captured by Prod-seq is reproduced between replicates. For each treatment condition (left: DMSO and right: Tazemetostat), two biological replicates are performed (#1, #2) each with two technical replicates. **(F)** The interactions between core PRC2 members and H3K27me3 are reduced upon Tazemetostat treatment. PPI enrichment values from (E) are plotted both as box plots (replicate 1: triangle, replicate 2: circle). Statistical significance are categorized using nominal p-values from two-sided Mann–Whitney U test using all technical replicates of all biological replicates.

Prod-seq analysis shows that upon Tazemetostat treatment there is a decrease in H3K27me3 levels and a significant decrease in the interactions between the core PRC2 members (p-value < 0.05 for each PPI; **Figure 3**). Thus, while prior reports showed a decrease in PRC2 bound to chromatin after K27me3 depletion^26^, we now observe that the interactions within the PRC2 complex itself are decreased despite individual components still being expressed at similar levels (**Figure 3D-F**).

Surprisingly, we also identify an increase in the PPI between H3K27ac and all three core PRC2 members (p-value < 0.05 for each PPI) upon Tazemetostat treatment (**Figure 3D-F**). This could be due to PRC2 recruitment being maintained at a subset of loci that gain H3K27ac in the absence of K27me3, or aberrant recruitment of PRC2 to additional open chromatin loci that contain H3K327ac.

To further showcase the capability of our framework to detect and quantify changes in PPIs and protein-abundance following perturbations, we focused on a male human induced pluripotent stem cell (hiPSC) line (S02315; iPSCORE_1_13^28^). We transduced these hiPSCs with a lentivirus that harbours a doxycycline-inducible cassette encoding either the oncohistone H3.3K27M or the non-mutant H3.3WT control (**Methods**). Following 6 days of induced expression (**Figure 4A**), we conducted Prod&PQ-seq and observed a noticeable difference in the levels and interactions of H3.3K27M between the cells expressing WT or mutant H3.3. The increase in H3.3K27M expression detected by PQ-seq was also confirmed by immunoblots (**Figure 4B, Supplementary Figure S11**). We observe that upon expression of H3.3K27M, this oncohistone primarily interacts with H3K27ac and core PRC2 (**Figure 4C-E, Supplementary Figure S12**). These observations are in line with a previous report conducted using patient derived Glioma cell lines^26^. We additionally observe that following the expression of H3K27M the interactions between core PRC2 members tend to be weaker, although only the reduction in PPIs between EED and SUZ12 is significant (p-value < 0.05).

**Figure 4.**
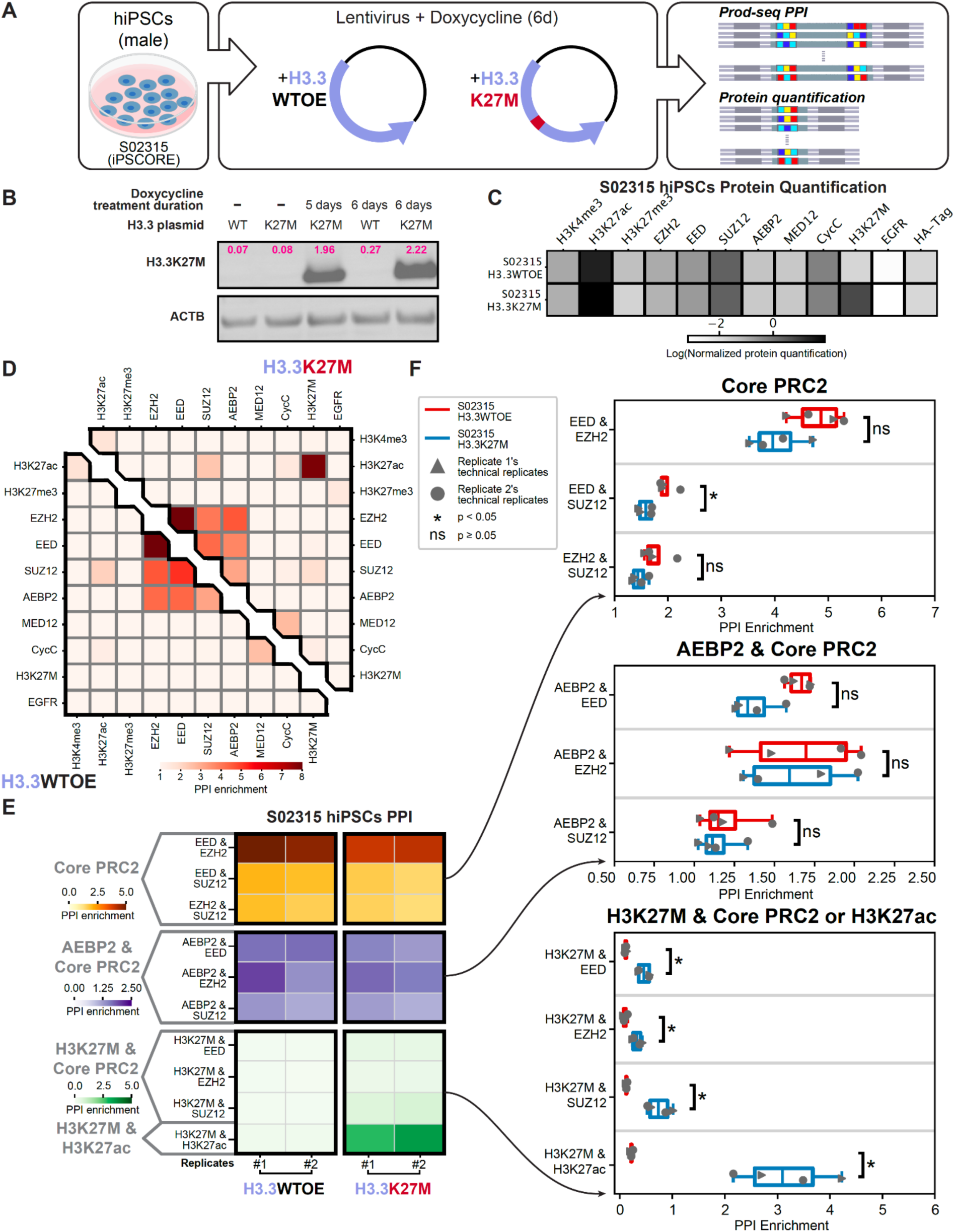
Prod&PQ-seq detects the impact of H3.3K27M on PPIs and protein abundance following expression of H3.3K27M in male hiPSCs. **(A)** Overview of the experimental design. Lentiviral vectors for inducible expression of H3.3WT (WTOE) or H3.3K27M are transduced into male hiPSCs iPSCORE_1_13^28^ (S02315) genomes (**Methods**). The cells are then treated with doxycycline for 6 days to induce the expression of H3.3WT or H3.3K27M. Cells are then harvested for Prod&PQ-seq. **(B)** Immunoblot confirms the expression of H3.3K27M. WT: doxycycline-inducible expression of H3.3WT; K27M: doxycycline-inducible expression of H3.3K27M. ACTB: Beta actin loading control. Magenta: intensity of the target band normalized to control analyzed using ImageJ (**Methods**). **(C)** Protein quantification subroutine detects the increase in H3K27M in the H3.3K27M expression condition. PQ-seq readouts (normalized by the positive control UMI counts, **Methods**) are averaged across the replicates for each condition and plotted in the heatmap. Colorbar–log normalized protein quantification. **(D)** The PPI enrichment values are calculated using the Prod-seq data analysis pipeline (**Methods**), and enrichment values of the PPIs are averaged across replicates to generate the heatmaps. Lower-left: induced expression of H3.3WT; upper-right: induced expression of H3.3K27M. **(E)** The change in H3.3K27M-H3K27ac interaction detected by Prod-seq is reproduced between replicates. For each treatment condition (left: induced H3.3WT expression and right: induced H3.3K27M expression), two biological replicates are performed (#1, #2) each with two technical replicates. The PPI enrichment values from Prod-seq are averaged between technical replicates to generate the heatmap. **(F)** Prod-seq detects the interaction of H3.3K27M with PRC2 and H3K27ac. The PPI enrichment values from Prod-seq (horizontal axes) of each technical replicate of the biological replicates (replicate 1: triangle, replicate 2: circle) are plotted both as scatter plots and box plots. Statistical significance are categorized using nominal p-values from two-sided Mann–Whitney U test using all technical replicates of all biological replicates.

### Detecting temporal changes in PPIs and protein abundance following removal of H3.3K27M

We developed Prod&PQ-seq to enable queries of PPIs and protein-abundance profiling across a wide range of samples (such as different cells) and conditions (such as different perturbations and/or time points). We first established cell lines with inducible expression of two oncohistone variants – H3.1K27M versus H3.3K27M – using lentiviral transduction into female adenocarcinoma cells (HeLa) followed by drug selection (**Figure 5A**). While the cassettes for the two oncohistones are identical, apart from the histone variant component, and we used the same selection procedure, we observed lower levels of H3.1K27M compared to H3.3K27M by PQ-seq (**Figure 5B**). This observation is in line with a previous report using these cassettes^29^. Although the expression levels of H3.1K27M and H3.3K27M differ, our computational model estimates interaction strengths in the context of the observed levels of the oncohistones, enabling us to compare their interactions across lines and conditions.

**Figure 5.**
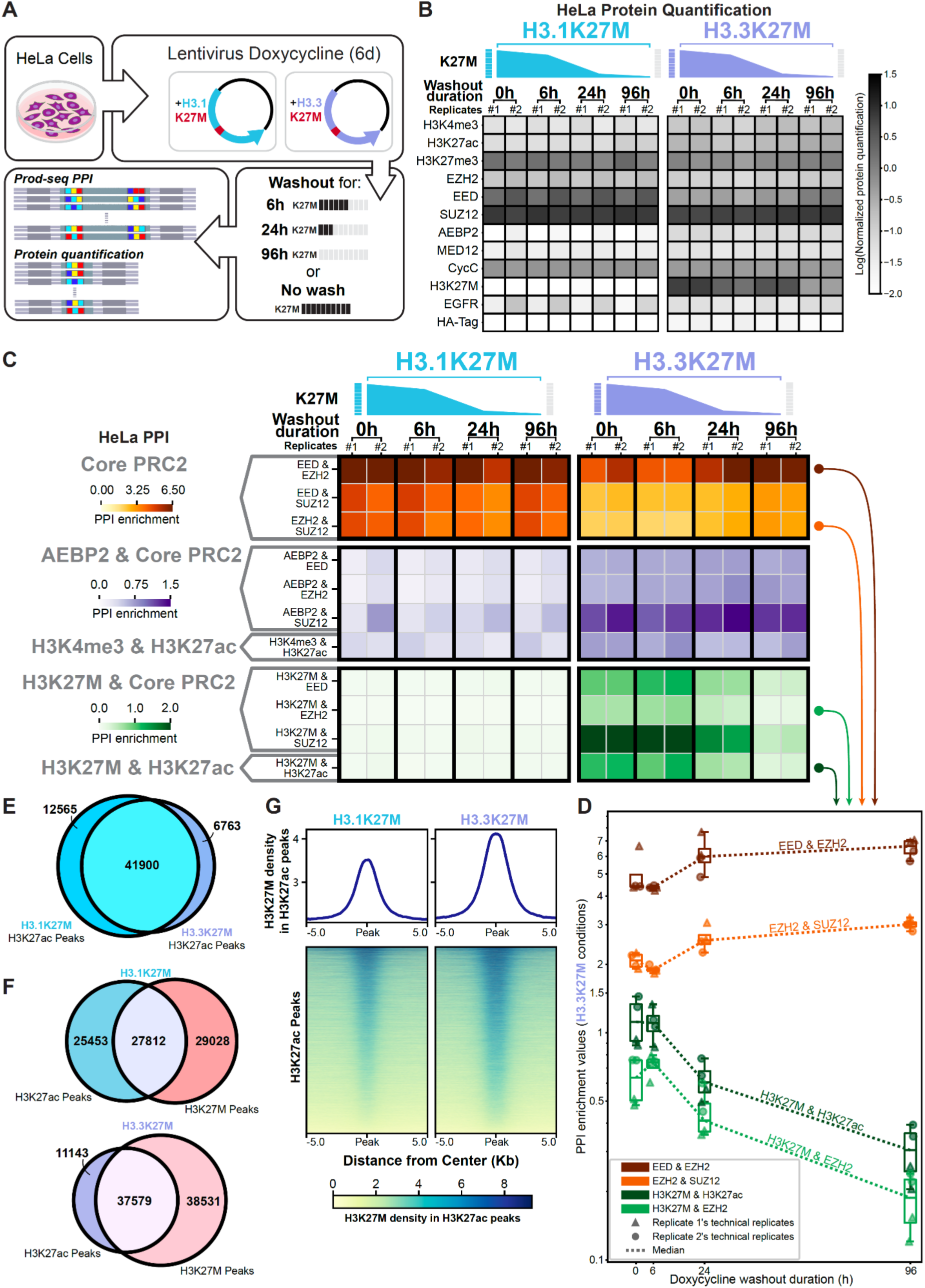
Prod-seq detects temporal changes in PPIs and protein abundance following removal of H3K27M oncohistones in HeLa cells. **(A)** Overview of the experiment design. Lentiviral vectors for inducible expression of H3.1K27M (cyan) or H3.3K27M (purple) are transduced into HeLa cells (**Methods**). The cells are then treated with doxycycline for 6 days to induce the expression of H3.1K27M or H3.3K27M. A subset of the cells from each condition are then immediately harvested, while the remaining cells went through 6 hours, 24 hours, or 96 hours of doxycycline removal (washout) and then harvested (**Methods**). **(B)** Protein quantification subroutine detects the temporal removal H3.3K27M. PQ-seq readouts (normalized by the positive control UMI counts, **Methods**) are averaged across the replicates for each condition and plotted in the heatmap. Colorbar–log normalized protein quantification. **(C)** Prod-seq detects the washout of H3.3K27M’s interaction with core PRC2 and H3K27ac. For each treatment condition (0h: doxycycline-induced expression; 6h, 24h, 96h: washout for 6h, 24h or 96h respectively), two biological replicates are performed (#1, #2) each with two technical replicates. The PPI enrichment values from Prod-seq are averaged between technical replicates to generate the heatmap. **(D)** Prod-seq detects a temporal trend of PPI enrichment. The PPI enrichment values (vertical axis) of the replicates from the different washout duration conditions (horizontal axis) are plotted as scatter plots and box plots. Dashed lines: trajectories of the median PPI enrichment values. **(E)** H3K27ac ChIP-seq peaks highly overlap between H3.1K27M and H3.3K27M expression. H3K27ac ChIP-seq is performed on H3.1K27M-expressing HeLa cells and H3.3K27M-expressing HeLa cells without washout of doxycycline. Numbers: number of ChIP-seq peaks for each category. **(F)** H3K27ac ChIP-seq peaks overlap more with H3.3K27M peaks than H3.1K27M peaks. H3K27M ChIP-seq is additionally performed on the same cell samples in (E). The numbers represent the counts of ChIP-seq peaks in each category. **(G)** H3.3K27M has higher coverage than H3.1K27M in H3K27ac peaks. H3.1K27M and H3.3K27M coverages are calculated on all H3K27ac peaks in H3.1K27M-expressed HeLa cells and H3.3K27M-expressed HeLa cells without washout of doxycycline (the union in (E)).

To demonstrate the ability of Prod&PQ-seq to survey a broad range of samples and conditions and to gain insight into the dynamic changes following the removal of H3K27M, we designed an experiment building on our doxycycline inducible cassette. We first induced the expression of H3K27M for 6 days, and then stopped the induction by removing doxycycline (washout). We then followed the changes in the PPIs and protein abundance for a total of 4 timepoints as oncohistone expression is slowly depleted (0h, 6h, 24h, and 96h; **Figure 5A**).

In our previous male hiPSCs experiments, the expression of H3.3K27M leads to weaker interactions within the core PRC2 members (EZH2, SUZ12, and EED; **Figure 4**). To evaluate whether the pattern extends to HeLa cells, we focused on these PPIs over the 4 timepoints. In a similar manner to the hiPSCs, H3.3K27M expression leads to weaker interactions within the core PRC2, which is gradually reverted following removal of the oncohistone variant (**Figure 5B-D, Supplementary Figure S13**).

When comparing the effect of expressing the two oncohistone variants, we observe that H3.3K27M exhibits stronger interaction with core PRC2 members as well as H3K27ac compared to the interaction of H3.1K27M with these proteins (**Figure 5C**). Previous studies using ChIP-seq demonstrated a genomic colocalization between H3.3K27M and H3K27ac^30^ and distinct H3K27ac patterns between diffuse intrinsic pontine Glioma (DIPG) cells expressing H3.1K27M or H3.3K27M^31^. Additionally, the interaction between H3.3K27M and H3K27ac gradually decreases along with the reduction in the levels of H3.3K27M (**Figure 5C, D**). Lastly, we observed that expression of H3.3K27M variant, compared to H3.1K27M, leads to higher PPI enrichments between core PRC2 members and AEBP2. These PPIs in cells that expressed H3.3K27M appear to be stable, and do not change with the reduction in the levels of this oncohistone variant (**Figure 5C**).

To provide support to the observations obtained by our novel framework, we conducted orthogonal experiments using the original cell lysates used for Prod&PQ-seq (**Figure 5E-G**). In addition to the aforementioned immunoblots showing concordant results to the PQ-seq readouts (**Figure 3B, 4B**), we focused on samples with maximal and minimal levels of H3K27M (0h and 96h washout timepoints, respectively) and performed ChIP-seq for H3K27ac and H3K27M using our automated protocol^32^. Of note, we do not expect that the results of the two methods to perfectly agree, since ChIP-seq is limited to a subset of the interactions as it only captures DNA-associated instances, while Prod-seq in theory, can detect all targeted PPIs.

Analysis of the ChIP-seq results showed that our H3K27M antibody was able to detect genomic localization of both H3.1K27M and H3.3K27M (**Figure 5G**). Further analysis of the data shows that generally, H3K27ac peaks had a high level of overlap between the two lines (68.4% of the union of the two sets of peaks; **Figure 5E**). However, when we compared the peak overlap of H3.3K27M and H3.1K27M, the H3.3 variant had higher overlap with H3K27ac peaks (43.1% vs 33.8%), in agreement with the higher PPIs of the H3.3K27M oncohistone with H3K27ac (**Figure 5C, F)** detected by Prod-seq. Further, H3.3K27M signal was higher than H3.1K27M signal in H3K27ac peaks (**Figure 5G**), in line with the Prod-seq result showing higher enrichment of PPIs between H3K27ac and H3.3K27M when compared to H3.1K27M (**Figure 5C**).

### Detection of different temporal interaction and protein-abundance patterns induced by the oncohistone variants H3.1K27M and H3.3K27M in hiPSCs

Next, to study if the impact of the two oncohistone variants is similar between cancer and stem-cell contexts, we established an inducible expression system in hiPSCs. We focused on the female hiPSC line S02316 (iPSCORE_1_14^28^) and transduced the same lentiviral constructs used in HeLa cells mentioned above to allow doxycycline inducible expression of H3.1K27M or H3.3K27M. We additionally conducted a similar oncohistone removal experiment; we first induced expression for 6 days using doxycycline (0hr), followed by 5 days of washout (120 hours; **Figure 6A**). The PQ-seq results and immunoblots, as well as the reduced interactions between H3.3K27M and core PRC2 members, confirm the depletion of the oncohistone variants following the washout (**Figure 6B, C, Supplementary Figure S14**). Similar to our observations in the HeLa cells, H3.1K27M and H3.3K27M exhibit different interaction patterns. While H3.1K27M shows minimal interactions with H3K27ac or PRC2 in these hiPSCs, H3.3K27M has strong interactions with both H3K27ac and PRC2 (**Figure 6D**), which we also observed in the male hiPSC line (S02315). Intriguingly, we detect a strong interaction between AEBP2 and SUZ12 induced by H3.1K27M expression phased out following the removal of this oncohistone variant, indicating an increased PRC2.2 complex formation in the presence of K27M in pluripotent cells but not in the HeLa cancer cell line (**Figure 5C**).

**Figure 6.**
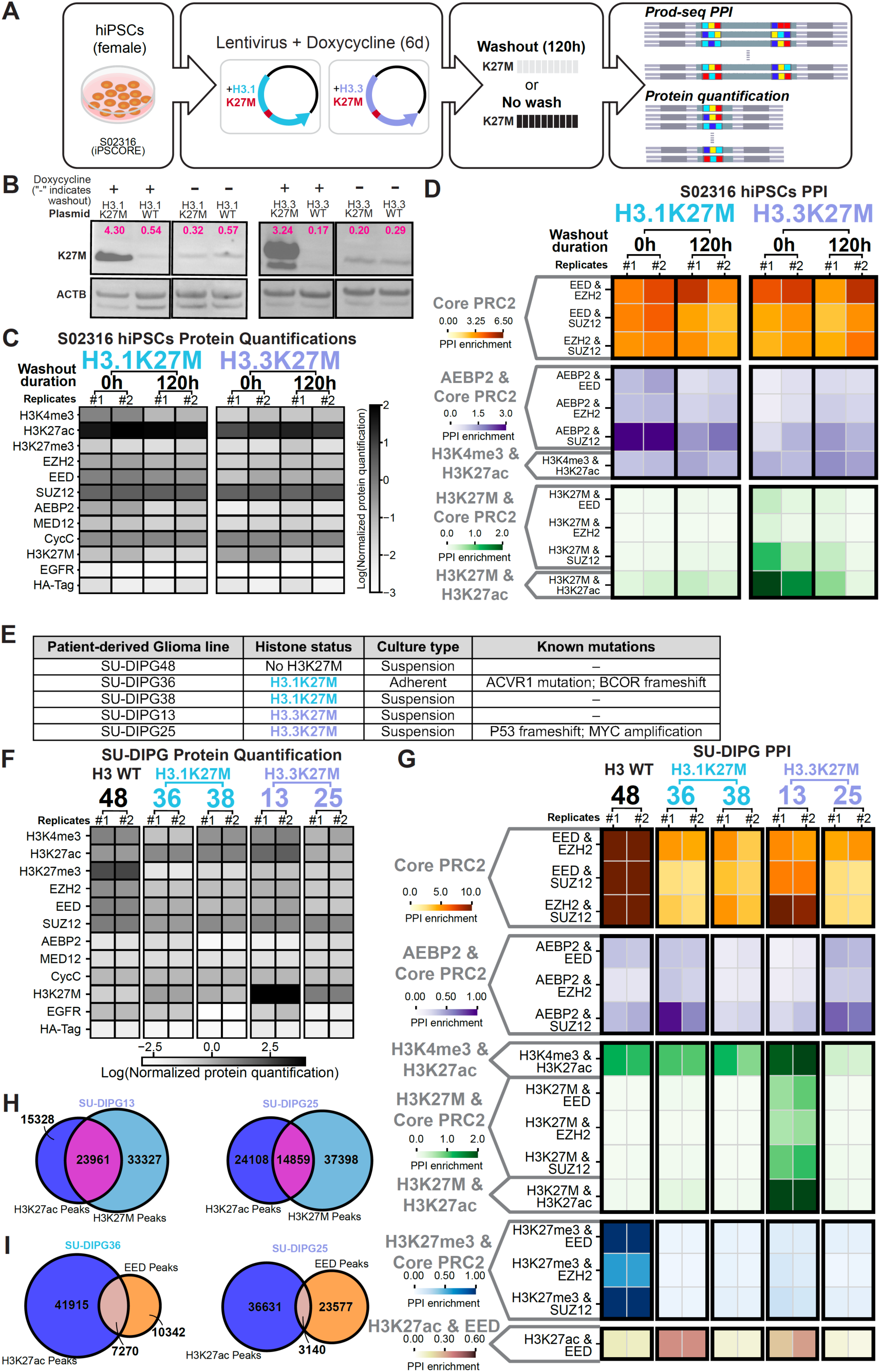
Prod-seq detects different interaction profiles for H3.1K27M and H3.3K27M in female hiPSCs (S02316) and detects different PPI and protein-abundance patterns in a range of patient-derived Glioma lines. **(A)** Overview of the experiment design. Lentiviral vectors for inducible expression of H3.1K27M (cyan) or H3.3K27M (purple) are transduced into female hiPSCs iPSCORE_1_14^28^ (S02316) (**Methods**). The cells are then treated with doxycycline for 6 days to induce the expression of H3.1K27M or H3.3K27M. Half of the cells from each condition are then immediately harvested, while the remaining half went through 5 days (120h) of doxycycline removal (washout) and then harvested (**Methods**). **(B)** Immunoblot confirms the induced expression and washout of H3.1K27M and H3.3K27M. Plus sign: doxycycline-induced expression; minus sign: doxycycline-induced expression followed by 5 days of washout for doxycycline. ACTB: Beta actin loading control. Magenta: intensity of the target band normalized to control analyzed using ImageJ (**Methods**). **(C)** Protein quantification subroutine detects the washout of H3.1K27M and H3.3K27M. PQ-seq readouts (normalized by the positive control UMI counts, **Methods**) are averaged across the replicates for each condition and plotted in the heatmap. Colorbar–log normalized protein quantification. **(D)** Prod-seq detects that H3.3K27M’s interaction with H3K27ac and PRC2 is stronger than those of H3.1K27M in both replicates. For each treatment condition (Dox: doxycycline-induced expression; wash: 5 days of washout after doxycycline treatment), two biological replicates are performed (#1, #2) each with two technical replicates. The PPI enrichment values from Prod-seq are average between technical replicates to generate the heatmap. **(E)** List of the SU-DIPG cell lines used and their histone mutation status. The cell lines include a H3WT line (DIPG48), two lines with H3.3K27M mutation (DIPG13 and DIPG25), and two lines with H3.1K27M mutation (DIPG36 and DIPG38). **(F)** Protein quantification subroutine detects distinct protein-abundance profiles for the five patient-derived SU-DIPG lines. PQ-seq readouts (normalized by the positive control UMI counts, **Methods**) are averaged across the replicates for each condition and plotted in the heatmap. Colorbar–log normalized protein quantification. **(G)** Prod-seq detects distinct PPI profiles for the five patient-derived SU-DIPG lines. For each SU-DIPG line, two biological replicates are performed (**Supplementary Figure 1**) each with two biological replicates. The PPI enrichment values from Prod-seq are average between technical replicates to generate the heatmap. **(H)** H3K27ac and H3K27M ChIP-seq peaks overlap more in SU-DIPG13 than in SU-DIPG25 (both cell lines harbour the H3.3K27M mutation). The numbers represent the counts of ChIP-seq peaks in each category. **(I)** H3K27ac and EED ChIP-seq peaks overlap more in SU-DIPG36 (H3.1K27M) than in SU-DIPG25 (H3.3K27M). The numbers represent the counts of ChIP-seq peaks in each category.

### Prod-seq reveals differences in PRC2 and H3.1/3.3K27M PPI patterns across a range of patient-derived Glioma cell lines

We next evaluated if our observations in cancer and pluripotent stem cell contexts can be extended to patient-derived Glioma lines where the oncohistone variants naturally arose. H3K27M has been reported to be a key oncogene in diffuse intrinsic pontine Glioma (DIPG)^33^. We conducted Prod&PQ-seq focusing on five patient-derived DIPG cell lines (SU-DIPG lines 13, 25, 36, 38 and 48; **Methods**). These include two lines harbouring H3.1K27M (SU-DIPG36 and 38), two harbouring H3.3K27M (SU-DIPG13 and 25), and one without the oncohistone (SU-DIPG48; **Figure 6E, Supplementary Figure S15**).

In agreement with widely reported observations in previous studies^34-36^, analysis of our Prod&PQ-seq data shows that the presence of H3.1K27M or H3.3K27M is associated with a noticeable decrease in the activity of PRC2 in catalyzing H3K27me3, reflected by a reduction in the levels of this histone mark (**Figure 6F**). Further, the PRC2 core members (EED and SUZ12) exhibit reduced interaction with H3K27me3 in the four lines harboring the K27M mutation when compared to the line without K27M (SU-DIPG48). Our PQ-seq results showed that SU-DIPG48 which has no oncohistone had the highest level of H3K27me3, and a line harbouring H3.1K27M had the lowest level of H3K27me3 (SU-DIPG36).

Consistent with results previously reported^37^, we find that SU-DIPG lines vary in their expression of K27M, with lines harboring H3.1K27M mutations showing slightly lower abundance of H3K27M than lines with H3.3K27M mutations, with line SU-DIPG13 exhibiting the highest abundance of H3K27M (**Figure 6F**). Looking at PPI interactions, we saw variation across the Glioma cell lines, possibly due to the impact of additional oncogenes on the biology of these cell lines or variation in cell line derivation from patient samples. We found the abundance of the core PRC2 members (EED, EZH2 and SUZ12) was reduced in all the lines compared to the line that had no H3K27M mutation. The Prod-seq results also showed that the PPIs between the PRC2 core members were weaker in all four H3K27M lines **(Figure 6F,G, Supplementary Figure S15)**. Similar to the changes observed in HEK293T cells treated with Tazemetostat, K27M expression led to the increased interaction of PRC2 components (e.g. EED) with H3K27ac in SU-DIPG36 and SU-DIPG13, but not the other lines. Of note, in SU-DIPG13 (harbouring H3.3K27M), EZH2&SUZ12 interaction was maintained at high levels. Our Prod-seq results also show that the PRC2.2 accessory protein AEBP2 presents strong interactions with SUZ12 in one H3.1K27M (SU-DIPG36) and one H3.3K27M lines (SU-DIPG25). Lastly, the interactions between H3K27M and H3K27ac ranged across the lines surveyed, with SU-DIPG25 showing the weakest PPI enrichment while SU-DIPG13 showing the strongest, despite both lines harboring H3.3K27M (**Figure 6E-G**).

To obtain support for these observations, we supplemented published ChIP-seq data^38^ from these DIPG lines with additional ChIP-seq samples we performed targeting EED, H3K27ac and H3K27M in a subset of the Glioma lines. Confirming the variability in interactions between K27M and H3K27ac observed in Glioma lines expressing H3.3K27M mutations, we detect a higher overlap of H3K27ac and H3K27M peaks in SU-DIPG13 (33.0% union of the two sets of peaks) compared to the SU-DIPG25 line (19.5%), which is in agreement with the Prod-seq result (**Figure 6G-H**). In addition, we observe that two lines with different H3K27M variants exhibited distinct degrees of genomic overlap between H3K27ac and EED in the SU-DIPG36 line (12.2%) vs. the SU-DIPG25 line (5.0%), reflecting the differences in H3K27ac and EED PPI enrichment obtained from these cells (**Figure 6G, I**).

Altogether, these results show that when comparing the different SU-DIPG cell lines, we observe notable interaction pattern variations between cell lines, even if they express the same oncohistone variant. These observations suggest that in addition to the specific H3K27M variant, other cell line-specific factors may come into play and impact the PPIs and protein abundance. These results could serve as the basis for follow-up studies interrogating the specific impact of the mutant variant within different cellular contexts.

## DISCUSSION

We presented here a novel joint experimental and computational framework to quantify PPIs and protein-abundance across a scalable number of cellular conditions with minimal requirements for instrumentation, cellular engineering, and cost. Our framework builds on simple synthetic molecules, centered both on the DNA-caliper, a unique detector of proximal interactions, and the optimized ab-oligos reagents which are enriched for a single oligonucleotide on each antibody molecule. Here, we showcased the capabilities of our framework focusing on a variety of cellular conditions (HEK293T, HeLa, male and female hiPSCs and five patient-derived Glioma cell lines), perturbations (small molecule and inducible expression of oncohistones) and measurements of temporal changes (**Supplementary Figure S16**, **Supplementary Table 3**).

In order to encourage adoption of our framework by the community, we rigorously benchmarked and optimized our methods. We conducted our evaluations in a stepwise manner. We initiated our effort by vetting individual components such as oligonucleotide design parameters and enzymatic reactions, followed by evaluation of the method on recombinant complexes serving as a known ground truth. Cell lysate mixing experiments allowed us quantitatively benchmark our results on a predefined range of PPI and protein abundances. Using these conditions, we demonstrated the reproducibility, specificity, and ability of our framework to quantify changes in the composition of PPIs and protein abundance in multiple samples. Additionally, we built a computational analysis pipeline that processes the readouts and calculates enrichment values for the cellular measurements. Our manuscript includes a step-by-step experimental protocol (**Appendix**) as well as packaged implementation of the computational pipeline along with detailed documentation and tutorial (**Code Availability**).

Notably, the DNA-caliper has the potential to serve as a game-changer in profiling of interactions between biomolecules. We have streamlined the production of the DNA-caliper, to allow easy generation of a variety of designs. For instance, during the optimization steps, we tested more than 20 different versions of the DNA-caliper to evaluate a variety of Anchor sequences, arm lengths, and restriction sites. While here we focused on PPIs and protein abundance, as discussed in detail below, future variations of this unique molecule could allow extensions of our framework to allow the integration of a broad range of additional applications such as detecting protein-DNA, protein-RNA, and DNA-RNA interactions, barcoding the DNA-caliper itself for single-cell measurements, and capturing interactions at different distances by introducing a variety of arm lengths.

In this study, we demonstrate the capabilities of our framework by focusing on PRC2. We studied changes in PPIs and protein abundance across a wide range of cell conditions as well as drug treatments and biological perturbations. The ability of our framework to allow multiplexed measurements creates room for novel observations in addition to confirming expected biological results. We first demonstrated in the HEK293T cell line that upon Tazemetostat (an FDA-approved EZH2 inhibitor^25^) treatment, the interactions between members of PRC2 as well as between PRC2 and its deposited histone mark, H3K27me3, were weakened. Further, in a wider range of cellular conditions – including cancer cell lines, stem cells and patient-derived Glioma – we observed that the presence of oncohistone H3K27M alters the PPIs within the PRC2 complex as well as members of this complex and histone marks such as H3K27me3 and H3K27ac, and that different versions of H3K27M mutations and different patient derived Glioma lines show distinct changes in protein abundance and PPIs within the epitopes we studied.

The Prod&PQ-seq framework has high potential to enhance a wide range of applications, for example: (***i***) enableing the discovery and quantification of previously overlooked PPIs, (***ii***) advancing understanding of the molecular impact of diseases and small molecule treatments, (***iii***) allowing for profiling of individuals at a consortium level, similar to the integration of Olink proteomic data into UK Biobank^39^ (given the low costs of our framework), (***iv***) providing rich PPI data for AI studies, especially neural network-based models that are extremely powerful but require a plethora of input data, and (**v**) extending the framework to single-cell level PPI and protein abundance measurements.

### Limitations of the study

The primary limitation of our framework is that it is currently built on the usage of antibodies to target a pre-defined set of epitopes of interest, which may miss novel PPIs detected by antibody-free methods such as mass spectrometry that provide information at an unbiased, whole proteome level. As immunoprecipitation (IP) efficiency and epitope abundance will vary between antibodies, assembly of properly titrated ab-oligo Prod&PQ-seq panels may encounter challenges if specific reagents dominate signal capture or non-specific signal impedes PPI assessments. The set of ab-oligos also needs to target epitopes that are known to not be present in the surveyed samples (e.g., HA-Tag in this study) for proper estimation and calculation of PPI enrichments. In addition, our approach requires ab-oligos where each antibody molecule is conjugated with exactly one oligo (degree of labeling of 1, DOL=1). While we use purification methods to enrich for the desired DOL=1, antibodies with DOL=2 and possible antibody aggregations are detectable in the purified fractions (**Supplementary Figure S3**). These can form byproducts that associate two copies of the same barcode (self-self associations) from a single antibody, resulting in a false positive detection. In addition to reducing the percentage of usable sequencing reads in a given sample, these byproducts obscure our ability to detect true protein dimerizations. While our current analysis excludes all matching barcodes as possible self-self associations (and thus homodimer complexes are ignored and not quantified), we envision the emergence of methods to produce higher purity DOL=1 ab-oligos would further extend our framework to include the detection and quantification of protein dimerizations.

## RESOURCE AVAILABILITY

### Lead contact

Correspondence should be sent to Christopher Benner and Alon Goren.

### Materials availability

The DNA sequence designs and the instructions for generating the reagents are available in **Appendix.**

## Data and code availability

All Prod&PQ-seq and ChIP-seq data included in the manuscript are available on GEO with accession numbers GSE313772 and GSE313635. The Prod-seq data analysis pipeline has been implemented as a command-line executable Python program and is available on GitHub (https://github.com/Goren-Lab-at-UC-San-Diego/Prod-seq) together with the code used for generating the figures.

## ACKNOWLEDGMENTS

Research reported in this publication was supported in part by CIRM DISC0 14405. (A.G., C.B. and S.H); NIH/NHGRI R21HG010078 (A.G.); Cancer Cell Map Initiative (CCMI) NIH/NCI U54 CA274502 and NIH NCI U54 CA209891 via the award to MPIs Nevan Krogan and Trey Ideker (A.G.), UC San Diego Stem Cell Program - Innovative Projects (A.G.), NIH/NIGMS R35 GM149520 (C.B.). T.X. was supported in part by a CIRM fellowship via the EDUC4-12804 award. Ed.M was supported in part by an institutional award to the UCSD Genetics Training Program from the National Institute for General Medical Sciences, T32 GM145427.

This publication includes data generated at the UC San Diego IGM Genomics Center utilizing an Illumina NovaSeq 6000 that was purchased with funding from a National Institutes of Health SIG grant (#S10 OD026929). We thank Robert Nicol, Chad Nusbaum and Yuting Liu for their input and effort in the early conceptual and method development. The DIPG and EZH2-KO HEK293T cell lines were kind gifts from Michelle Monje-Deisseroth, and Thomas Keck respectively. The H3.1WT, H3.1K27M, H3.3WT and H3.3K27M plasmids were kind gifts from Efrat Shema. We thank AlphaThera for their support in antibody-oligonucleotide conjugations and Rahel Wachs for her help on figure illustrations. We thank Dorit Rousseau and Jenna Satovsky for their contributions to general wet-lab organization and lab administration.

## AUTHOR CONTRIBUTIONS

A.G., C.B., and T.X. conceptualized and designed the study as well as the computational pipeline which was implemented by T.X.; T.X., J.W., Y.S., Y.C., L.Z., Ed.M, T.D. performed the experiments and T.X. Y.C. and C.B. conducted the analyses. J.F., M.N, and C.J.F. provided input in initial establishment of the ab-oligos as well as the conjugation, purification, and biophysical characterization of the ab-oligos used in this study. E.M, S.H. and Y.F. provided key input in designing, interpreting and analyzing the results and the establishment of the framework. A.G. and C.B. supervised the study. All authors contributed to writing and reviewing the manuscript.

## METHODS

### KEY RESOURCES TABLE

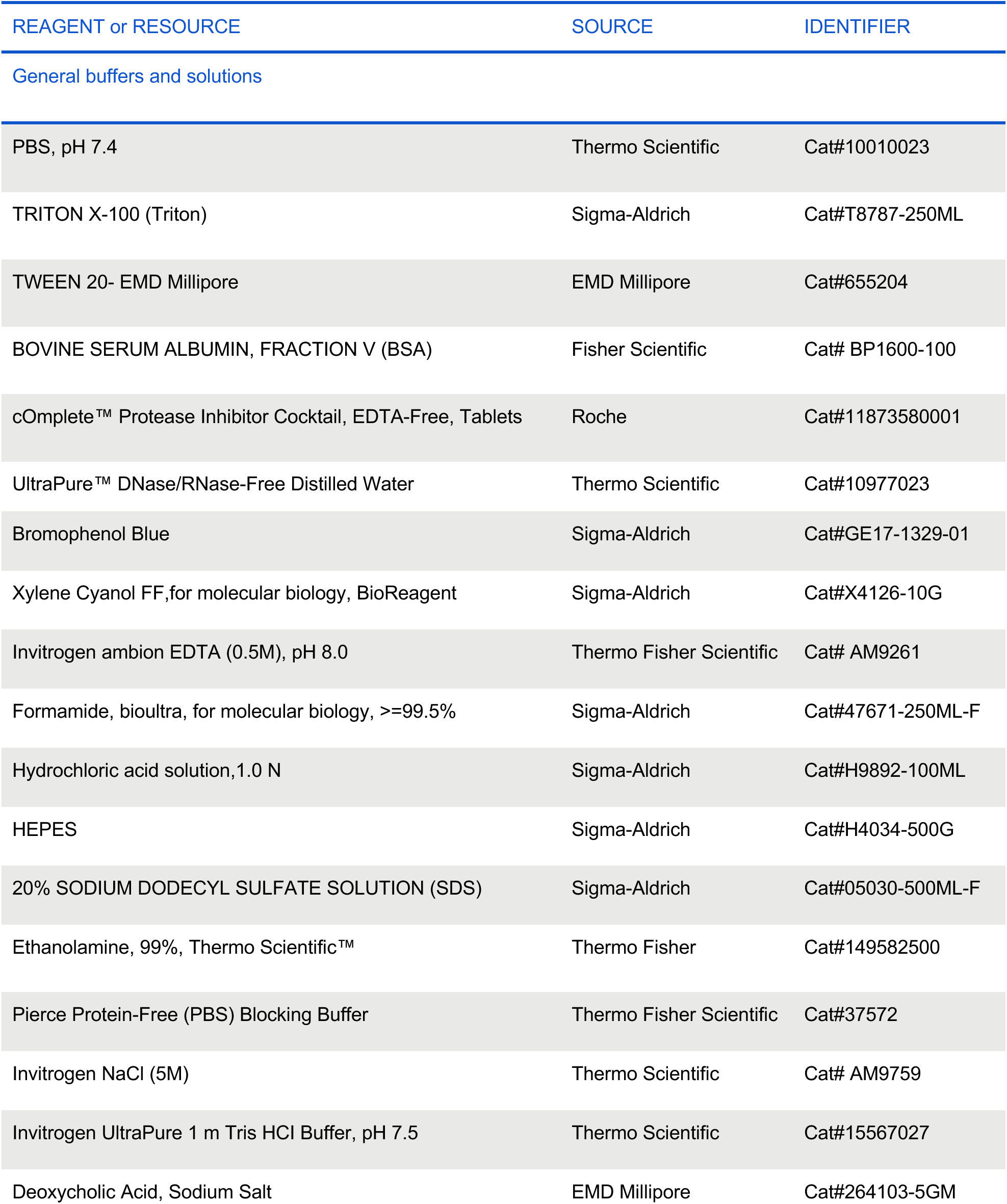
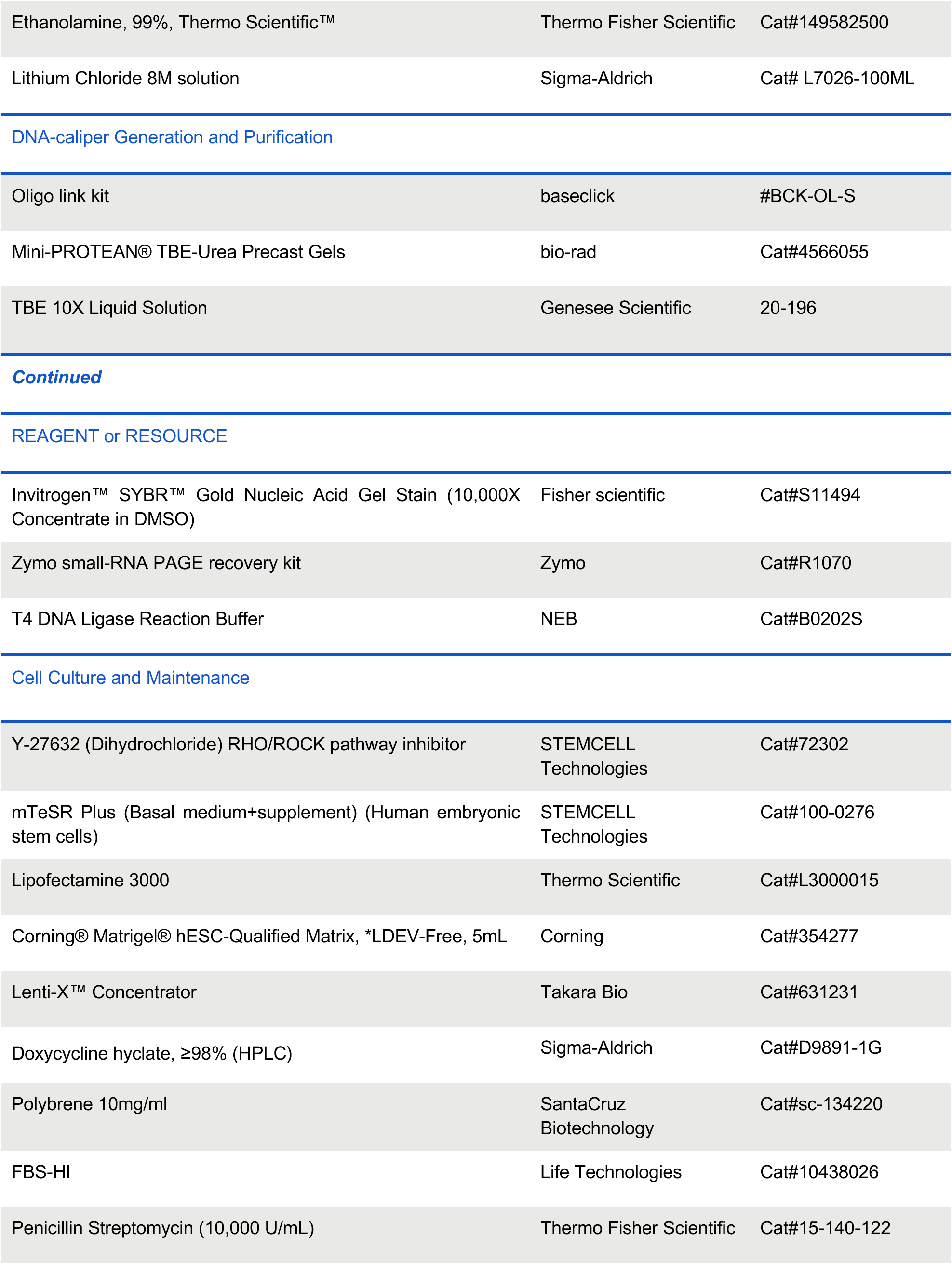
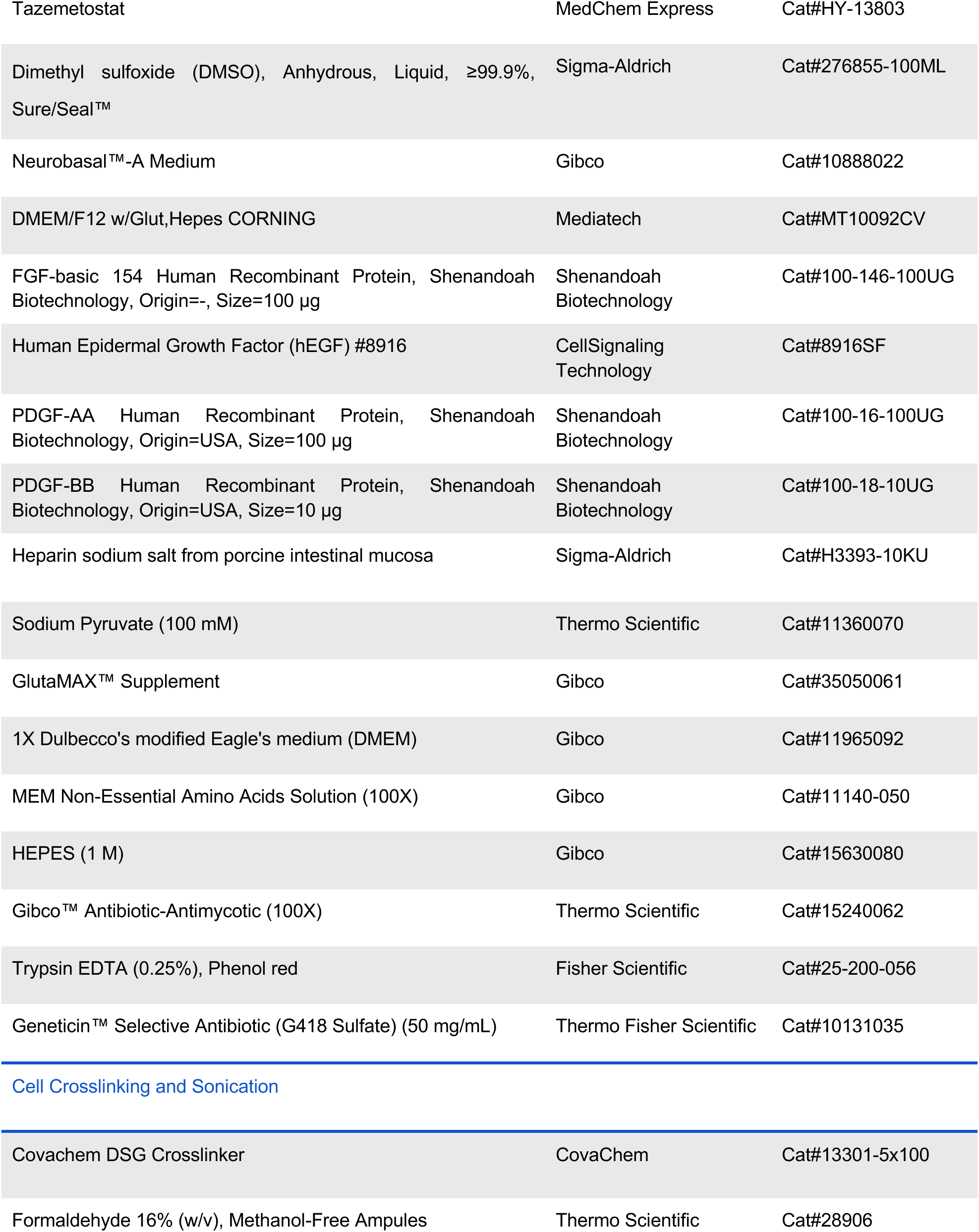
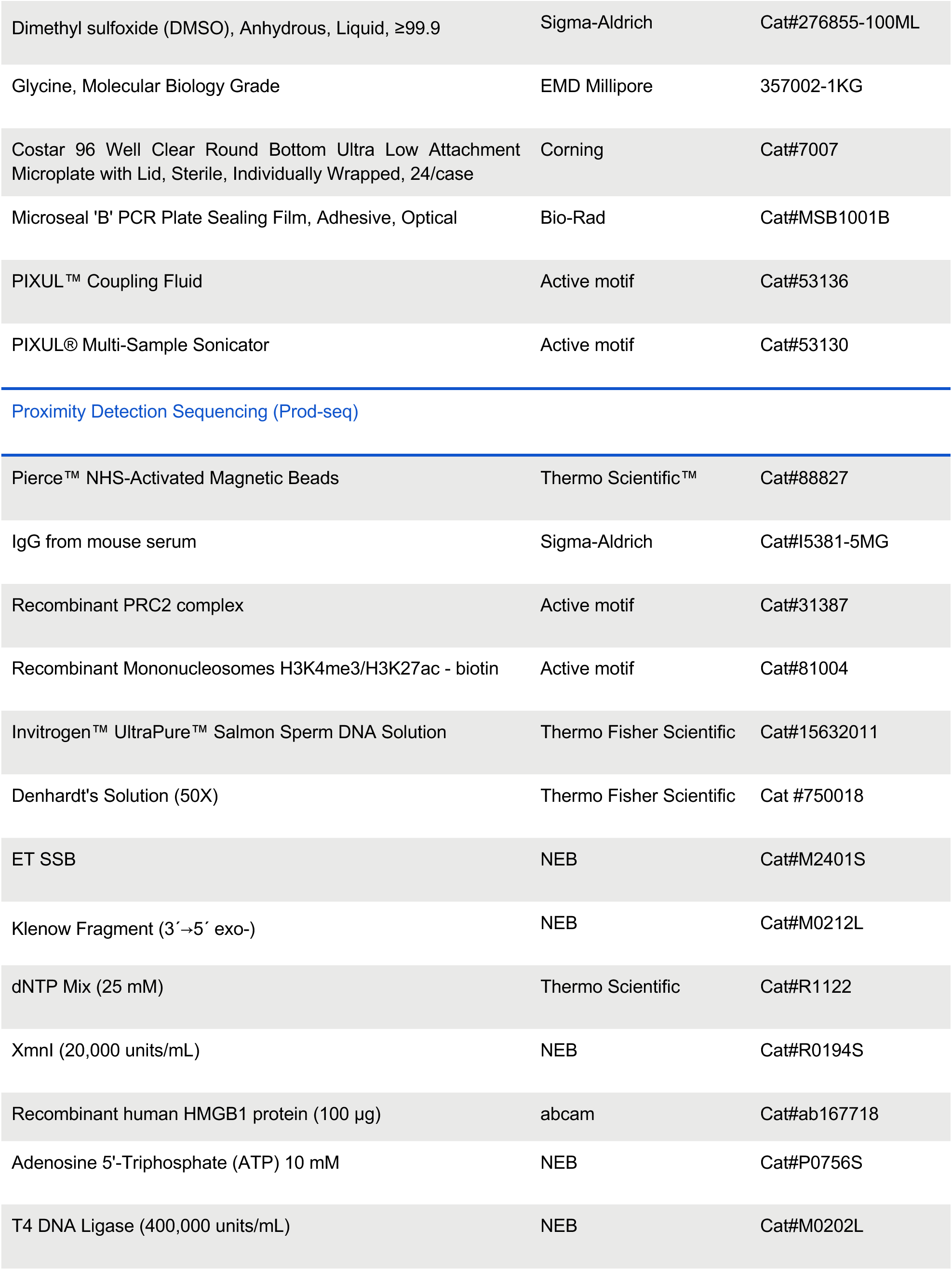
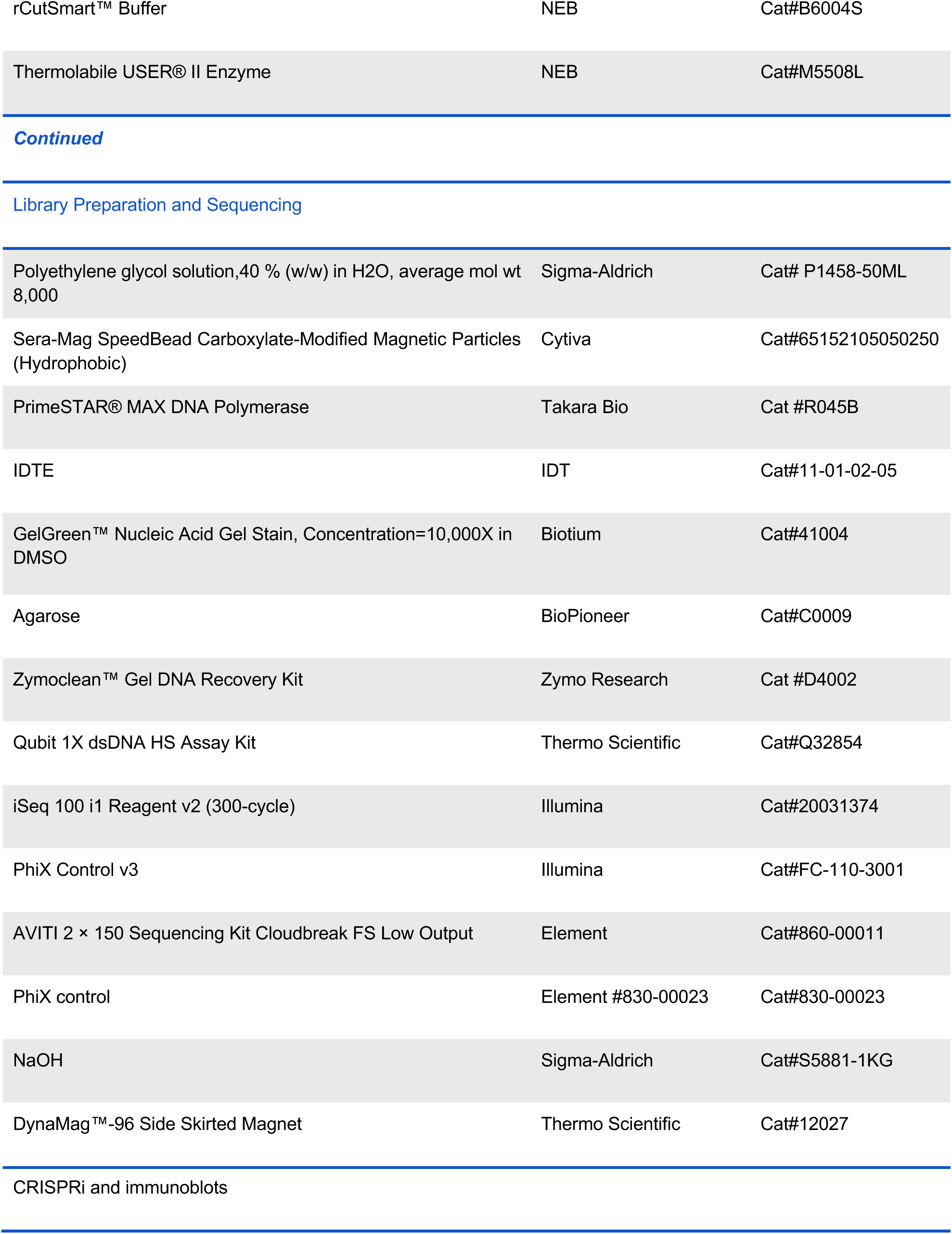
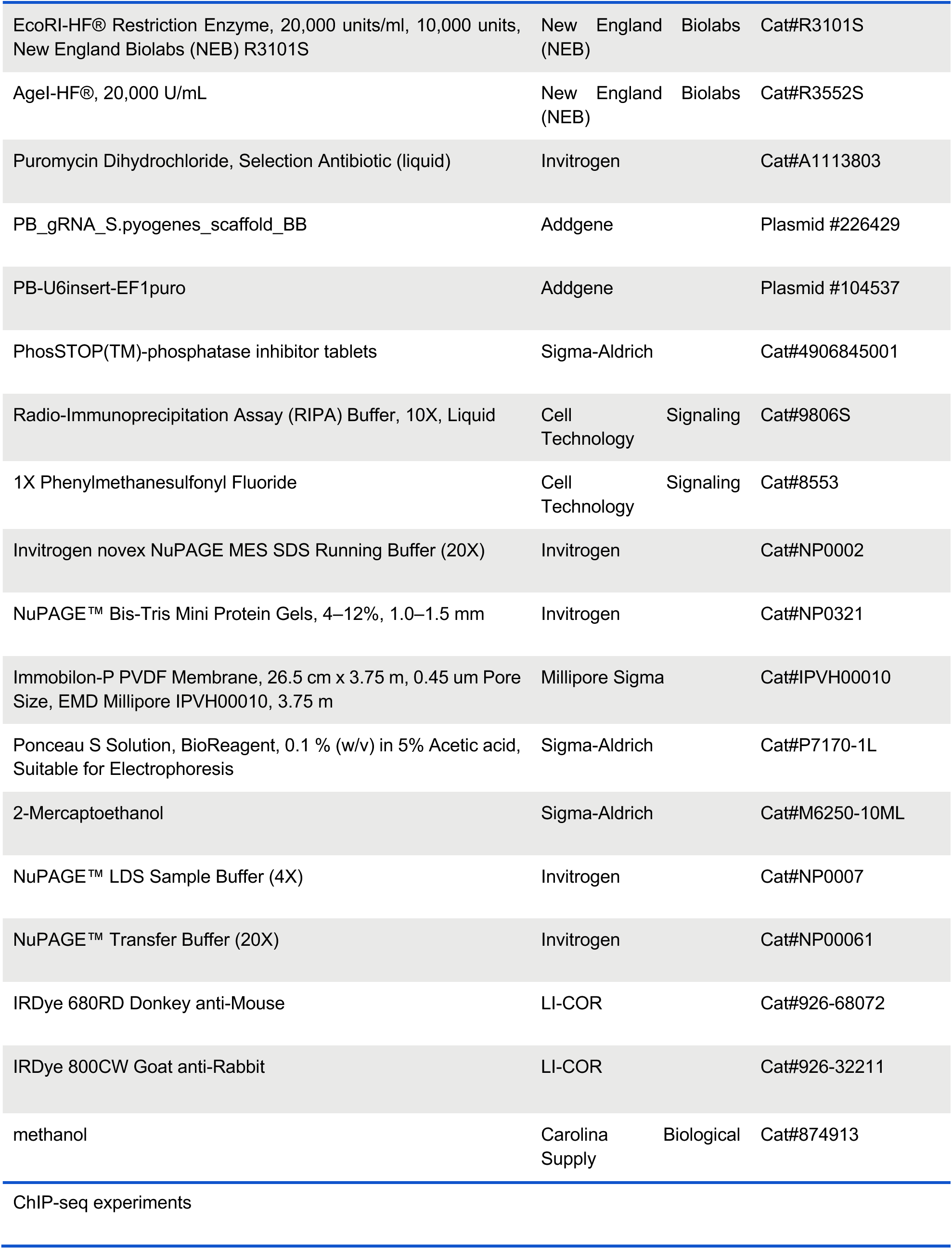
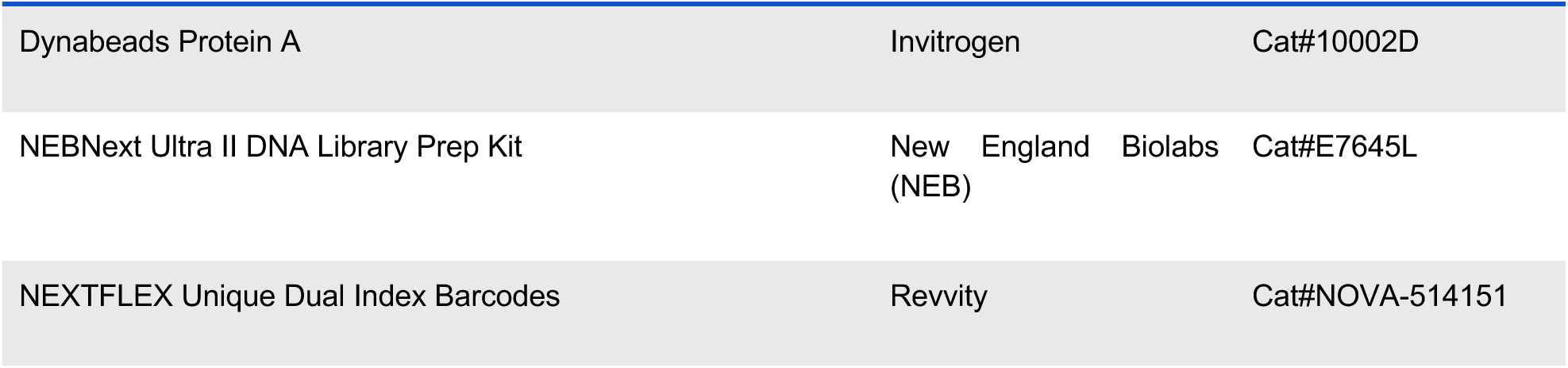

### METHOD DETAILS

#### Cell lines and culture

All cell lines were maintained at 37 °C with 5% CO_2_. HEK293T cells were cultured in Dulbecco’s modified Eagle’s medium (DMEM, Gibco) supplemented with 10% fetal bovine serum (FBS-HI; Life Technologies), 100U/mL Penicillin/Streptomycin (Thermo Fisher) and 2mM L-Glutamine (Thermo Fisher). HEK293T cells were treated with a final concentration of 10µM Tazemetostat (MedChem Express) or equivalent volume of DMSO (Sigma-Aldrich) as control. HEK293T cells were treated for 48 hours before harvesting. HeLa S3 stable cell lines were grown in DMEM (Gibco) supplemented with 10% FBS (Life Technologies), 1 mM Sodium Pyruvate (Gibco) and 2 mM L-Glutamine (Thermo Fisher).

Two strains of human pluripotent stem cells (iPSCs), S02315 (iPSCORE_1_13), and S02316 (iPSCORE_1_14) were gifts from Dr. Kelly Frazer. All hiPSC cells were grown in mTeSR Plus (STEMCELL Technologies) medium on Matrigel-coated plates (Corning).

Five DIPG cell lines were obtained from biopsy or autopsy tissues of DIPG patient tumors. SU-DIPG-XIII (H3.3K27M, female), SU-DIPG 25 (H3.3K27M, female), SU-DIPG 36 (H3.1K27M, female), SU-DIPG 38 (H3.1K27M, female), and SU-DIPG 48 (H3WT, male) were gifts from Dr. Michelle Monje. All DIPG cell lines were verified via Sanger sequencing for their K27M mutation (**Supplementary Figure S15**). DIPG Cells were cultured in 1:1 mixture of Neurobasal™-A Medium and DMEM/F12 w/Glut, Hepes (Mediatech) supplemented with 20 ng/ml of human recombinant H-EGF (Cell Signaling Technology), 20 ng/ml of human recombinant H-FGF-basic-154 (Shenandoah Biotechnology), 10 ng/ml of human recombinant H-PDGF-AA and -BB (Shenandoah Biotechnology), 1% of Glutamax (Gibco), 1% of sodium pyruvate (Thermo Scientific), 1% of MEM Non-Essential Amino Acids Solution (Gibco), 10 mM of HEPES (Gibco), 2 μg/ml of heparin (Sigma-Aldrich), and 1× Anti-anti (Thermo Scientific).

#### Cell crosslinking and preparation of lysates

For all cells for harvest, after growing to about 80% confluency (or at the end of the desired treatment), the cells were crosslinked in 2mM disuccinimidyl glutarate (DSG, CovaChem) for 30 min, followed by 10 min crosslinking with formaldehyde (Thermo Scientific). The crosslinking reactions were quenched with 125 mM Glycine (EMD Millipore) and 0.5% BSA (Fisher Scientific) for 10 min, followed by washes with PBS and protease inhibitor cocktail (PIC; Roche). The crosslinked cell pellets were resuspended and lysed in Prod-seq lysis buffer consists of 10mM EDTA (Thermo Fisher Scientific), 50 mM HEPES (Gibco), 0.5% SDS (Sigma-Aldrich); 133.3 µL used for resuspending each 1 million crosslinked cells) followed by sonication using the PIXUL® Multi-Sample Sonicator (Active Motif, Cat#53130). Detailed protocols for crosslinking the cells for harvest and the subsequent lysing and sonication are provided in the Appendix.

#### Generation of doxycycline inducible K27M

Plasmids of doxycycline(dox)-inducible H3.1WT, H3.1K27M, H3.3WT, H3.3K27M were gifted from Dr. Efrat Shema’s Lab (The Weizmann Institute). Lentiviral (third generation) packaging was performed by Lipofectamin 3000 (Thermo Scientific) transfection of HEK293T with appropriate plasmids. Virus-containing supernatants were collected after 48h transfection, filtered and concentrated with lenti-X (Takara Bio), and added to HeLa S3, S02315, and S02316, supplemented with 8 μg/mL Polybrene (SantaCruz Biotechnology). Infected cells were selected with 0.5 mg/mL G418 (Thermo Fisher Scientific).

#### Dox-inducible expression and washout

The HeLa and hiPSCs (S02315/S02316) lines with H3.1K27M or H3.3K27M integrated by lentivirus were induced with 1 µg/mL doxycycline (Sigma-Aldrich) for 6 days for expression. Cells were dual-crosslinked at the end of the 6-day doxycycline treatment for expression-treatment cell harvest for Prod-seq. For washout experiments, a subset of the plates were crosslinked at the end of the 6-day doxycycline treatment and the remaining plates had their cell culture medium replaced with the matching medium without doxycycline as washout treatment. For HeLa cells, subsets of cells were subsequently harvested by crosslinking at the 6h, 24h, and 96h timepoints after the medium was replaced. For hiPSCs, washout cells were maintained in doxycycline-free medium and crosslinked after 5 days. The crosslinked cells are lysed and sonicated with PIXUL® Multi-Sample Sonicator for Prod-seq.

#### Proximity detection sequencing (Prod-seq)

For each Prod-seq sample, 40 μL of Pierce NHS-activated magnetic beads (Thermo Scientific) were washed once with ice-cold 1mM HCl and resuspended in 110 μL PBS (Thermo Scientific) working solution which contains 1X PIC (Roche) and 0.1% Triton (Sigma-Aldrich) in PBS buffer. Cell lysates were spinned down at 4 °C and 16,000g for 10 min to remove cell debris. 40 μL of the supernatant, approximately 300,000 cells, were added to the beads suspension (for cell lysate mixing experiment, the 40 μL volume is split between the HEK293T and SU-DIPG36 cell lysates where the lysate ratios are the volume ratios). Bead-cell lysate mixtures were incubated for 2 hours at room temperature on HulaMixer (Thermo Fisher), with frequent rotating and shaking, same below. The beads were subsequently washed twice with 90 μL PBS working solution, and then quenched with 90 μL of 3M ethanolamine (Thermo Fisher Scientific; pH 9.0) with 0.1% Triton for 2 hours at room temperature. The beads were then washed again and then blocked with 180 μL of blocking solution containing 2.78 ng/μL Mouse IgG (Sigma Aldrich), 27.78 ng/μL Sheared Salmon Sperm DNA (Thermo Fisher), 1X PIC, 5X Denhardt’s solution (Thermo Fisher Scientific), 0.1% Triton, 10% BSA (Fisher Scientific) in PBS-based Pierce Protein-Free blocking solution (Thermo Fisher Scientific) for 30 min on HulaMixer. Freshly-pooled ab-oligos (**Appendix**) are then added to the beads suspension and incubated overnight at 4 °C on a rotating mixer.

After the overnight incubation, the beads were washed 3 times with 1X PIC in Wash Buffer I (WBI: 20 mM Tris-HCl pH 7.5, 1% Triton X-100, 2 mM EDTA, 150 mM NaCl and 0.1% SDS), and then washed 3 times with 1X PIC in Wash Buffer III (WBIII: 10 mM Tris-HCl pH 7.5, 0.7% Sodium Deoxycholate, 1% Triton X-100, 250 mM LiCl, 1 mM EDTA), and then washed 1 time with 1X PIC in TET buffer (10 mM Tris-HCl pH 7.5, 0.2% TWEEN 20, 1 mM EDTA). Then the beads were resuspended with 1X PIC in TET buffer. From here, half of the beads suspension were taken for Prod-seq PPI profiling,and the other half of the beads were taken for the optional protein quantification subroutine.

For PQ-seq, the supernatant was removed and the beads were resuspended in 8 μL of freshly prepared 0.05% Tween (EMD Millipore) and 1 X PIC in 1X rCutSmart buffer (NEB). 2 μL of 1 uM protein quantification capture oligo (**Appendix**) was then added to the beads suspension and incubated for 30 min at room temperature. The beads suspension then underwent Klenow Fragment (3′→5′ exo-) (NEB) strand extension for 20 min at 25 °C with heat inactivation at 75 °C for 20 min; the extension product is then further diluted using 0.05% Tween and 1 X PIC in 1X rCutSmart buffer to 50 μL. Sequencing libraries are subsequently generated using PCR with PrimeSTAR Max Master Mix (Takara Bio).

For the Prod-seq PPI profiling, the supernatant was removed and the beads were resuspended in 5 μL of freshly prepared 0.05% Tween, 1 X PIC in 1X rCutSmart buffer. 10 nM DNA-calipers were pre-incubated in 5% ETSSB (NEB) at 55 °C for 10 min and then at 37 °C for at least 30 min; 5 μL of the DNA-caliper incubation mix was then added to the beads suspension and with an incubation at room temperature for 30 min. Following the annealing, the workflow proceeded sequentially as follows: strand extension using Klenow Fragment (3′→5′ exo-; NEB) for 20 min at 25 °C, wash the beads with WBI, WBIII and TET each for one time, heat inactivation of the residual Klenow fragment at 75°C for 20 min, XmnI (NEB) digestion (37°C for 1 hour, then heat inactivate at 65°C for 20 min) to release oligonucleotide product from the antibodies and beads, remove the beads and keep the supernatant containing the released oligonucleotide product, further dilute the samples using 0.05% Tween and 1 X PIC in 1X rCutSmart buffer, HMGB1 (Abcam) incubation (30 min at room temperature), and T4 ligase (NEB) ligation (25°C for 20 min, then heat inactivated at 65°C for 10 min). The resulting circular double-stranded nucleotides were then linearized using Thermolabile USER II enzyme (NEB) at 37°C for 30 min and heat inactivation at 65°C for 10 min. The DNA products were size-selected using Sera-Mag SpeedBead (Cytiva) with 8.5% PEG 8000. Sequencing libraries are subsequently generated using PCR with PrimeSTAR Max (Takara Bio) Master Mix.

Sequencing of the libraries were performed on an Element AVITI system using Cloudbreak FS kits with either Low or High output (depending on the sequencing depth needed) with 151 bp reads for both reads 1 and 2.

The complete, detailed protocol for Prod-seq is provided in **Appendix**.

#### Prod-seq on recombinant protein complexes

Prod-seq involving recombinant protein complexes used recombinant PRC2 (Active Motif) and/or recombinant mononucleosomes with H3K4me3 and H3K27ac modifications (Active Motif). When adding samples to the NHS magnetic beads, (i) recombinant PRC2 samples used 5.65 μL of recombinant PRC2 (original concentration 1.4 μg/μL, 1:250 diluted using PBS working solution before using) to replace the cell lysate; (ii) recombinant mononucleosomes with H3K4me3 and H3K27ac modifications used 3 μL of recombinant mononucleosomes with H3K4me3 and H3K27ac modifications (original concentration 0.36 μg/μL, undiluted) to replace the cell lysate; (iii) recombinant mixing sample used 2.825 μL of recombinant PRC2 (original concentration 1.4 μg/μL, 1:250 diluted using PBS working solution before using) and 3 μL of recombinant mononucleosomes with H3K4me3 and H3K27ac modifications (original concentration 0.36 μg/μL, undiluted) to replace the cell lysate. For all recombinant protein complex Prod-seq samples, additional PBS working solution was added to make up for the volume difference between recombinant protein complexes added and the original volume of cell lysates (40 μL).

#### Antibody-oligonucleotide conjugation and purification

All of the antibodies used in this study were recombinant monoclonal (**Supplementary Table 2).** The conjugation and purification processes to generate the DOL=1 ab-oligos used for producing main Prod&PQ-seq results of this manuscript were customized by Cell Signaling Technology (CST) with catalog numbers of the ab-oligos listed in **Supplementary Table 2**. For the additional ab-oligo conjugates mentioned in **Supplementary Figure S2**, the DOL 2.5-3.8 ab-oligos are generated by the CST antibody conjugation team, and the DOL 1&2 mixture ab-oligos were conjugated and purified by AlphaThera. Both CST and AlphaThera used the same recombinant monoclonal antibodies for oligo conjugation

#### Prod-seq sequencing result processing and analysis

Sequencing results for Prod&PQ-seq are processed and analyzed using custom python scripts following the model detailed in **Supplementary Note S4**. The pipeline used is made available on GitHub (https://github.com/Goren-Lab-at-UC-San-Diego/Prod-seq).

For **Figures 3F** and **4F**, as we did not observe the PPI enrichment values to be normally distributed, the p-values are calculated from two-sided Mann–Whitney U tests. Given the sample sizes for those Mann–Whitney U tests (n_1_ = n_2_ = 4), the smallest theoretically possible p-value is 0.029, meaning direct multiple hypothesis corrections on those p-values are inadequate with this sample size.

#### ChIP-seq experiments

The crosslinked cell pellets from HeLa H3.1K27M and H3.3K27M used for Prod-seq are used for ChIP-seq. H3K27ac ChIP-seq were performed with a dual spike in method as described in Patel et al (with Drosophila S2 and yeast cells^40^); H3K27M ChIP-seq were performed without spike-ins. The standard ChIP-seq experiment steps were carried out as previously reported in spa-ChIP-seq^32^. Briefly, the HeLa, S2, and yeast cell pellets were suspended with LB3 lysis buffer and sheared with Active Motif’s PIXUL Multi-Sample Sonicator. For H3K27ac ChIP with dual spike-in, a mix of S2 and yeast chromatin was added to HeLa chromatin. For H3K27M ChIP samples, only the sonicated HeLa chromatin was added. ChIP samples were incubated overnight with Dynabeads Protein A (Invitrogen) and either H3K27ac or H3K27M antibody (CST). On the next day, ChIP samples were washed with WBI, WBIII and TET and resuspended in TT buffer (0.05% Tween 20, 10 mM Tris pH 8.0). Sequencing libraries of ChIP and input samples were constructed with NEBNext Ultra II DNA Library Prep Kit (NEB) and NEXTFLEX Unique Dual Index Barcodes (Revvity). Libraries were size-selected with Sera-Mag SpeedBead + 8.6% PEG 8000 + 0.8M NaCl and PCR amplified with primers Solexa 1GA (IDT, AATGATACGGCGACCACCGA) and Solexa 1GB (IDT, CAAGCAGAAGACGGCATACGA). Final cleanup of the libraries were done with Sera-Mag SpeedBead + 8.5% PEG 8000 + 1M NaCl. The concentrations of the libraries were measured using a Qubit fluorometer and the sizes were estimated using 2% agarose gel electrophoresis. Libraries were sequenced on the Element AVITI system along with the Prod-seq samples.

#### ChIP-seq data analysis

Sequencing fastq files were trimmed with *fastp*^41^. Reads were aligned using default parameters of *bwa*^42^ *mem* to either the human reference genome (hg38) or, for ChIP samples with dual spike-in, a concatenated genome containing hg38, dm6 (with suffix “_dm6”), and sacCer3 (with suffix “_sac3”) chromosomes^40^. Duplicated reads were removed with *samtools*^43^ *markdup*. For reads aligned with the concatenated genome for dual spike-in, bam files were sorted and split by chromosome using bamUtil^44^ *splitChromosome*; individual chromosome bam files for each species were then combined with *samtools merge*, and suffixes (“_sac3” or “_dm6”) were removed from the chromosome names of the spike-in species^40^. HOMER^45^ *makeTagDirectory* was used to generate tag directories for all ChIP-seq data. For H3K27ac ChIP-seq samples with dual spike-in, the normalization factor for each ChIP sample was calculated and applied as described in Patel et al.^40^. Peaks were called using HOMER *findPeaks* with “*-style histone*” for histone mark peaks, and *“-style histone -size 1000 -minDist 2500*” for broader peaks, using matched input samples as background. The bigWig files were created with deepTools^46^ *bamCoverage* (calculated dual spike-in normalization factor was applied to *--scaleFactor*) and visualized using IGV^47^ genome browser. Peak overlaps were identified with the default parameters of *bedtools*^48^ *intersect.* Heatmaps and histograms centered on peaks were generated using deepTools *computeMatrix* and *plotHeatmap*. For the peak overlap analyses in SU-DIPG lines, the published data are downloaded using the accession numbers GSE94259 and GSE126319 from GEO. The same analyses process as above were performed except that the replicates were combined into single meta-experiments prior to peak calling and that peak overlaps were identified with HOMER’s *mergePeaks* program.

#### Immunoblots

Harvested cell cultures were lysed for 30 minutes on ice, vortex every 10 minutes, with 1x RIPA buffer (Cell Signaling Technologies) with 1X PIC, 1X Phenylmethanesulfonyl Fluoride (PMSF; Cell Signaling Technology), and 1X phosphatase inhibitor (Sigma-Aldrich). The lysed cells were sonicated using the PIXUL® Multi-Sample Sonicator for 24 minutes and subsequently centrifuged for 10 minutes at 4°C for 14,000 rcf. From the cell lysates, protein samples were prepared with NuPAGE LDS Sample buffer x1 (Invitrogen) and 4% 2-Mercaptoethanol (Sigma-Aldrich). Proteins were run in NuPAGE 4-12% Bis-Tris Gel (Invitrogen) at 200V in NuPAGE MES SDS running buffer x1 (Invitrogen). After separation, proteins in gel were transferred to the membrane (Millipore Sigma) at 100V, 90 min, with NuPAGE Transfer Buffer x1 (Invitrogen) with 20% methanol (Carolina Biological Supply). To check transfer, wash the membrane with DI water, stain with Ponceau (Sigma-Aldrich) and image. After checking, wash off the Ponceau with DI water and TBS (150 mM NaCl, 15 mM Tris-HCl, and 4.6 mM Tris base). Block the membrane in PBS with 5% BSA for 1 hour on a shaker. After blocking, wash the membrane with TBS for two times and TBST (150 mM NaCl, 15 mM Tris-HCl, 4.6 mM Tris base, and 0.05% (v/v) Tween-20) for one time, 5 minutes for each wash. Incubate the washed membrane with primary antibodies overnight at 4°C. Same as Prod-seq, but unconjugated antibodies were used as primary antibodies in immunoblots. After overnight incubation with primary antibodies, the membranes were washed with TBS for two times and TBST for one time, 5 minutes for each wash. The membrane was incubated with fluorescent secondary antibodies (LI-COR) for 1 hour. Blotted with secondary antibodies, the membrane was washed another two times with TBS, and one time with TBST, 5 minutes each wash. After washing, the membranes were imaged. Immunoblots were performed to validate antibodies and cell lines.

Band intensities in immunoblot images were analyzed with ImageJ. For each target protein or loading control band, the greyness of its nearby background is subtracted from the greyness of the band to compute the background-subtracted intensity for each band. For each protein target band, the normalized intensity was then calculated as the ratio of its background-subtracted intensity to that value of the corresponding loading control band.

#### CRISPRi

HeLa S3 cells with EED, EZH2 or SUZ12 knocked down were generated using CRISPRi. A gRNA donor backbone vector, PB_gRNA_S.pyogenes_scaffold_BB (Addgene plasmid #226429) was generated using the plasmid PB-U6insert-EF1puro (Addgene Plasmid #104537). The PB-U6insert-EF1puro plasmid was digested with EcoRI-HF (NEB) and AgeI-HF (NEB) followed by ligation of annealed oligos from IDT containing a BbsI Golden Gate cloning site upstream of the S. Pyogenes gRNA scaffold sequence. A CRISPRi gRNA Donor Vector Library was generated for EED, EZH2, or SUZ12 gene by pooling together six different protospacer sequences targeting the promoter. The promoter was selected from the Human CRISPR Inhibition Pooled Library^49^. The HeLa S3 cell line expressing dCas9-tag-BFP-KRAB (gift from Dr. Komor’s lab at UCSD) was transfected with a human PiggyBac transposase expressing plasmid69 as well as the CRISPRi gRNA Donor Vector Library using Lipofectamine 3000 Transfection Reagent (Thermo Scientific) at a 1:2.5 molar ratio of transposase to transposon vector. Cells were selected with puromycin (Invitrogen) for ∼5 days.

## SUPPLEMENTARY INFORMATION

**Supplementary Figure S1.**
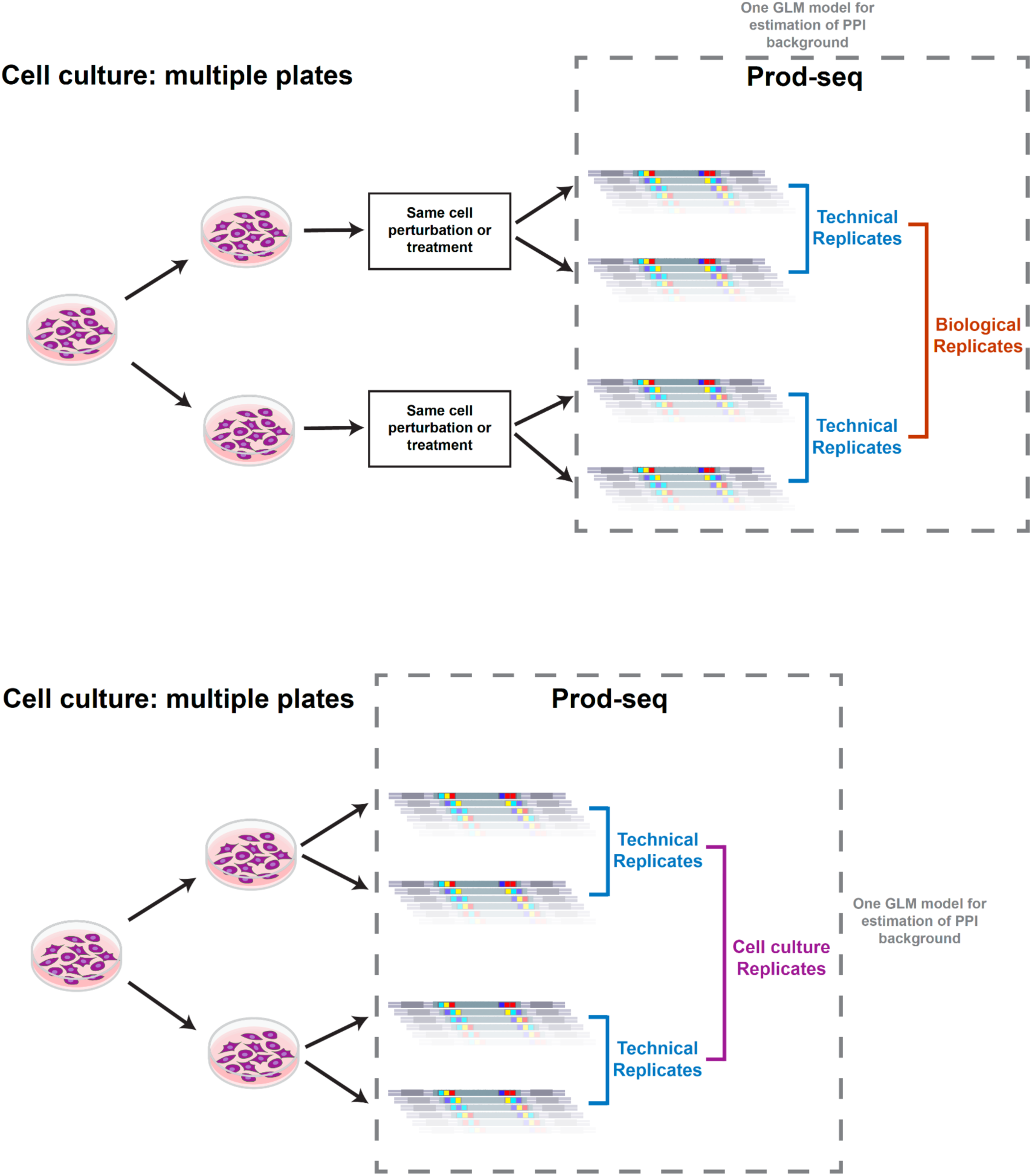
Prod-seq uses GLM to estimate the background levels for samples that are assumed to have similar distributions. Top: situations where cells are split into separate plates and go through the same treatment are termed “biological replicates” in this manuscript. Prod-seq samples that use the lysate from the same plate(s) of cells are termed “technical replicates”. Prod-seq results for biological replicates and technical replicates of the treatments are grouped into the same two-round GLM analysis, together with other treatment conditions to be compared together, to estimate the same background distribution function. Bottom: situations where cells are simply split into different plates and then harvested for Prod-seq after growing to ideal confluency are termed “cell culture replicates”. Cell culture replicates are also grouped into the same two-round GLM analysis, together with other cell lines to compare with, to estimate the same background distribution function.

**Supplementary Figure S2.**
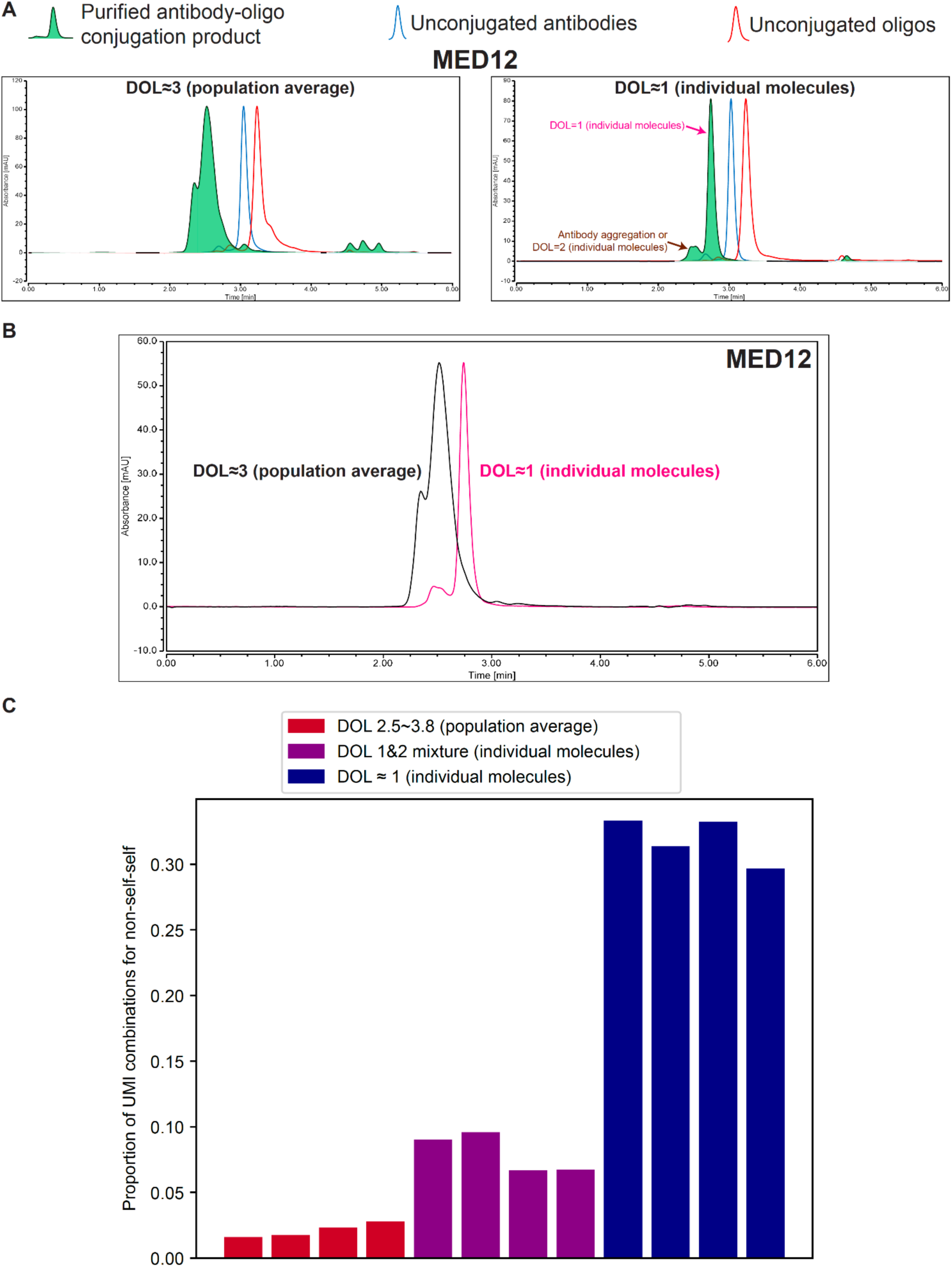
Controlling for the degree of labeling (DOL) of the ab-oligos is important for reducing byproducts originating from multiple oligos on the same antibody molecule. **(A)** Comparison of two batches of MED12 ab-oligos as an example using size-exclusion chromatography (SEC) images. Left: each antibody molecule is conjugated to a varied number of oligos, and the average DOL per antibody molecule is about 3 (population average). Right: most of the antibody molecules in the green-shaded fraction are conjugated to exactly one oligo per antibody molecule (individual molecule level of DOL ∼1). Pink arrow: the peak that corresponds to DOL=1 for each molecule; brown arrow: the peak corresponds to residual antibody aggregation and DOL=2 conjugates. **(B)** Overlay of the two green-shaded MED12 conjugates shown in panel A. **(C)** Enriching for DOL=1 noticeably reduces the self-self byproducts. Each group shows four Prod-seq samples conducted using a group of 12 ab-oligos with different DOLs (as labeled by color).

**Supplementary Figure S3.**
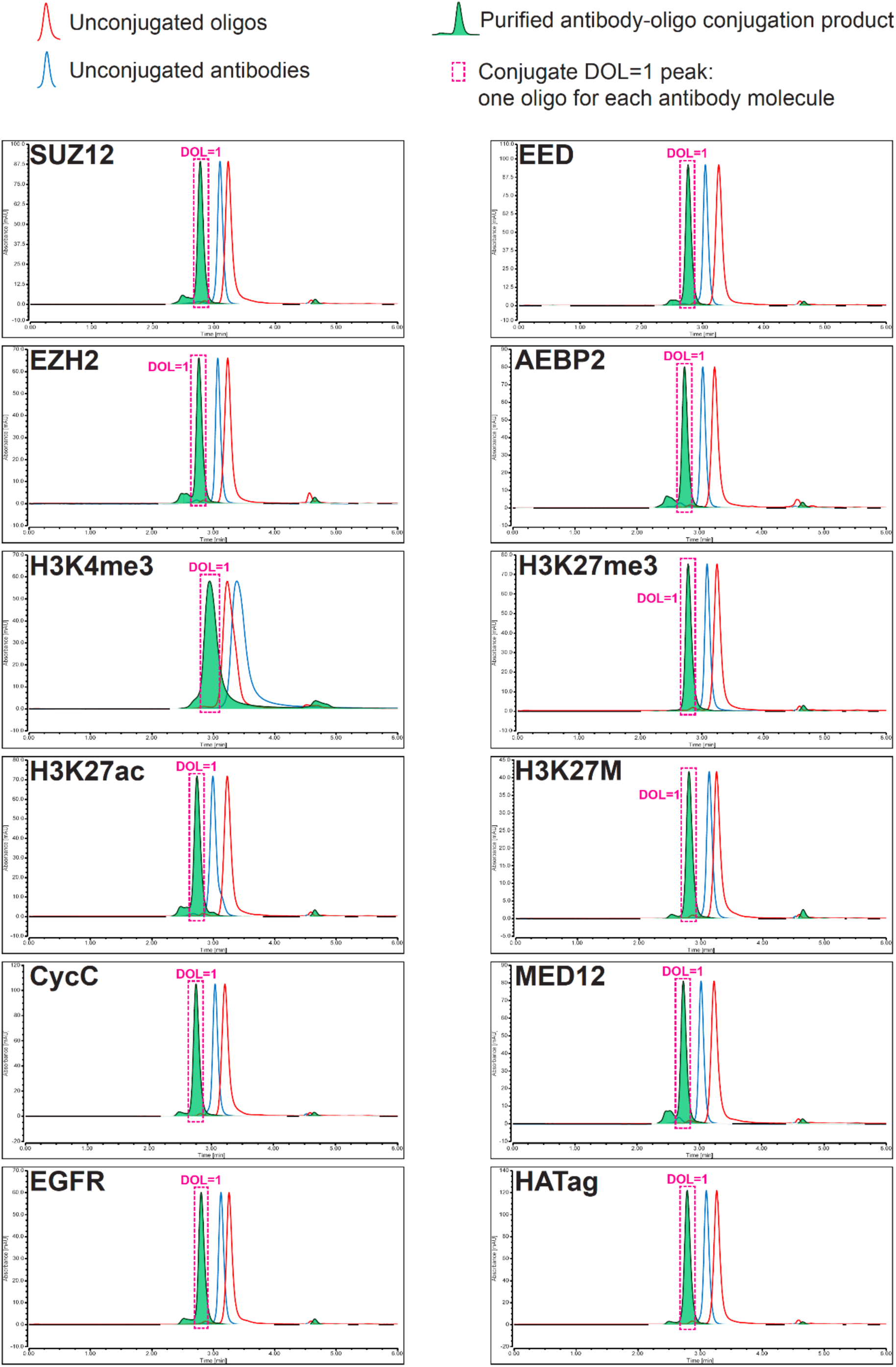
ab-oligos used for Prod-seq are purified to enrich for DOL=1 per antibody molecule. Size-exclusion chromatography (SEC) chromatograms of the purified ab-oligos (green-shaded) compared with unconjugated antibodies (blue) and unconjugated oligos (red). Pink box: the peak that corresponds to DOL=1 for each molecule.

**Supplementary Figure S4.**
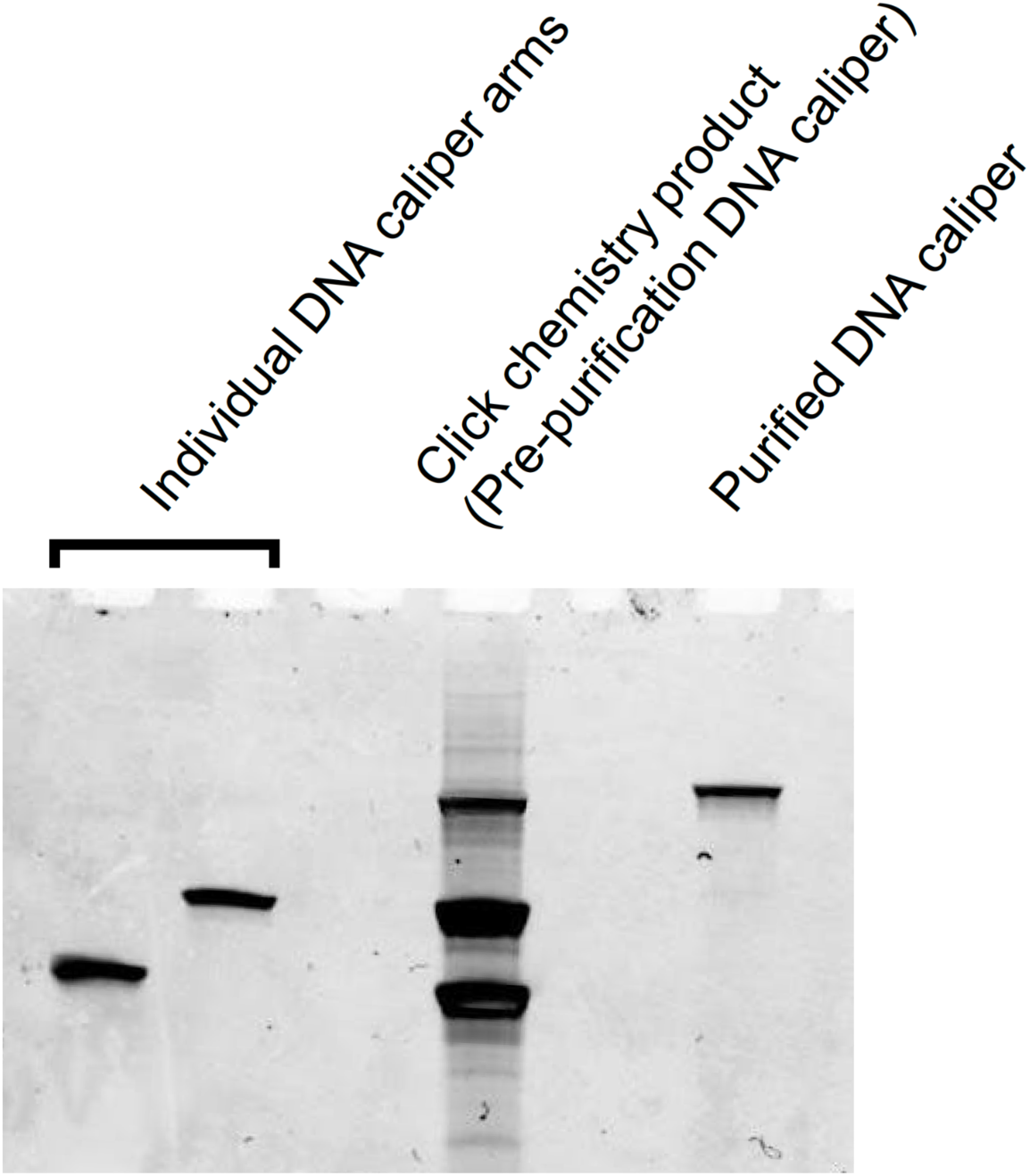
Example gel for the production and purification of a DNA-caliper. Left two lanes: the two individual DNA-caliper arms prior to click-chemistry conjugation. Middle: click-chemistry product (a mixture of the individual arms and the correctly connected DNA-calipers). Right: the purified DNA-caliper generated by gel extraction of the top band of the click-chemistry product (more details in **Appendix**).

**Supplementary Figure S5.**
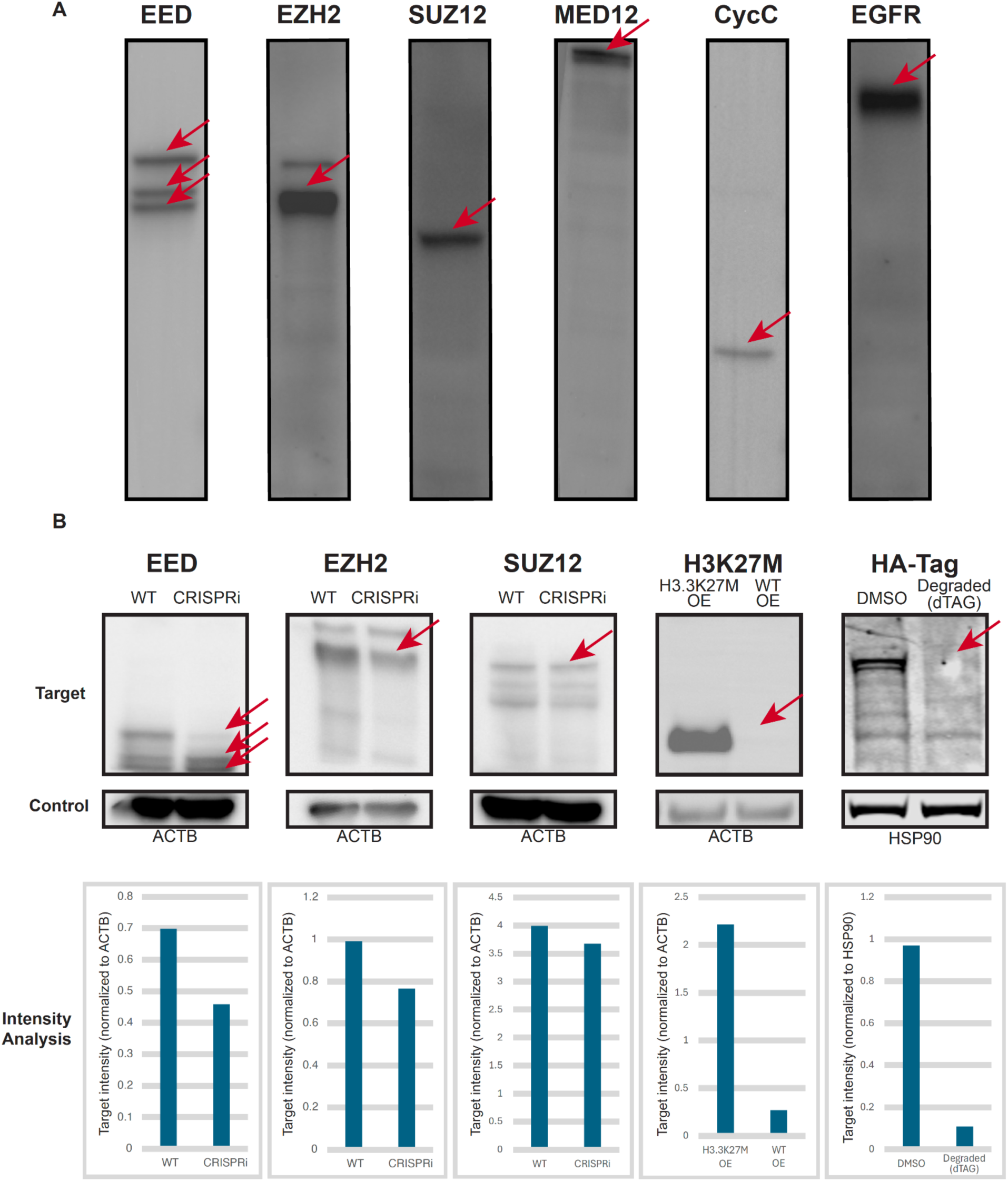
Evaluation of antibodies by immunoblots. **(A)** Antibodies used to generate the ab-oligos are evaluated for specificity. Example immunoblot images for the evaluation of the specificity of each antibody. Red arrows: expected bands. **(B)** Evaluation of a subset of the antibodies using CRISPRi. Top: example immunoblot images of WT versus CRISPRi targeting EED, EZH2, and SUZ12; red arrow: expected bands. Bottom: ImageJ analysis of the band intensities for WT versus CRISPRi conditions.

**Supplementary Figure S6.**
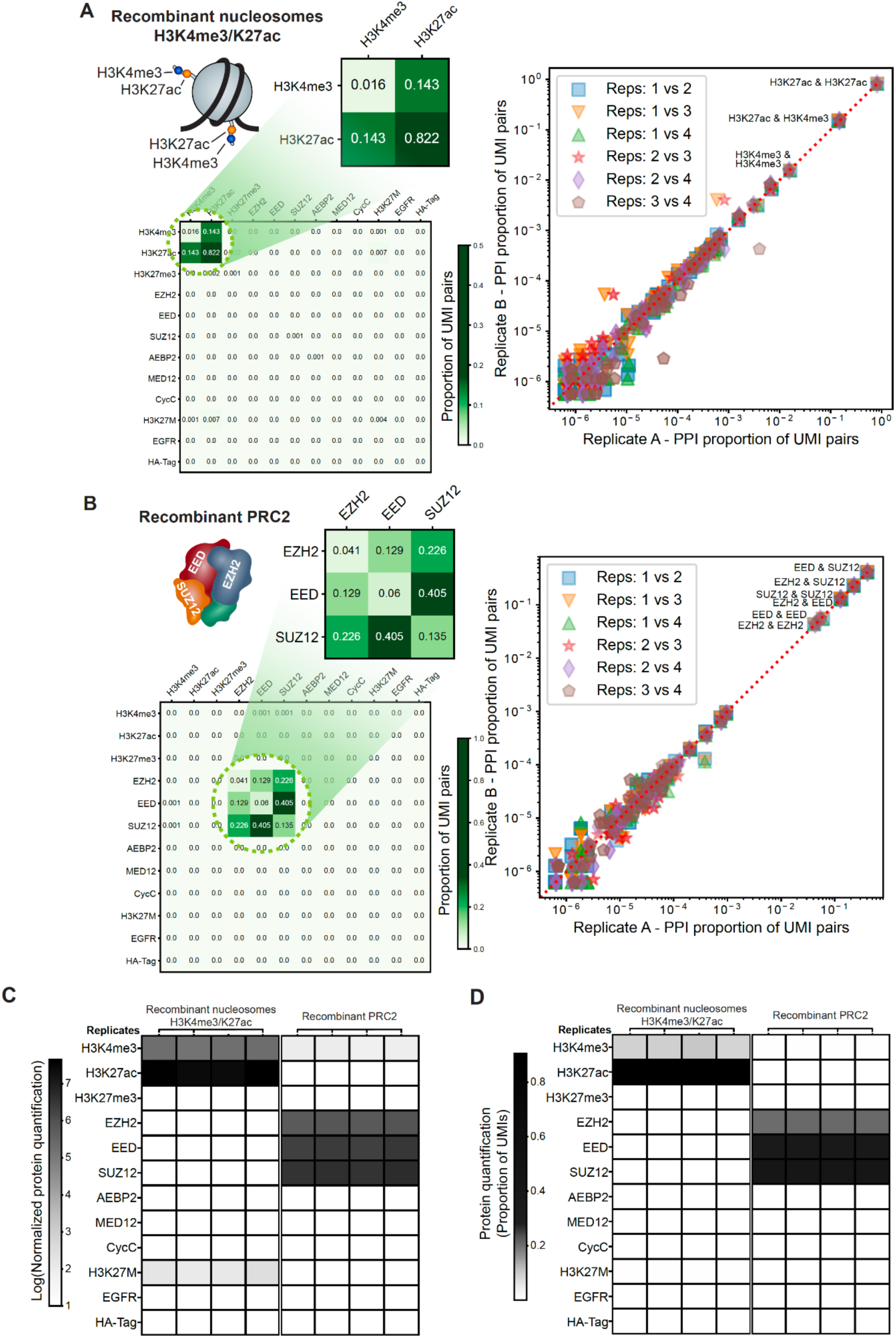
Recombinant protein complexes provide a ground truth for demonstrating the ability of Prod&PQ-seq to generate specific and reproducible PPI detection and protein abundance profiles. **(A)** Prod-seq generates reproducible results with low noise levels on recombinant mononucleosomes with H3K4me3 and H3K27ac modifications. Left: Example real Prod-seq data of UMI pairs distribution and the heatmap zoomed in on the true signals. Right: scatter plot comparing the reproducibility among four technical replicates of Prod-seq on recombinant nucleosomes with H3K4me3 and H3K27ac modifications. **(B)** Prod-seq generates reproducible results with low noise levels on recombinant PRC2. Left: Example Prod-seq data of UMI pairs distribution and the heatmap zoomed in on the true signals. Right: scatter plot comparing the reproducibility among four technical replicates of Prod-seq on recombinant PRC2. **(C)** PQ-seq detects the protein targets in recombinant complexes. For protein quantification readout, the UMI counts of each protein barcode is normalized by the positive control UMI counts (MED12 and Cyclin C, **Methods**) to plot in the heatmap. The resulting values are averaged among the replicates for each condition. Left: recombinant mononucleosomes with H3K4me3 and H3K27ac modifications. Right: recombinant PRC2. **(D)** Heatmap of protein quantification UMI counts normalized by the total number of UMIs for the recombinant complexes. In each sample, the UMI count of each protein is divided by the sum of all proteins’ UMI counts in that sample as an alternative normalization method for cross-sample comparison. The resulting values are averaged among the replicates for each condition.

**Supplementary Figure S7.**
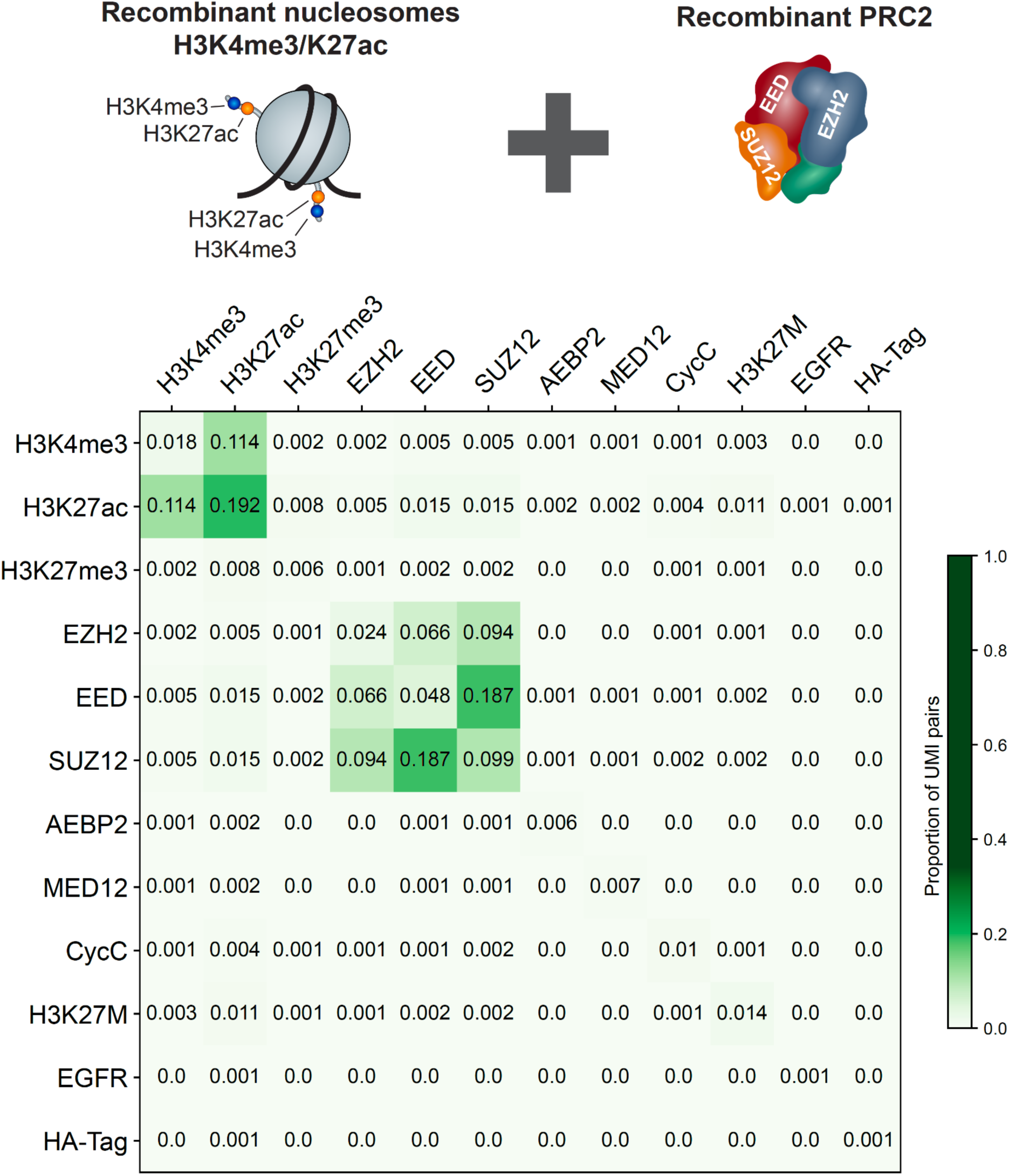
Protein co-localization on the NHS beads does not lead to detection of false interactions. Recombinant PRC2 and recombinant mononucleosomes with H3K4me3 and H3K27ac modifications were mixed for Prod-seq (**Methods**) to determine if proteins located nearby on the NHS beads could lead to a false positive signal. Prod-seq signal for each complex remains significant only within the two complexes, not across the complexes.

**Supplementary Figure S8.**
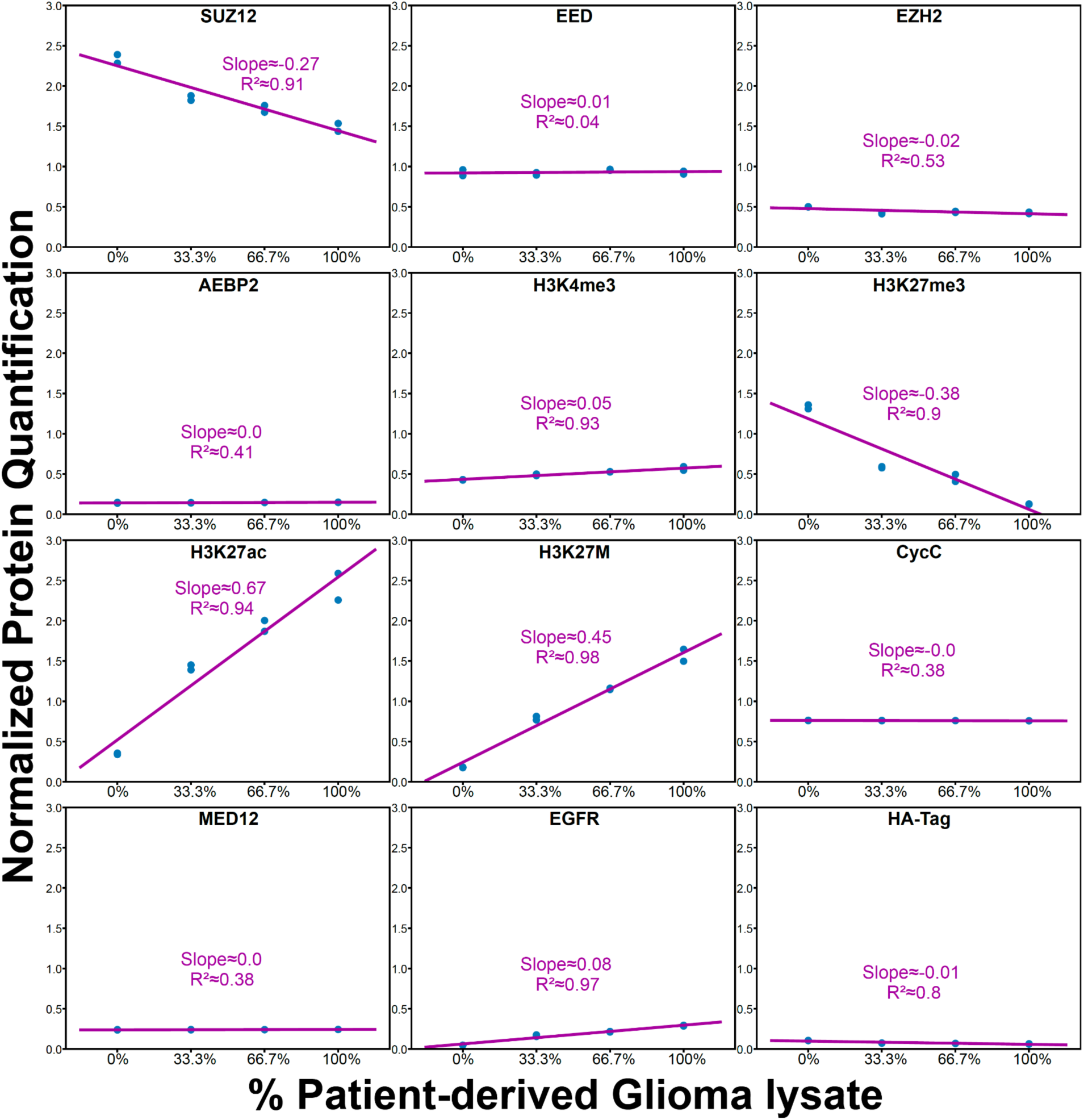
PQ-seq quantification results change linearly with respect to the cell sample mixing ratios. This figure shows the individual linear regression results for each of the proteins plotted in Figure 2C.

**Supplementary Figure S9.**
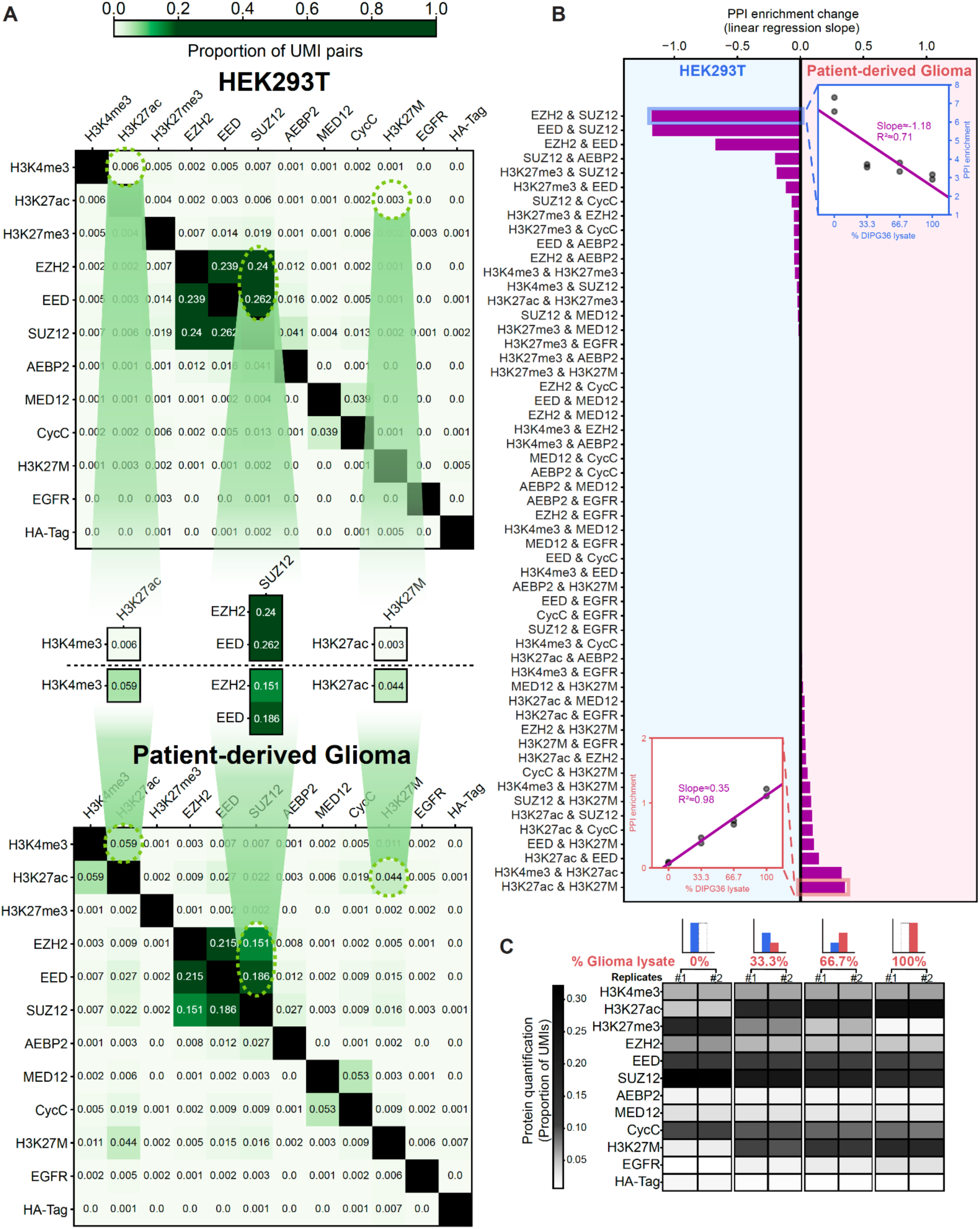
Prod-seq detects the change in PPI composition in a range of cell lysate mixtures. **(A)** Prod-seq results of the original cell lysates used. Top: Prod-seq UMI combination distribution from HEK293T cell lysate; middle: the main PPIs that differ between the two cell lines; bottom: Prod-seq UMI combination distribution from patient-derived Glioma cell lysate. **(B)** Magnitudes of change for each PPI in the range of cell lysate mixtures. The magnitude of change (horizontal axis) is the slope of the linear regression line of the PPI enrichment values of each PPI from the Prod-seq samples. **(C)** Heatmap of cell mixing protein quantification UMI counts normalized on the total number of UMIs. In each sample, the UMI count of each protein is divided by the sum of the UMI counts of all proteins in that sample as an alternative normalization method for cross-sample comparison. The resulting values are averaged among the replicates for each condition.

**Supplementary Figure S10.**
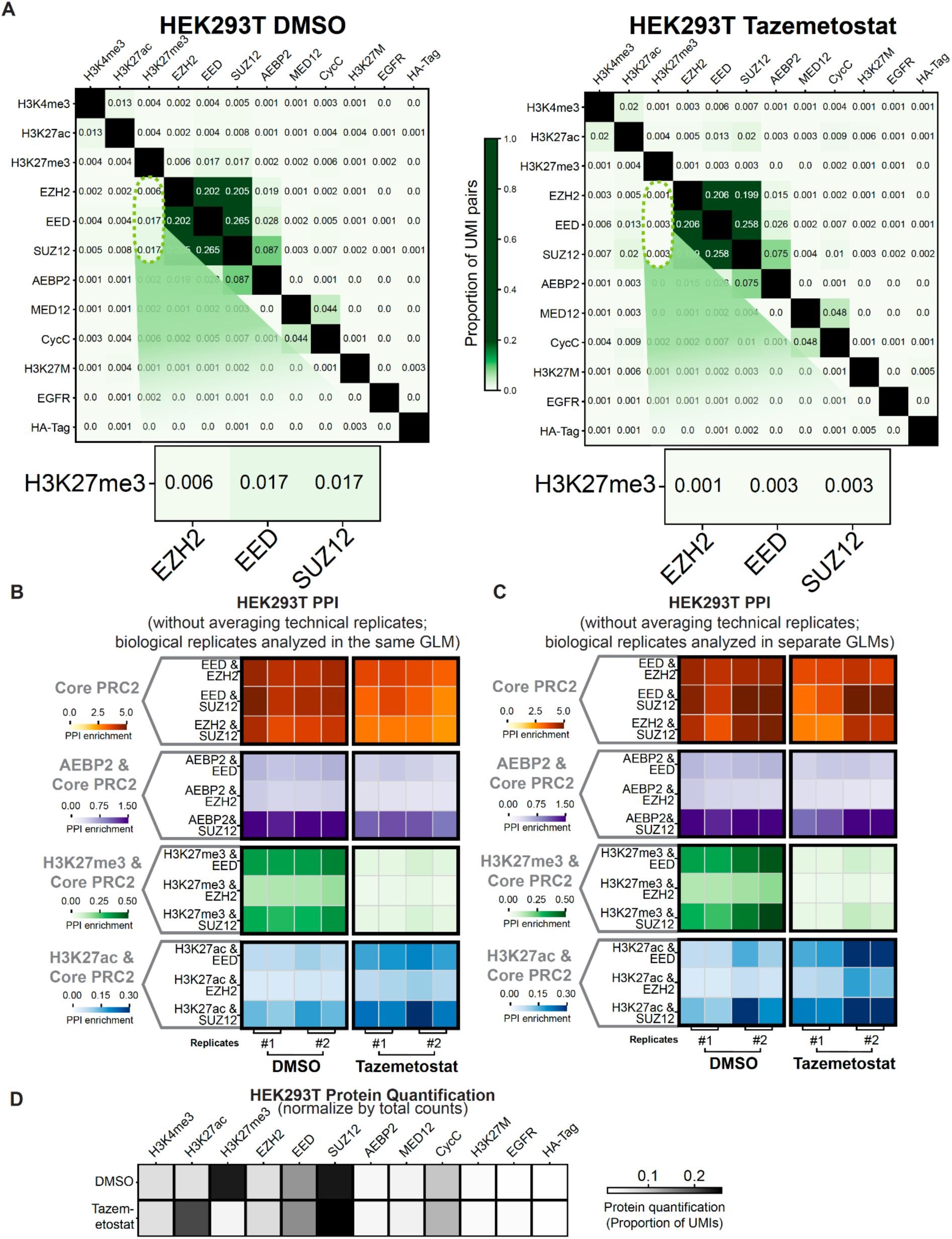
Prod-seq detects the change in PPI profiles upon Tazemetostat treatment on HEK293T cells. **(A)** Example raw UMI distribution. Bottom: zoomed-in heatmap highlighting the interaction between core PRC2 and H3K27me3. **(B)** Heatmap of the main PPIs’ enrichment values without averaging technical replicates. The PPI enrichment values are plotted in the same way as shown in main Figure 3 without averaging the values of the technical replicates. **(C)** Heatmap of PPI enrichment from separate GLM runs conducted on the biological replicates. One GLM model was run on the set of samples consisting of all biological replicate #1 samples, and another GLM model was run on the set of samples consisting of all biological replicate #2 samples. **(D)** Heatmap of protein quantification UMI counts normalized on the total number of UMIs. In each sample, the UMI count of each protein is divided by the sum of the UMI counts of all proteins in that sample as an alternative normalization method for cross-sample comparison. The resulting values are averaged among the replicates for each condition.

**Supplementary Figure S11.**
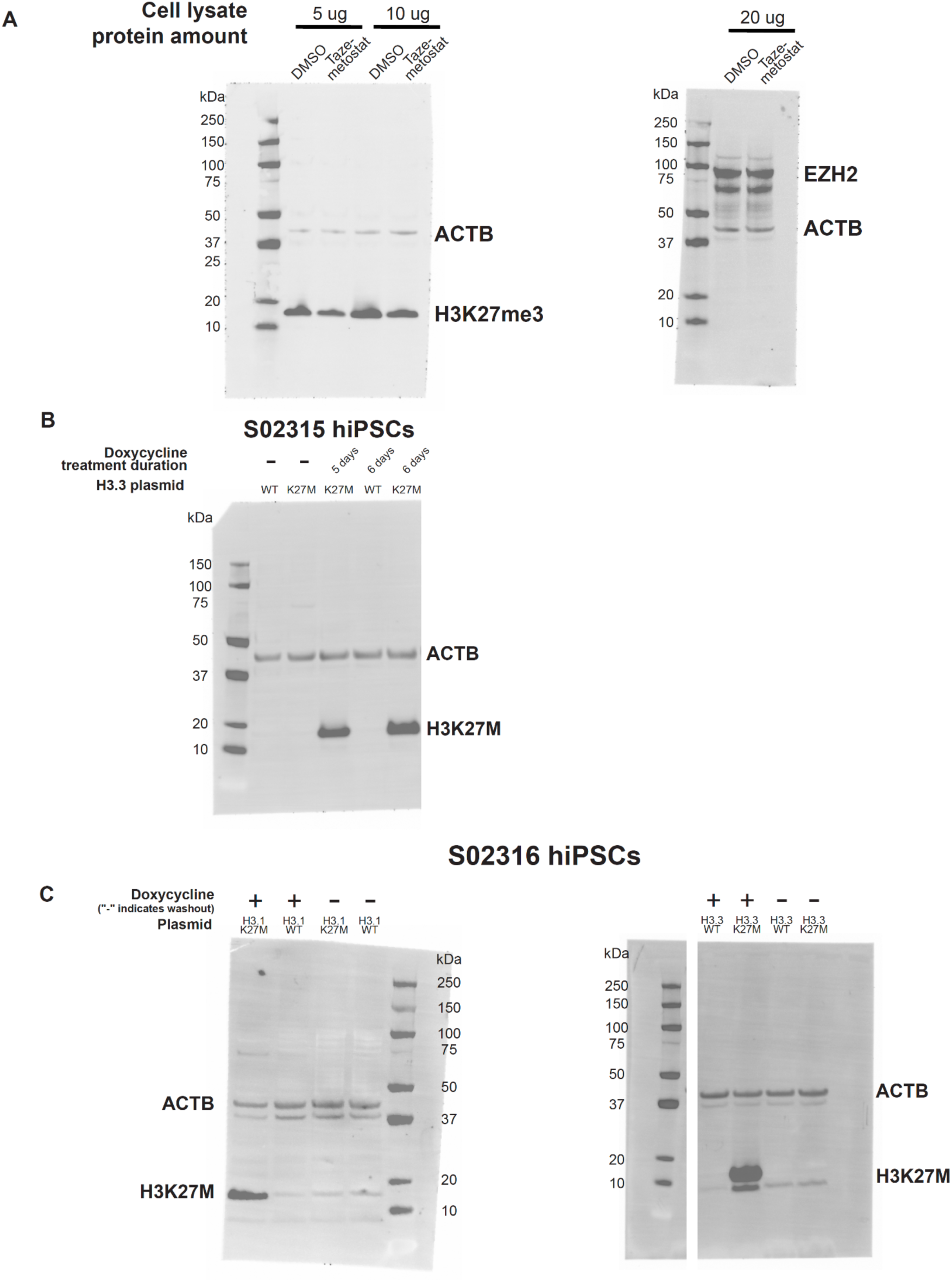
Membrane images of the immunoblot evaluations of the Tazemetostat/DMSO treatment, H3.1/3.3K27M expression and washout. **(A)** Membrane images for Tazemetostat or DMSO treatment on HEK293T cells. **(B)** Membrane image of cells expressing H3.3WT or H3.3K27M in male hiPSCs (S02315). **(C)** Membrane images for cells expressing,or following washout of, H3.1WT, H3.1K27M, H3.3WT, or H3.3K27M in female hiPSCs (S02316).

**Supplementary Figure S12.**
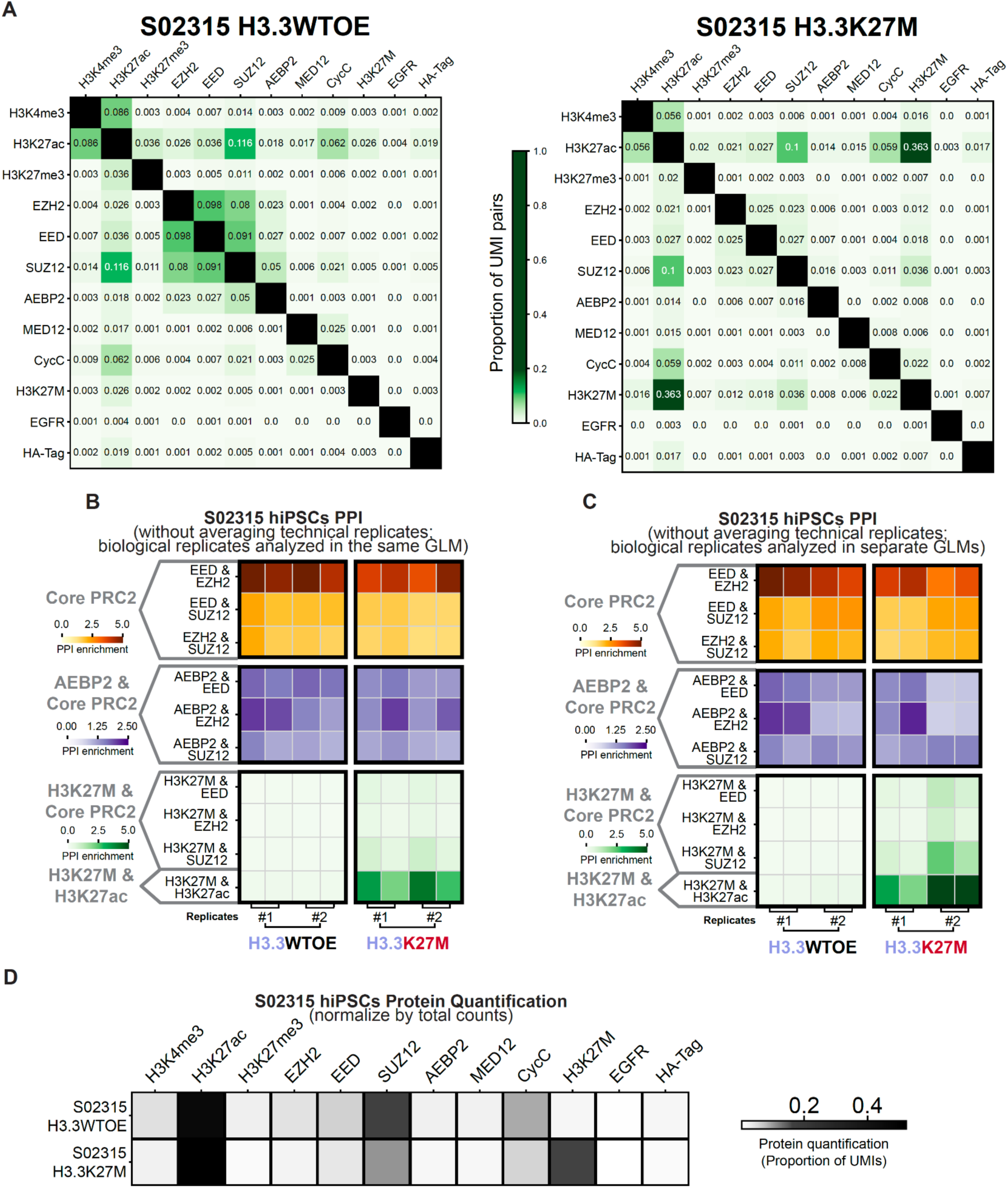
Prod-seq detects change in PPI profiles following expression of H3.3WT or H3.3K27M in male hiPSCs (S02315). **(A)** Example raw UMI distribution real data from the cells. **(B)** The PPI enrichment values are plotted in the same way as shown in main Figure 4 without averaging the values from technical replicates. **(C)** Heatmap of the enrichment values for the main PPIs calculated from the two-round GLM model run separately on biological replicates. One GLM model was run on the set of samples consisting of all biological replicate #1 samples, and another GLM model was run on the set of samples consisting of all biological replicate #2 samples. **(D)** Heatmap of protein quantification UMI counts normalized on the total number of UMIs. In each sample, the UMI count of each protein is divided by the sum of the UMI counts of all proteins in that sample as an alternative normalization method for cross-sample comparison. The resulting values are averaged among the replicates for each condition.

**Supplementary Figure S13.**
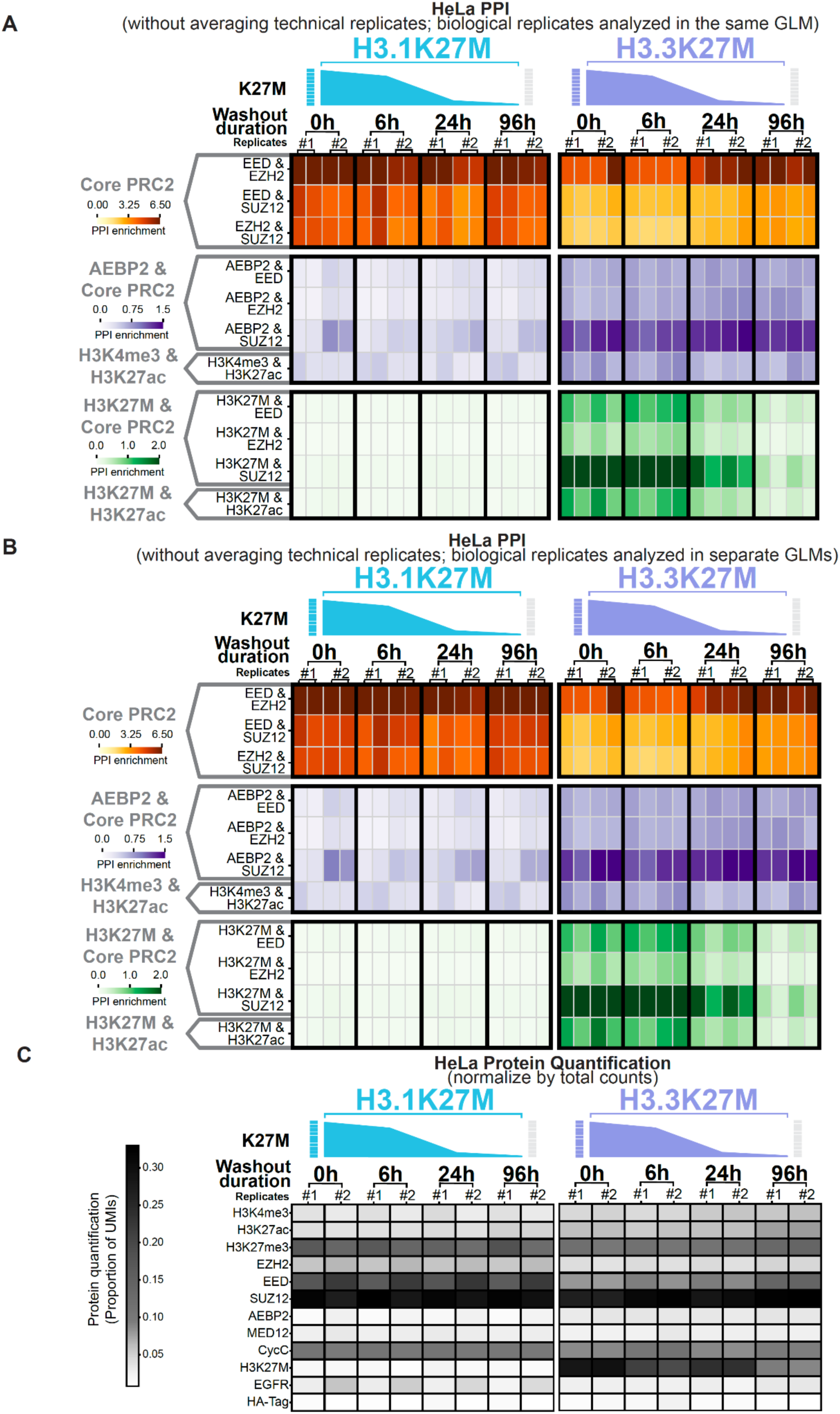
Prod-seq detects temporal changes in PPI profiles following inducible expression of H3.1K27M and H3.3K27M followed by washout timepoints in HeLa cells. **(A)** The PPI enrichment values are plotted in the same way as shown in Figure 6 without averaging the values from technical replicates. **(B)** One GLM model was run on the set of samples consisting of all biological replicate #1 samples, and another GLM model was run on the set of samples consisting of all biological replicate #2 samples. **(C)** Heatmap of protein quantification UMI counts normalized on the total number of UMIs. In each sample, the UMI count of each protein is divided by the sum of the UMI counts of all proteins in that sample as an alternative normalization method for cross-sample comparison. The resulting values are averaged among the replicates for each condition.

**Supplementary Figure S14.**
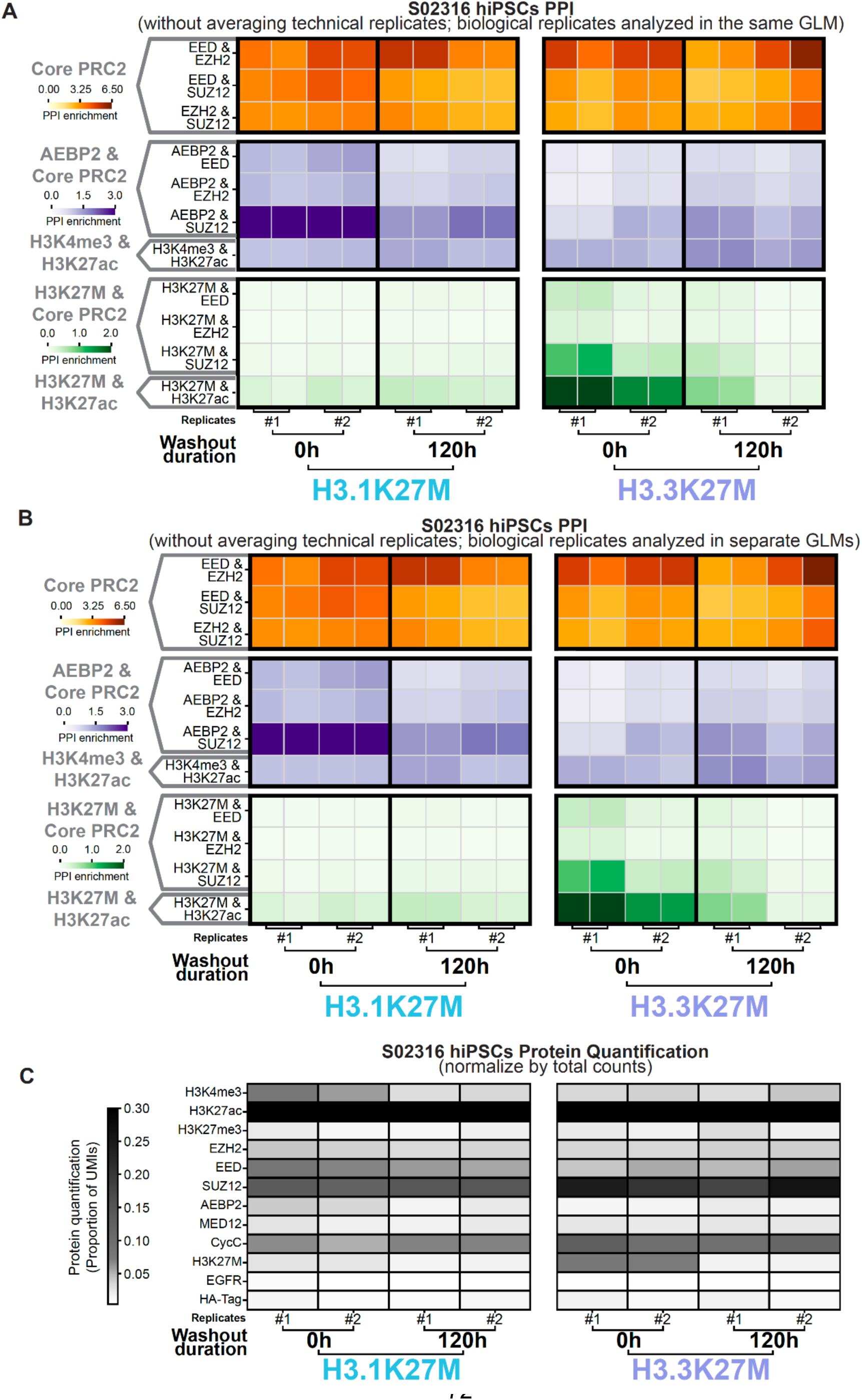
Prod-seq detects the difference in PPI profiles between H3.1K27M and H3.3K27M expression in female hiPSCs (S02316). **(A)** Heatmap of the main PPIs’ enrichment values without averaging technical replicates. The PPI enrichment values are plotted in the same way as shown in Figure 5 without averaging the values from technical replicates. **(B)** Heatmap of the main PPIs’ enrichment values calculated from the two-round GLM model run separately on biological replicates. One GLM model was run on the set of samples consisting of all biological replicate #1 samples, and another GLM model was run on the set of samples consisting of all biological replicate #2 samples. **(C)** Heatmap of protein quantification UMI counts normalized on the total number of UMIs. In each sample, the UMI count of each protein is divided by the sum of the UMI counts of all proteins in that sample as an alternative normalization method for cross-sample comparison. The resulting values are averaged among the replicates for each condition.

**Supplementary Figure S15.**
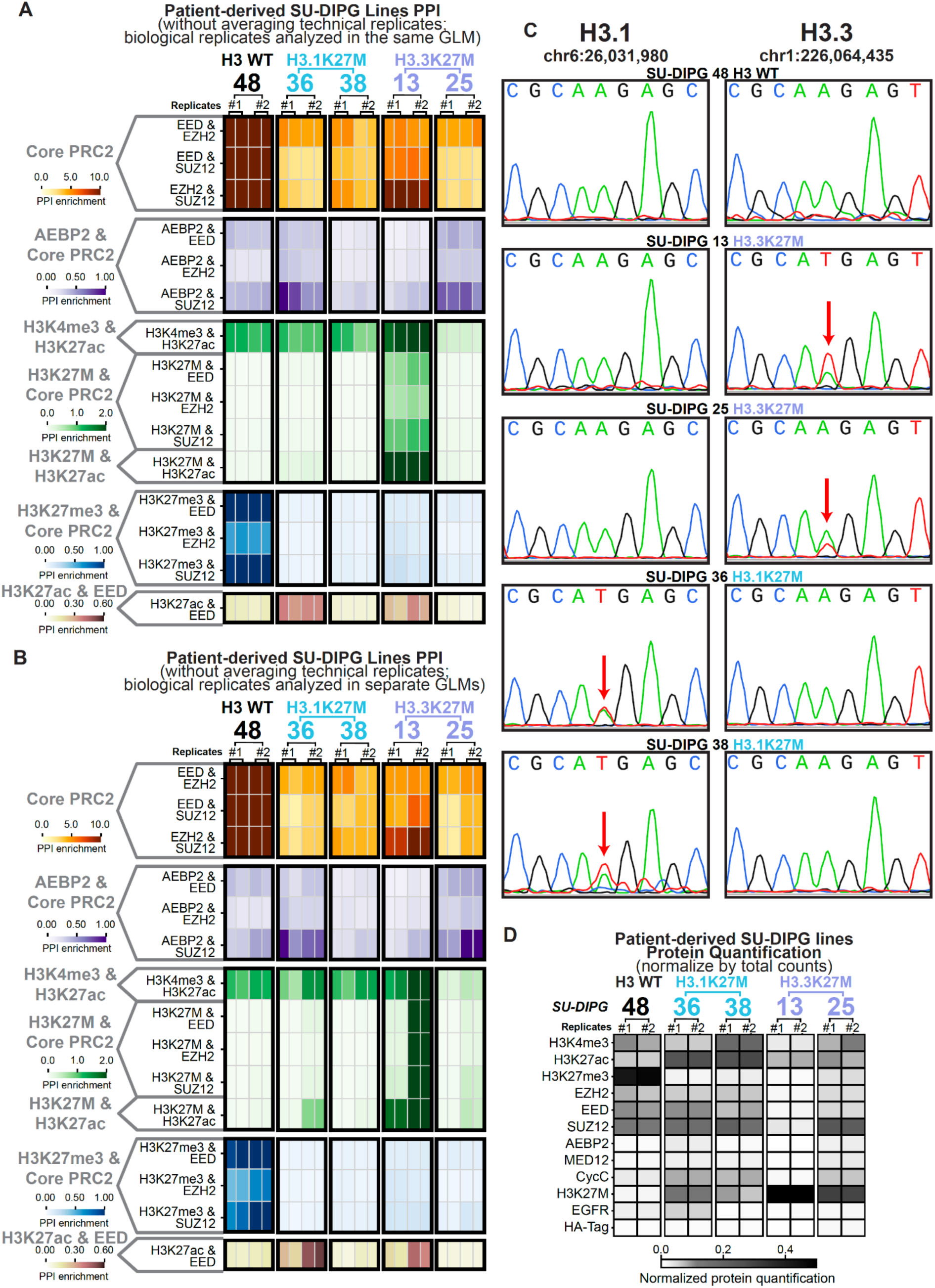
Prod-seq detects the differences in PPI profiles from patient-derived Glioma cells (SU-DIPG). **(A)** Heatmap of the main PPIs’ enrichment values without averaging technical replicates. The PPI enrichment values are plotted in the same way as shown in Figure 7 without averaging the values from technical replicates. **(B)** Heatmap of the main PPIs’ enrichment values calculated from running the two-round GLM model run separately on biological replicates. One GLM model was run on the set of samples consisting of all biological replicate #1 samples, and another GLM model was run on the set of samples consisting of all biological replicate #2 samples. **(C)** Sanger sequencing verification of the mutation status of the SU-DIPG lines. Red arrow: position of mutation. **(D)** Heatmap of protein quantification UMI counts normalized on the total number of UMIs. In each sample, each protein’s UMI count is divided by the sum of all proteins’ UMI counts in that sample as an alternative normalization method for cross-sample comparison. The resulting values are averaged among the replicates for each condition.

**Supplementary Figure S16.**
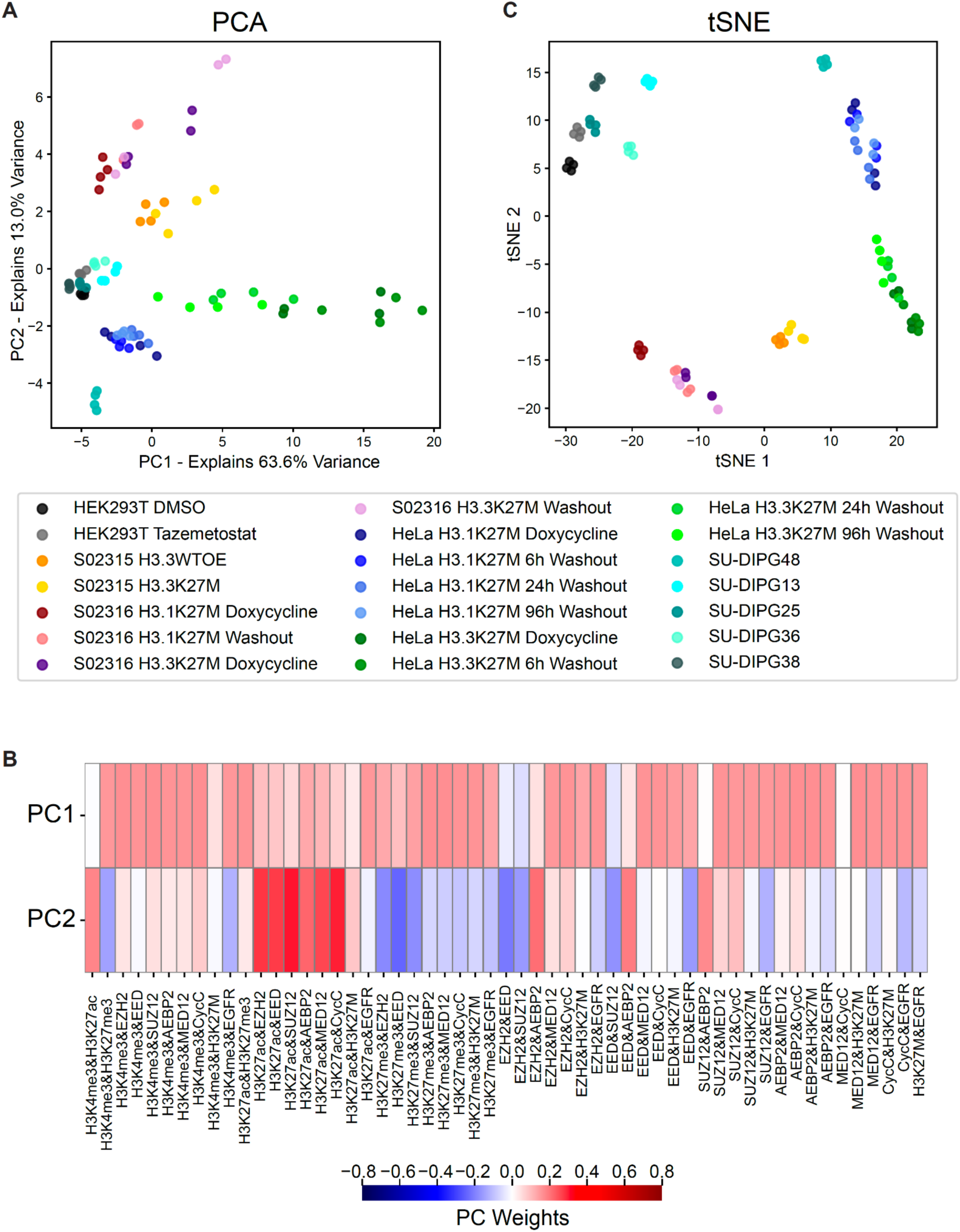
Prod-seq PPI profiles cluster by cell line. **(A)** PCA analysis of the Prod-seq PPI profiles. PCA analysis is conducted on PPI enrichment values of all the PPIs for all the cell lysate samples (for each experiment, two-round GLM was run on both biological replicates together). **(B)** The weight of each PPI for the top two PCs. **(C)** tSNE embedding of the Prod-seq PPI profiles. tSNE embedding is conducted on PPI enrichment values of all the PPIs for all the cell lysate samples (for each experiment, two-round GLM was run on both biological replicates together).

## SUPPLEMENTARY NOTES

### Note S1. Design of the DNA-caliper and ab-oligos

#### DNA-caliper arms

/5AzideN/ACACTCTTTCCCTACACGACGCTCTTCCGATCTNNNNNNNNCAGTCTGGTTGTAA GC

/5Hexynyl/GC/ideoxyU/AGCGATATCTGGAGTTCAGACGTGTGCTCTTCCGATCTNNNNNNNNCAGTCTGGTTGTAAGC

Red: Generate DNA-caliper via click chemistry
Orange: Second restriction enzyme cutting site to linearize the final product (Thermolabile USER enzyme)
Green: Library preparation primer annealing sites
Blue: Randomized bases at the beginning of sequencing reads 1 and 2 to improve sequencing quality
Purple: universal anchor sequences on the DNA-calipers to anneal with ab-oligos

#### Ab-oligo (5’ end conjugate to antibodies)

CCTTGAACCACTTCTCTANNNNNNNNNNNNNNNNNNNNNNNNNgcttacaaccagactg

Orange: First restriction enzyme cutting site for releasing the extended caliper from the original proteins (XmnI)
Green: Protein barcode (different fixed sequences of lengths 10)
Blue: UMIs (randomized bases)
Purple: annealing with anchor sequences on the DNA-calipers

All the functional and fixed sequences above except the protein barcodes are designed and empirically tested first (see the “Testing the DNA-caliper and ab-oligo designs” section below). For the protein barcodes, sequences of length 10 are selected by making sure (1) the barcode sequences combined with the fixed sequences upstream and downstream do not contain XmnI or Thermolabile USER enzyme cutting sites, and (2) avoiding overlapping or being the reverse complement of a long stretch of bases in the fixed sequences or functional sequences or other protein barcodes (to inhibit unwanted byproducts originating from undesired strand annealing).

> *Note: As an example, the criteria (2) can be achieved by running local alignments of a set of candidate 10-mers (pre-filtered for any pairwise high similarity or annealing tendency within the candidate set) against all the other fixed sequences and selecting the desired number of barcodes that result in the worst local alignment results with the fixed and functional sequences*.

The designed protein barcodes are then also empirically tested (see the “Testing the DNA-caliper and ab-oligo designs” section below) before conjugating with antibodies.

### Note S2. Purifying antibody-oligo conjugates to enrich for one oligo per antibody molecule significantly reduces the formation of “self-self” byproduct

Despite the wide use of ab-oligo conjugates in a variety of methods and protocols, most of the current applications for ab-oligos do not have strict requirements for the degree of labeling (DOL) for antibodies (i.e., the number of oligos conjugated onto the same antibody molecule). These applications typically focus on either (a) the oligos individually without mutual association/ligation/connection between oligos where all antibodies have multiple oligos almost proportionally scale up the general distribution (e.g. CITE-seq^20^) or (b) association/ligation/connection between oligos only happen when oligos are of different designs/structures so multiple oligos of the same design/structure on the same antibody would not generate byproduct with each other (e.g PLA^9^). In either of these cases, controlling for the DOL of antibodies is not a major concern and multiple oligos per Ab can in fact be beneficial by amplifying the signal.

However, in the case of Prod-seq and similar potential applications, different oligos need to be associated/ligated/connected for higher-order readout, and all oligos of different barcodes need to be designed to be the same structure to achieve the detection coverage of the full spectrum of possible target combinations. In this scenario, ab-oligo conjugates with DOL > 1 would greatly increase the amount of byproduct generated: oligos on the same antibody molecule can mutually form byproduct independent of the presence of another ab-oligo molecule or even simply resulting from ab-oligo’s unspecific binding to non-epitopes.

For example, in the case of Prod-seq, the DNA-calipers can bind to two oligos on the same antibody molecule and go through all the downstream molecular biology steps to form a standard dsDNA molecule that has no structural difference than a dsDNA product molecule resulting from true PPI detection; each byproduct molecule formed this way associates a protein barcode with itself without the need of detecting a true protein dimer or even without the need of any specific ab-oligo binding to epitopes. These byproduct molecules that we term as “self-self” can take up significant amounts of sequencing depth while providing little useful information if the DOL of the ab-oligos are not properly controlled. Moreover, controlling the DOL needs to be performed on the level of individual molecules instead of population average DOL: a population average DOL of 1 resulting from the mixture of unconjugated antibody and antibody with DOL >1 would still result in high levels of byproduct formation.

Here, we compared two sets of ab-oligos using exactly the same antibodies and oligo designs to generate conjugates, with one set enriched for a DOL of 2.5-3.8 and the other set enriched for a DOL of 1 (the set used for Prod-seq in this manuscript). Results (**Supplementary Figure S2C**) shows that: (1) self-self byproduct is strongly preferred versus true PPI detection, due to the self-self byproduct originating from any single ab-oligo molecule while true detection requires the proximity of two distinct epitope binding events; (2) enriching for DOL of 1 significantly increases the proportion of non-self-self reads, resulting in significantly higher percent of sequencing depth representing useful information. These results highlight the importance of controlling for DOL of 1 for ab-oligos used in Prod-seq or similar future methods; the DOL should be as close to purely 1 as possible as the self-self byproducts are very strongly preferred versus the true PPI products. It is also worth noting that the ab-oligos used in Prod-seq are not entirely purely DOL of 1 even though we highly enriched for DOL of 1 (**Supplementary Figure S3**), suggesting room for improvement in the future to increase the percent of meaningful sequencing depth via further improving the purification approaches and/or conjugation chemistry.

### Note S3. Testing the DNA-caliper and ab-oligo designs

This section describes the processes the current design of DNA-caliper, ab-oligo, and protein barcode sequences underwent for the validation of (1) the sequence design and the Prod-seq day 2 molecular biology steps can correctly and efficiently generate the expected product, and (2) there is no significant bias among the protein barcode sequences designed. For a new set of newly designed DNA-caliper, ab-oligos, and protein barcodes (for more details, see the section below), a similar process can be conducted before producing ab-oligos and/or conducting a full Prod-seq experiment.

In short, the testing process uses the oligo sequences in the ab-oligos to directly anneal with the DNA-calipers and proceed all the way to the final sequencing. For each protein barcode, a standard DNA oligo can be ordered with significantly reduced processing time and cost; to test the ab-oligo sequence, use 1 μL of 1 μM of the ab-oligo to dilute with 4 μL of rCutSmart working solution (Prod-seq protocol in appendix); this 5 μL of ab-oligo suspension can proceed with Prod-seq treated as if the product of step 3 of “DNA-caliper binding and strand extension” in the Prod-seq protocol. Namely, add 5 μL of incubated DNA-caliper with ETSSB and proceed with the downstream steps all the way to sequencing.

> *Note: as the ab-oligos are in suspension in this mimic Prod-seq experiment, all washing and taking supernatant steps for the original NHS beads in the Prod-seq protocol day 2 should be skipped; the speedbeads steps in the library preparation process should be proceeded as in a regular Prod-seq library preparation*.

Sequencing results and the band intensity of the Prod-seq library preparation PCR product should be able to provide an estimate of the efficiency of the sequenced design. To further confirm that no significant bias is observed in a protein barcode set, different barcodes can be mixed equimolarly to achieve 1 μL of 1 μM of the ab-oligo different barcodes mix, and then proceed with the testing process as mentioned in the paragraph above. Relative levels between the different PPIs in the sequencing result can indicate whether any protein barcode is significantly favored during the Prod-seq process.

### Note S4. Two-round GLM for the estimation of PPI background distribution

Prod-seq employs a two-round GLM approach similar to the one developed in HiC-DC^24^ for the estimation of general PPI detection background. Similar to HiC-DC, the first round of GLM is to exclude the outliers (nominal p-value < 0.025; more details below) with extremely high counts that are very likely to have high number of reads denoting true PPIs; this helps to more accurately estimate the background using the remaining values in the second GLM for increased power. Using UMI-deduplicated barcode pairwise association counts as input, the GLM process takes the readout from a group of Prod-seq samples (cell lines/conditions/perturbations to compare) for the collective estimation of the background distribution function with the assumption that the general background distribution is not highly dependent on the biological conditions and perturbations. When normalizing the input data, the UMI-deduplicated barcode pairwise association counts for the samples in the group are downscaled to the one with the least number of total UMI-deduplicated barcode pairwise association counts to account for sequencing depth differences before running the GLMs.

In both GLMs, the non-present negative control (HA-Tag)’s interaction readout with other protein targets are used as the independent variable for the estimation of all other protein’s pairwise background levels (HA-Tag itself does not get any estimation of background from the GLMs).

The background distribution is assumed to be beta-binomial distributed, namely (for protein *i* and *j* other than HA-Tag with *i* ≠ *j* in sample/condition *k*):

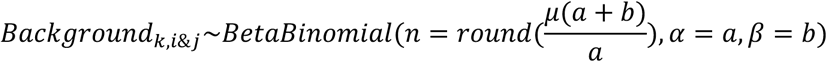

with the mean *μ* = *C*_0_ + *C*_1_(*X*_*k,i&HATag*_ + *X*_*k,j&HATag*_) being the linear combination part,

where *X*_*k,i&HATag*_ and *X*_*k,j&HATag*_ are protein *i* or *j*’s downscaled UMI-deduplicated barcode association counts with HA-Tag in sample/condition *k* respectively, and *a*, *b*, *C*_0_, *C*_1_ are parameters estimated by maximum likelihood estimation (MLE); namely, *a*, *b*, *C*_0_, *C*_1_ take the value of:

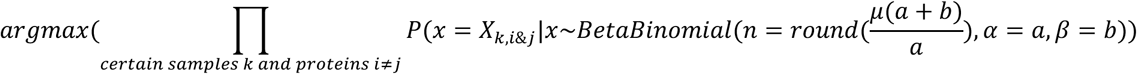

with *μ* = *C*_0_ + *C*_1_(*X*_*k,i&HATag*_ + *X*_*k,j&HATag*_) as described above and *certain samples k, proteins i* ≠ *j* depends on which round of GLM is being run with neither *i* nor *j* is HA-Tag.

The first GLM’s MLE is run on all PPI barcode association counts in all samples in the group such that *certain samples k*, *proteins i* ≠ *j* for the first GLM is all combinations of *k*, *i*, *j* that satisfies *i* ≠ *j* and neither *i* not *j* is HA-Tag; the second GLM’s MLE is rerun on all combinations of *k*, *i*, *j* included in round one GLM except the combinations that results in 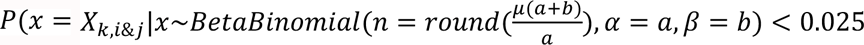 from the result of the first GLM (with the MLE estimators of *a*, *b*, *C*_0_, *C*_1_ from the first round GLM plugged into the calculation and *μ* = *C*_0_ + *C*_1_(*X*_*k,i&HATag*_ + *X*_*k,j&HATag*_)).

The resulting MLE estimators of the second GLM are used to calculate the estimated background level for each PPI in each condition as the mean of its corresponding beta-binomial distribution. The enrichment value of each PPI is then calculated as the ratio of observed counts over the estimated background level:

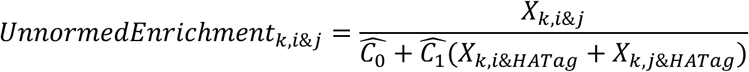

Where *X*_*k,i&j*_ is protein *i* and *j*’s downscaled UMI-deduplicated barcode association counts in each sample/condition *k*, with protein *i* ≠ *j* and neither is HA-Tag, and 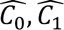 are the MLE estimators of *C*_0_, *C*_1_ from the second round of GLM respectively.

To account for any possible signal-to-noise ratio variation among the Prod-seq samples in the group, the PPI enrichment values are further normalized according to the enrichment value of the positive control PPI (MED12 & Cyclin C) in the same sample to obtain the enrichment values for each PPI for cross-sample comparison:

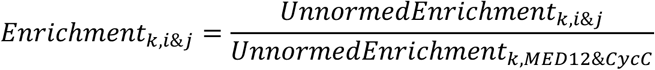

The GLM fitting process is performed using custom python script (**Code Availability**) with the MLE estimations performed using Scipy’s minimize function with the “Nelder-Mead” solver. The round 1 GLM parameters are initialized as 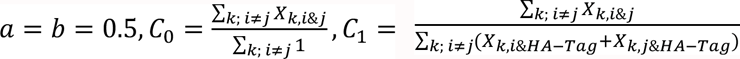 and the round 2 GLM takes the MLE estimators of round 1 GLM as the initial guesses. Round one GLM is fitted for a maximum of 500 iterations of the minimize function and round two GLM is fitted for a maximum of 1000 iterations of the minimize function.

## SUPPLEMENTARY TABLES

**Supplementary Table 1. Per-sample cost calculations for Prod&PQ-seq.**

Provided as an xlsx file.

**Supplementary Table 2.**
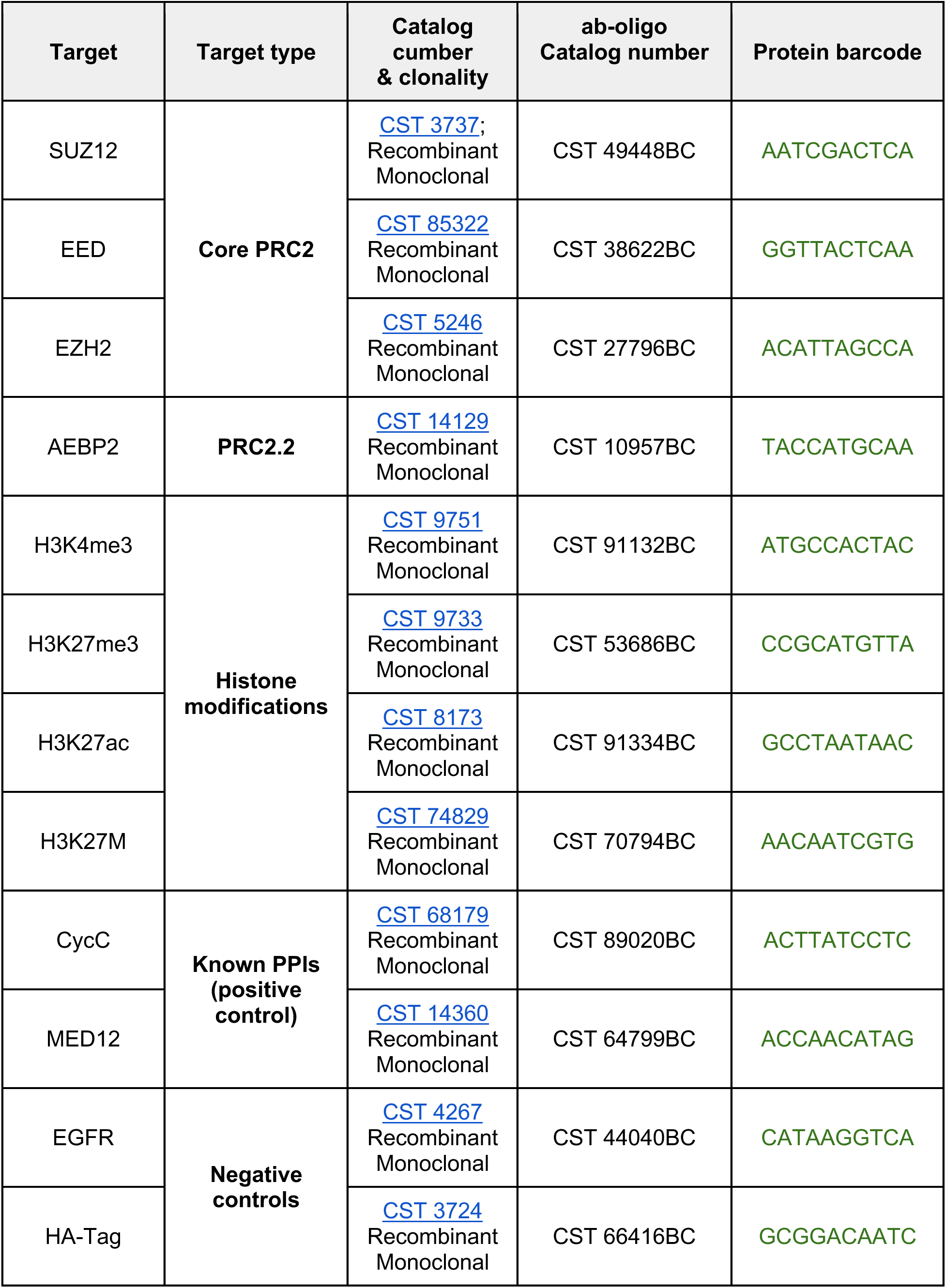
Antibody-oligo information. Note: the full sequences of the oligonucleotides conjugated for each ab-oligo are the of following format with the green section replaced by the protein barcode sequences in this table (more in-depth descriptions is provided in **Supplementary Note S1**): CCTTGAACCACTTCTCTANNNNNNNNNNNNNNNNNNNNNNNNNgcttacaaccagactg

| Target | Target type | Catalog number & clonality | ab-oligo Catalog number | Protein barcode |
| --- | --- | --- | --- | --- |
| SUZ12 | **Core PRC2** | CST 3737;<br>Recombinant Monoclonal | CST 49448BC | AATCGACTCA |
| EED |  | CST 85322<br>Recombinant Monoclonal | CST 38622BC | GGTTACTCAA |
| EZH2 |  | CST 5246<br>Recombinant Monoclonal | CST 27796BC | ACATTAGCCA |
| AEBP2 | **PRC2.2** | CST 14129<br>Recombinant Monoclonal | CST 10957BC | TACCATGCAA |
| H3K4me3 | **Histone modifications** | CST 9751<br>Recombinant Monoclonal | CST 91132BC | ATGCCACTAC |
| H3K27me3 |  | CST 9733<br>Recombinant Monoclonal | CST 53686BC | CCGCATGTTA |
| H3K27ac |  | CST 8173<br>Recombinant Monoclonal | CST 91334BC | GCCTAATAAC |
| H3K27M |  | CST 74829<br>Recombinant Monoclonal | CST 70794BC | AACAATCGTG |
| CycC | **Known PPIs (positive control)** | CST 68179<br>Recombinant Monoclonal | CST 89020BC | ACTTATCCTC |
| MED12 |  | CST 14360<br>Recombinant Monoclonal | CST 64799BC | ACCAACATAG |
| EGFR | **Negative controls** | CST 4267<br>Recombinant Monoclonal | CST 44040BC | CATAAGGTCA |
| HA-Tag |  | CST 3724<br>Recombinant Monoclonal | CST 66416BC | GCGGACAATC |

**Supplementary Table 3.**
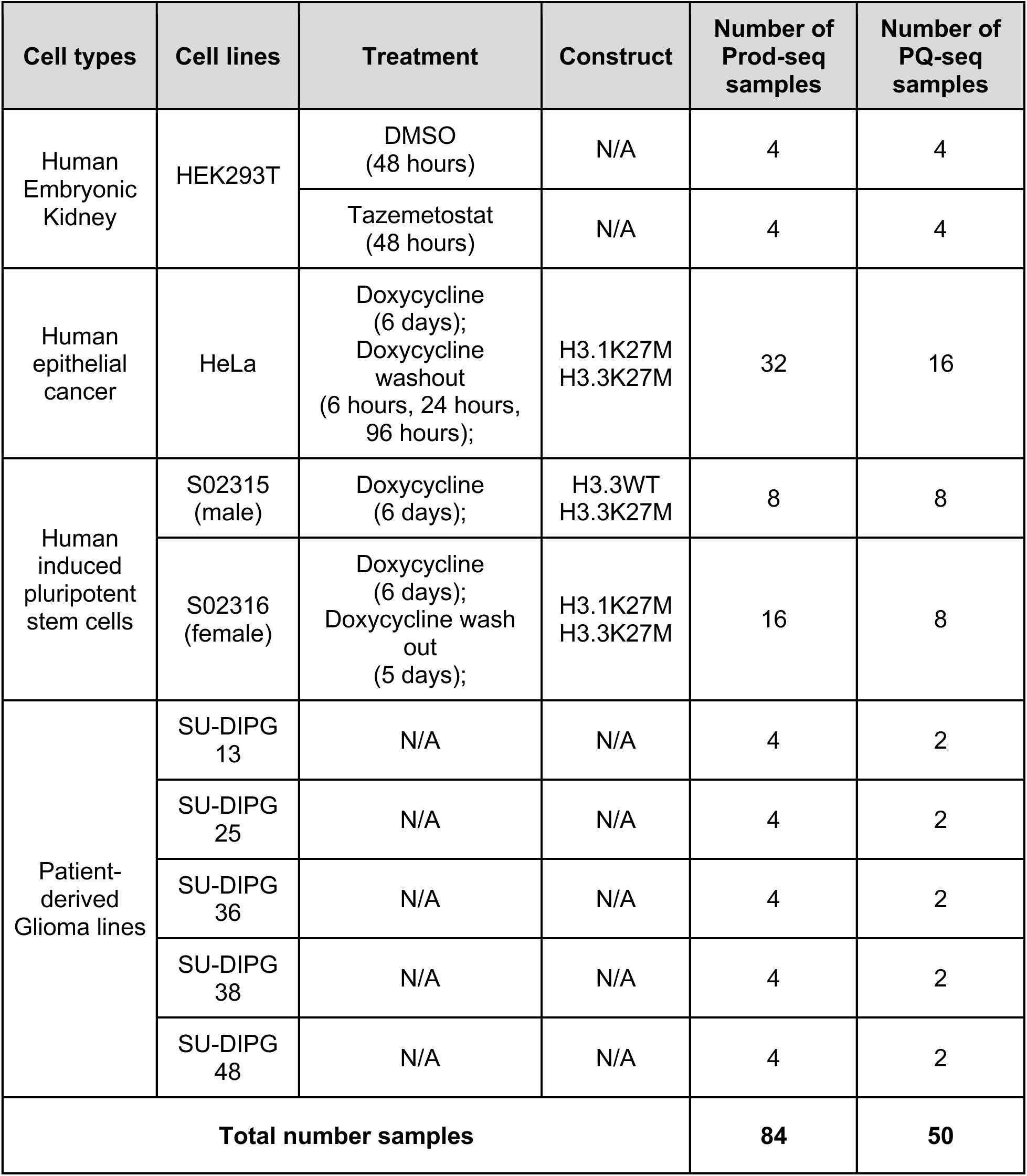
List of cell samples Prod-seq and PQ-seq tested in this study.

| Cell types | Cell lines | Treatment | Construct | Number of Prod-seq samples | Number of PQ-seq samples |
| --- | --- | --- | --- | --- | --- |
| Human Embryonic Kidney | HEK293T | DMSO (48 hours) | N/A | 4 | 4 |
|  |  | Tazemetostat (48 hours) | N/A | 4 | 4 |
| Human epithelial cancer | HeLa | Doxycycline (6 days);<br>Doxycycline washout (6 hours, 24 hours, 96 hours); | H3.1K27M<br>H3.3K27M | 32 | 16 |
| Human induced pluripotent stem cells | S02315 (male) | Doxycycline (6 days); | H3.3WT<br>H3.3K27M | 8 | 8 |
|  | S02316 (female) | Doxycycline (6 days);<br>Doxycycline wash out (5 days); | H3.1K27M<br>H3.3K27M | 16 | 8 |
| Patient-derived Glioma lines | SU-DIPG 13 | N/A | N/A | 4 | 2 |
|  | SU-DIPG 25 | N/A | N/A | 4 | 2 |
|  | SU-DIPG 36 | N/A | N/A | 4 | 2 |
|  | SU-DIPG 38 | N/A | N/A | 4 | 2 |
|  | SU-DIPG 48 | N/A | N/A | 4 | 2 |
| **Total number samples** |  |  |  | **84** | **50** |

## REFERENCES

1. Silveira, M.A.D., and Bilodeau, S. (2018). Defining the Transcriptional Ecosystem. Mol. Cell 72, 920–924. 10.1016/j.molcel.2018.11.022.

2. Wright, P.E., and Dyson, H.J. (2015). Intrinsically disordered proteins in cellular signalling and regulation. Nat. Rev. Mol. Cell Biol. 16, 18–29. 10.1038/nrm3920.

3. Berger, S.L. (2007). The complex language of chromatin regulation during transcription. Nature 447, 407–412. 10.1038/nature05915.

4. Pawson, T. (2007). Dynamic control of signaling by modular adaptor proteins. Curr. Opin. Cell Biol. 19, 112–116. 10.1016/j.ceb.2007.02.013.

5. Singh, A. (2024). Understanding protein interaction dynamics. Nat. Methods 21, 2226–2227. 10.1038/s41592-024-02545-7.

6. Brückner, A., Polge, C., Lentze, N., Auerbach, D., and Schlattner, U. (2009). Yeast Two-Hybrid, a Powerful Tool for Systems Biology. Int. J. Mol. Sci. 10, 2763–2788. 10.3390/ijms10062763.

7. Richards, A.L., Eckhardt, M., and Krogan, N.J. (2021). Mass spectrometry-based protein–protein interaction networks for the study of human diseases. Mol. Syst. Biol. 17, e8792. 10.15252/msb.20188792.

8. Yu, C., and Huang, L. (2018). Cross-Linking Mass Spectrometry (XL-MS): an Emerging Technology for Interactomics and Structural Biology. Anal. Chem. 90, 144–165. 10.1021/acs.analchem.7b04431.

9. Hegazy, M., Cohen-Barak, E., Koetsier, J.L., Najor, N.A., Arvanitis, C., Sprecher, E., Green, K.J., and Godsel, L.M. (2020). Proximity Ligation Assay for Detecting Protein-Protein Interactions and Protein Modifications in Cells and Tissues In Situ. Curr. Protoc. Cell Biol. 89, e115. 10.1002/cpcb.115.

10. Laman Trip, D.S., van Oostrum, M., Memon, D., Frommelt, F., Baptista, D., Panneerselvam, K., Bradley, G., Licata, L., Hermjakob, H., Orchard, S., et al. (2025). A tissue-specific atlas of protein–protein associations enables prioritization of candidate disease genes. Nat. Biotechnol., 1–14. 10.1038/s41587-025-02659-z.

11. Fischer, S.N., Claussen, E.R., Kourtis, S., Sdelci, S., Orchard, S., Hermjakob, H., Kustatscher, G., and Drew, K. (2025). hu.MAP3.0: atlas of human protein complexes by integration of >25,000 proteomic experiments. Mol. Syst. Biol. 21, 911–943. 10.1038/s44320-025-00121-5.

12. Oughtred, R., Rust, J., Chang, C., Breitkreutz, B.-J., Stark, C., Willems, A., Boucher, L., Leung, G., Kolas, N., Zhang, F., et al. (2021). The BioGRID database: A comprehensive biomedical resource of curated protein, genetic, and chemical interactions. Protein Sci. Publ. Protein Soc. 30, 187–200. 10.1002/pro.3978.

13. Scott, D.E., Bayly, A.R., Abell, C., and Skidmore, J. (2016). Small molecules, big targets: drug discovery faces the protein-protein interaction challenge. Nat. Rev. Drug Discov. 15, 533–550. 10.1038/nrd.2016.29.

14. Arrowsmith, C.H., Bountra, C., Fish, P.V., Lee, K., and Schapira, M. (2012). Epigenetic protein families: a new frontier for drug discovery. Nat. Rev. Drug Discov. 11, 384–400. 10.1038/nrd3674.

15. Vistain, L., Van Phan, H., Keisham, B., Jordi, C., Chen, M., Reddy, S.T., and Tay, S. (2022). Quantification of extracellular proteins, protein complexes and mRNAs in single cells by proximity sequencing. Nat. Methods 19, 1578–1589. 10.1038/s41592-022-01684-z.

16. Johnson, K.L., Qi, Z., Yan, Z., Wen, X., Nguyen, T.C., Zaleta-Rivera, K., Chen, C.-J., Fan, X., Sriram, K., Wan, X., et al. (2021). Revealing protein-protein interactions at the transcriptome scale by sequencing. Mol. Cell 81, 4091–4103.e9. 10.1016/j.molcel.2021.07.006.

17. Liu, Y., Sundah, N.R., Ho, N.R.Y., Shen, W.X., Xu, Y., Natalia, A., Yu, Z., Seet, J.E., Chan, C.W., Loh, T.P., et al. (2024). Bidirectional linkage of DNA barcodes for the multiplexed mapping of higher-order protein interactions in cells. Nat. Biomed. Eng. 8, 909–923. 10.1038/s41551-024-01225-3.

18. Kivioja, T., Vähärautio, A., Karlsson, K., Bonke, M., Enge, M., Linnarsson, S., and Taipale, J. (2012). Counting absolute numbers of molecules using unique molecular identifiers. Nat. Methods 9, 72–74. 10.1038/nmeth.1778.

19. Tornøe, C.W., Christensen, C., and Meldal, M. (2002). Peptidotriazoles on solid phase: [1,2,3]-triazoles by regiospecific copper(i)-catalyzed 1,3-dipolar cycloadditions of terminal alkynes to azides. J. Org. Chem. 67, 3057–3064. 10.1021/jo011148j.

20. Stoeckius, M., Hafemeister, C., Stephenson, W., Houck-Loomis, B., Chattopadhyay, P.K., Swerdlow, H., Satija, R., and Smibert, P. (2017). Simultaneous epitope and transcriptome measurement in single cells. Nat. Methods 14, 865–868. 10.1038/nmeth.4380.

21. Healy, E., Mucha, M., Glancy, E., Fitzpatrick, D.J., Conway, E., Neikes, H.K., Monger, C., Mierlo, G.V., Baltissen, M.P., Koseki, Y., et al. (2019). PRC2.1 and PRC2.2 Synergize to Coordinate H3K27 Trimethylation. Mol. Cell 76, 437–452.e6. 10.1016/j.molcel.2019.08.012.

22. Hauri, S., Comoglio, F., Seimiya, M., Gerstung, M., Glatter, T., Hansen, K., Aebersold, R., Paro, R., Gstaiger, M., and Beisel, C. (2016). A High-Density Map for Navigating the Human Polycomb Complexome. Cell Rep. 17, 583–595. 10.1016/j.celrep.2016.08.096.

23. Turunen, M., Spaeth, J.M., Keskitalo, S., Park, M.J., Kivioja, T., Clark, A.D., Mäkinen, N., Gao, F., Palin, K., Nurkkala, H., et al. (2014). Uterine leiomyoma-linked MED12 mutations disrupt Mediator-associated CDK activity. Cell Rep. 7, 654–660. 10.1016/j.celrep.2014.03.047.

24. Carty, M., Zamparo, L., Sahin, M., González, A., Pelossof, R., Elemento, O., and Leslie, C.S. (2017). An integrated model for detecting significant chromatin interactions from high-resolution Hi-C data. Nat. Commun. 8, 15454. 10.1038/ncomms15454.

25. Hoy, S.M. (2020). Tazemetostat: First Approval. Drugs 80, 513–521. 10.1007/s40265-020-01288-x.

26. Julia, E., and Salles, G. EZH2 inhibition by tazemetostat: mechanisms of action, safety and efficacy in relapsed/refractory follicular lymphoma. Future Oncol. 17, 2127–2140. 10.2217/fon-2020-1244.

27. Tao, J., Alessandri, L., Gasparetto, A., Zhao, L., Zhang, X., Alt, F.W., and Chiarle, R. (2025). Epigenetic changes by EZH2 inhibition increase translocations in B cells with high AID activity or DNA repair deficiency. Blood 146, 2203–2216. 10.1182/blood.2024026131.

28. Panopoulos, A.D., D’Antonio, M., Benaglio, P., Williams, R., Hashem, S.I., Schuldt, B.M., DeBoever, C., Arias, A.D., Garcia, M., Nelson, B.C., et al. (2017). iPSCORE: A Resource of 222 iPSC Lines Enabling Functional Characterization of Genetic Variation across a Variety of Cell Types. Stem Cell Rep. 8, 1086–1100. 10.1016/j.stemcr.2017.03.012.

29. Furth, N., Algranati, D., Dassa, B., Beresh, O., Fedyuk, V., Morris, N., Kasper, L.H., Jones, D., Monje, M., Baker, S.J., et al. (2022). H3-K27M-mutant nucleosomes interact with MLL1 to shape the glioma epigenetic landscape. Cell Rep. 39, 110836. 10.1016/j.celrep.2022.110836.

30. Piunti, A., Hashizume, R., Morgan, M.A., Bartom, E.T., Horbinski, C.M., Marshall, S.A., Rendleman, E.J., Ma, Q., Takahashi, Y., Woodfin, A.R., et al. (2017). Therapeutic targeting of polycomb and BET bromodomain proteins in diffuse intrinsic pontine gliomas. Nat. Med. 23, 493–500. 10.1038/nm.4296.

31. Nagaraja, S., Quezada, M.A., Gillespie, S.M., Arzt, M., Lennon, J.J., Woo, P.J., Hovestadt, V., Kambhampati, M., Filbin, M.G., Suva, M.L., et al. (2019). Histone variant and cell context determine H3K27M reprogramming of the enhancer landscape and oncogenic state. Mol. Cell 76, 965–980.e12. 10.1016/j.molcel.2019.08.030.

32. Cao, Y., Patel, L., Alcoser, L., Mendenhall, E., Benner, C., Heinz, S., and Goren, A. (2025). Automated chromatin profiling with spa-ChIP-seq uncovers the impacts of condition variations. Genome Res. 10.1101/gr.281320.125.

33. El-Hashash, A.H.K. (2021). Histone H3K27M Mutation in Brain Tumors. In Histone Mutations and Cancer, D. Fang and J. Han, eds. (Springer), pp. 43–52. 10.1007/978-981-15-8104-5_3.

34. Lewis, P.W., Müller, M.M., Koletsky, M.S., Cordero, F., Lin, S., Banaszynski, L.A., Garcia, B.A., Muir, T.W., Becher, O.J., and Allis, C.D. (2013). Inhibition of PRC2 activity by a gain-of-function H3 mutation found in pediatric glioblastoma. Science 340, 857–861. 10.1126/science.1232245.

35. Bender, S., Tang, Y., Lindroth, A.M., Hovestadt, V., Jones, D.T.W., Kool, M., Zapatka, M., Northcott, P.A., Sturm, D., Wang, W., et al. (2013). Reduced H3K27me3 and DNA hypomethylation are major drivers of gene expression in K27M mutant pediatric high-grade gliomas. Cancer Cell 24, 660–672. 10.1016/j.ccr.2013.10.006.

36. Harutyunyan, A.S., Krug, B., Chen, H., Papillon-Cavanagh, S., Zeinieh, M., De Jay, N., Deshmukh, S., Chen, C.C.L., Belle, J., Mikael, L.G., et al. (2019). H3K27M induces defective chromatin spread of PRC2-mediated repressive H3K27me2/me3 and is essential for glioma tumorigenesis. Nat. Commun. 10, 1262. 10.1038/s41467-019-09140-x.

37. Lewis, P.W., Müller, M.M., Koletsky, M.S., Cordero, F., Lin, S., Banaszynski, L.A., Garcia, B.A., Muir, T.W., Becher, O.J., and Allis, C.D. (2013). Inhibition of PRC2 Activity by a Gain-of-Function H3 Mutation Found in Pediatric Glioblastoma. Science 340, 857–861. 10.1126/science.1232245.

38. Nagaraja, S., Vitanza, N.A., Woo, P.J., Taylor, K.R., Liu, F., Zhang, L., Li, M., Meng, W., Ponnuswami, A., Sun, W., et al. (2017). Transcriptional Dependencies in Diffuse Intrinsic Pontine Glioma. Cancer Cell 31, 635–652.e6. 10.1016/j.ccell.2017.03.011.

39. Sun, B.B., Chiou, J., Traylor, M., Benner, C., Hsu, Y.-H., Richardson, T.G., Surendran, P., Mahajan, A., Robins, C., Vasquez-Grinnell, S.G., et al. (2023). Plasma proteomic associations with genetics and health in the UK Biobank. Nature 622, 329–338. 10.1038/s41586-023-06592-6.

40. Patel, L., Cao, Y., Xu, T., Modolo, E., Dishon, T., Zhang, L., Mendenhall, E., Heinz, S., Simon, I., Benner, C., et al. (2025). Improved spike-in normalization clarifies the relationship between active histone modifications and transcription. Preprint at bioRxiv, 10.1101/2025.11.25.690627 https://doi.org/10.1101/2025.11.25.690627.

41. Chen, S., Zhou, Y., Chen, Y., and Gu, J. (2018). fastp: an ultra-fast all-in-one FASTQ preprocessor. Bioinformatics 34, i884–i890. 10.1093/bioinformatics/bty560.

42. Li, H., and Durbin, R. (2009). Fast and accurate short read alignment with Burrows-Wheeler transform. Bioinforma. Oxf. Engl. 25, 1754–1760. 10.1093/bioinformatics/btp324.

43. Li, H., Handsaker, B., Wysoker, A., Fennell, T., Ruan, J., Homer, N., Marth, G., Abecasis, G., Durbin, R., and 1000 Genome Project Data Processing Subgroup (2009). The Sequence Alignment/Map format and SAMtools. Bioinforma. Oxf. Engl. 25, 2078–2079. 10.1093/bioinformatics/btp352.

44. Jun, G., Wing, M.K., Abecasis, G.R., and Kang, H.M. (2015). An efficient and scalable analysis framework for variant extraction and refinement from population scale DNA sequence data. Genome Res., gr.176552.114. 10.1101/gr.176552.114.

45. Heinz, S., Benner, C., Spann, N., Bertolino, E., Lin, Y.C., Laslo, P., Cheng, J.X., Murre, C., Singh, H., and Glass, C.K. (2010). Simple combinations of lineage-determining transcription factors prime cis-regulatory elements required for macrophage and B cell identities. Mol. Cell 38, 576–589. 10.1016/j.molcel.2010.05.004.

46. Ramírez, F., Ryan, D.P., Grüning, B., Bhardwaj, V., Kilpert, F., Richter, A.S., Heyne, S., Dündar, F., and Manke, T. (2016). deepTools2: a next generation web server for deep-sequencing data analysis. Nucleic Acids Res. 44, W160–W165. 10.1093/nar/gkw257.

47. Robinson, J.T., Thorvaldsdóttir, H., Winckler, W., Guttman, M., Lander, E.S., Getz, G., and Mesirov, J.P. (2011). Integrative genomics viewer. Nat. Biotechnol. 29, 24–26. 10.1038/nbt.1754.

48. Quinlan, A.R., and Hall, I.M. (2010). BEDTools: a flexible suite of utilities for comparing genomic features. Bioinformatics 26, 841–842. 10.1093/bioinformatics/btq033.

49. Sanson, K.R., Hanna, R.E., Hegde, M., Donovan, K.F., Strand, C., Sullender, M.E., Vaimberg, E.W., Goodale, A., Root, D.E., Piccioni, F., et al. (2018). Optimized libraries for CRISPR-Cas9 genetic screens with multiple modalities. Nat. Commun. 9, 5416. 10.1038/s41467-018-07901-8.

